# Selective killing of cancer cells via simultaneous destabilization of mitochondrial iron metabolism and induction of ferroptosis elicited by mitochondrially targeted iron chelator deferasirox

**DOI:** 10.1101/2024.01.17.575692

**Authors:** Sukanya B Jadhav, Cristian Sandoval-Acuña, Yaiza Pacior, Petra Potomova, Kristyna Klanicova, Kristyna Blazkova, Radislav Sedlacek, Jan Stursa, Lukas Werner, Jaroslav Truksa

## Abstract

In principle, two separate ways to target cancer cells exist, the first one using iron deprivation and subsequent dysfunction of iron-dependent enzymes, and the second one inducing iron overload that leads to cell death known as ferroptosis. In this study, we introduce a compound that uniquely employs both pathways at once - mitochondrially targeted deferasirox (mitoDFX). MitoDFX deprives cells of biologically active iron while simultaneously depleting the primary cellular antioxidant glutathione (GSH) and inducing lipid peroxidation, both of which are hallmarks of ferroptosis. The role of the GSH is further supported by enhanced cell death in glutathione peroxidase 4 KO cells (GPX4 KO) induced by mitoDFX. This response is further exacerbated by simultaneous inhibition of glutathione metabolism or the pentose phosphate pathway. MitoDFX strongly affects the structure and function of mitochondria, leading to its fragmentation, ROS production and induction of mitophagy. Moreover, we found that mitoDFX treatment markedly reduces mitochondrial translation and downregulates levels of enzymatic subunits coupled with the ETC/TCA cycle as well as DNA replication and transcription, which could explain its strong cytostatic effects. In summary, mitoDFX achieves high activity and selectivity against cancer cells that seems unattainable for the conventional untargeted iron chelators, which have adverse effects on systemic iron metabolism. Our novel, targeted chelator has also shown enhanced efficacy in animal models, surpassing other mitochondrially targeted drugs and making mitoDFX a promising candidate for the development of future selective and highly effective cancer therapies.

## Introduction

Iron is a critical trace element involved in many biological processes, including oxygen transport, metabolism, cell cycle progression, and DNA synthesis and repair.^1-6^ Furthermore, iron is a key component of enzyme cofactors, heme, and iron-sulfur [Fe-S] clusters, which are partially synthesized in mitochondria. ^4,7-10^ Despite its biological importance, excessive iron levels can lead to the production of reactive oxygen species (ROS) through Fenton and Haber-Weiss reactions.^10,11^ These ROS can cause lipid peroxidation, leading to a specific form of cell death known as ferroptosis.^12,13^

Cancers exhibit rapid proliferation and altered metabolic needs, including high iron demand.^11,14-16^ Therefore, targeting iron metabolism has emerged as a promising therapeutic strategy for cancer.^14,17,18^ The iron chelator deferasirox (DFX) has shown anti-proliferative effects on several cancer subtypes^19-22^ by inhibiting DNA replication and inducing G_0_ /G_1_ cell cycle arrest.^23^ However, existing chelators lack cancer cell specificity and rely on the higher sensitivity of these cells to iron deprivation and the leaky vasculature of tumors, which results in an enhanced permeability and retention effect. Recently, mitochondrial targeting of anti-cancer compounds *via* delocalized lipophilic cations has proven to be an effective and more specific strategy for delivering experimental anti-cancer drugs.^24-26^

Here, we show that mitochondrially targeted deferasirox (mitoDFX) potently induces cell death in cancer cells at nanomolar concentrations and exerts unique effects for an iron chelator.

## Materials and Methods

### Cell line sources and culture conditions

All cancer cells and non-malignant (BJ) cells were obtained from the American Type Culture Collection (ATCC). HFP-1 human fibroblasts were a gift from Dr. Smetana (Institute of Anatomy, Charles University, Prague, Czech Republic). Cells were cultured in a humid incubator at 37 °C with 5 % CO_2_ in DMEM (MERCK) supplemented with 10 % fetal bovine serum, 100 U/ml streptomycin/penicillin (MERCK), and 2 mM L-glutamine (PAN Biotech). Cultured cells were authenticated by the STR analysis (Generi Biotech), regularly tested for mycoplasma contamination and used within three months of thawing.

### mitoDFX synthesis

Dimethylformamide (DMF) (3 ml), was added to a round bottom flask containing deferasirox (114 mg, 0.31 mmol), 10-aminodecylphosphonium chloride hydrochloride (150 mg, 0.31 mmol), N-(3-dimethyl aminopropyl)-N⍰ -ethylcarbodiimide hydrochloride (EDC) (88 mg, 0.46 mmol) and N, N-diisopropylethylamine (DIPEA) (0.32 ml, 1.86 mmol), and the reaction was stirred for 72 h at room temperature. Solvents were evaporated and the mixture was dissolved in methanol (MeOH) and washed with citric acid (5%, 7 mL), brine (7 mL), and dichloromethane (DCM, 2 × 20 mL). Column chromatography (2% MeOH/CHCl_3_) afforded the product (25 mg, 10 %) as a light-yellow foam (**Fig. S1A**).

### Cell viability assay

Viable cells were stained with crystal violet as described before.^24^

### Real-time cell proliferation and death monitoring

Live-cell imaging was performed using a Lumascope S720 (Etaluma) or IncuCyte® S3 (Sartorius). Images were captured for 72 h at 3 h intervals. SYTOX green (0.5 µM, Thermo Fisher Scientific) was used to detect dead cells. Cell proliferation and death analyses were performed using Lumaview/IncuCyte software and are presented as the percentage of confluence or the number of dead cells over time. All results are shown as normalized confluence/time zero for proliferation and normalized dead counts/phase for cell death measurement.

### Flow cytometry

For all flow cytometry experiments, cells were analyzed using a BD LSRFortessa™ flow cytometer (BD Biosciences). Annexin V/propidium iodide (AV/PI; Biolegend/0.5 μg/ml MERCK) double staining was measured at 489 nm_Ex_/515 nm_Em_ for AV and 534 nm_Ex_/617 nm_Em_ for PI. The percentage of dead cells is represented as the sum of AV+/PI−, AV−/PI+, and AV+/PI+ cells. Cell cycle was determined using Vybrant DyeCycle™ Violet Stain (5 μM; Thermo Fisher Scientific) at 405 nm_Ex_/437 nm_Em_. Cellular ROS levels were measured using 2⍰, 7⍰ -dichlorofluorescein diacetate (DCF-DA; 5 µM; Sigma-Aldrich) at 488 nm_Ex_/585 nm_Em_ while mitochondrial ROS levels were determined using the MitoSOX probe (2.5 μM; Thermo FisherFisher Scientific) at 488 nm_Ex_/530nm_Em_. Tetramethylrhodamine methyl ester (TMRM; 5 µM; Sigma-Aldrich) was measured at 561 nm_Ex_/586 nm_Em_. To assess mitochondrial membrane potential. Lipid peroxidation in the mitochondrial membrane was determined using the MitoPeDPP fluorescent dye (0.1 µM, Dojindo) and analysed at 488 nm_Ex_/530nm_Em_.

### Immunoblotting and antibodies information

Immunoblotting was performed according to standard protocols.^24^ The protein concentration was determined using a Bicinchoninic Acid Protein (BCA) assay (Thermo Fisher Scientific). Membranes were visualized using chemiluminescent substrates Western Bright Sirius (Advansta) or Clarity (BioRad) in an Azure c600 camera (Azure Biosystems). The antibodies used in this study are listed in **Table S2**.

### Mitochondrial isolation, solubilization, and Blue-Native polyacrylamide (BN-PAGE) electrophoresis

Mitochondrial isolation, solubilization and BN-PAGE were performed as described before.^27^ The antibodies used are listed in **Table S2**.

### Cellular ATP measurement

Total cellular ATP levels were measured using the CellTiter-Glo® Luminescent Cell Viability Assay kit (Promega), according to the manufacturer’s instructions. Graphs were generated by normalizing values to the total cell number determined by the Alamar Blue™ HS Cell Viability Reagent (Thermo Fisher Scientific) and relative to the control condition.

### Oxygen consumption rate measurement

The oxygen consumption rate (OCR) was measured using a Seahorse XFe96 (Agilent Technologies), according to the manufacturer’s protocol, as described previously.^28^ Data were normalized to cell numbers obtained using the ImageXpress Micro XLS analysis system (Molecular Devices).

### Confocal microscopy

Cellular and mitochondrial Fe^2+^ levels were assessed using FerroOrange (1 µM; Dojindo) and MitoFerroGreen (5 µM; Dojindo) probes, respectively, according to the manufacturer’s instructions. Hoechst 33342 (2 µM; MERCK) and MitoTracker Deep Red (20 nM; Thermo Fisher Scientific) were used for staining of nuclei and mitochondria, respectively. The cells were visualized under a Leica SP8 or a Nikon CSU-WI Spinning Disk confocal microscopes. Images were acquired using a 63x water immersion objective detected at 543 nm_Ex_/580 nm_Em_ for FerroOrange, 488 nm_Ex_/535 nm_Em_ for MitoFerroGreen, 640 nm_Ex_/663 nm_Em_ for MitoTracker and at 405 nm_Ex_/450 nm_Em_ for Hoechst.

To detect mitophagy in cells, the fluorescent Mtphagy Detection Kit (Dojindo) was used according to the manufacturer’s instructions. Cells were visualized using a 63x water immersion lens in a Leica SP8 confocal microscope at 561 nm_Ex_/650 nm_Em_ for Mtphagy dye and 488 nm_Ex_/510 nm_Em_, for Lyso dye.

The mitochondrial network was evaluated in MCF7 cells transfected with mitochondrial GFP. Cells were imaged using a 63x water immersion lens in a Leica SP8 confocal microscope. Fluorescence was detected at 405 nm_Ex_/450 nm_Em_ for Hoechst and 488 nm_Ex_/510 nm_Em_ for GFP. Quantification of the fluorescence signal, radial analysis and object counting were carried out using the ImageJ software, as described before.^29^

### Autoradiography

MCF7 and MDA-MB-231 cells were cultivated for 72 h in complete medium supplemented with ^55^Fe (added in the form of iron citrate at the ratio of 1:10, total activity 0.5 µCi/ml; Lacomed). Medium was replaced with medium containing 50 nM mitoDFX or 2 µM DFX and the cells were then treated for 60 h. Mitochondrial and cytosolic fractions were obtained as described above for blue native electrophoresis (BNE). Mitochondrial fraction was then solubilized with either 1% digitonin (preserving supercomplexes) or 1% Lauryl Maltoside (dissolving supercomplexes) for 1 h at 4°C and spun. Protein content of the supernatant was measured by the BCA method and 50 µg of proteins were separated by BNE. The gel was visualized *via* the Fuji imaging plate (GE Healthcare) using the Typhoon instrument (GE Healthcare).

### Quantitative Polymerase chain reaction (qPCR) with reverse transcription

RNA isolation was performed using RNAzol RT (Molecular Research Centre), according to the manufacturer’s instructions. The RNA quantity was measured using a NanoDrop spectrometer (Thermo Fisher Scientific, ND-1000). cDNA was generated using the RevertAid RT Reverse Transcription Kit (Fermentas), according to the manufacturer’s instructions. Quantitative PCR (qPCR) was performed using a 5x HOT FIREpol Eva Green qPCR mix kit (Solis BioDyne) in 384-well plates. Triplicate samples were run on a c1000 Thermal cycler (BIORAD). Data were analyzed using GenEx software version 6 and normalized to the reference gene *RPLP0*. The sequences and primers used for qPCR are listed in **Table S3**.

### Determination of GSH and GSSG levels

The levels of reduced glutathione (GSH), total glutathione (GSH+GSSG), and the GSH/GSSG ratio were measured using the GSH/GSSG-Glo assay kit (Promega) according to the manufacturer’s instructions. The amounts of total glutathione and oxidized glutathione (GSSG) were normalized to the cell number, which was determined using the Alamar Blue™ HS Cell Viability Reagent (Thermo Fisher Scientific). Determination of NAD+(P+)/ NAD(P)H levels Cellular NAD^+^/NADH and NADP^+^/NADPH ratios were measured using the NAD/NADH-Glo Assay (Promega) and NADP/NADPH-Glo Assay kit (Promega), respectively, according to the manufacturer’s instructions. The amounts of NADH, NAD^+^, NADPH, and NADP^+^ were normalized to the cell number, which was assessed using the Alamar Blue™ HS Cell Viability Reagent (Thermo Fisher Scientific).

### Proteomic nLC-MS 2 analysis

Samples were homogenized and lysed by boiling at 95 °C for 10 min in 100 mM Triethylammonium bicarbonate (TEAB) containing 2 % sodium deoxycholate (SDC), 40 mM chloroacetamide, and 10 mM Tris(2-carboxyethyl)phosphine, and further sonicated (Bandelin Sonoplus Mini 20, MS 1.5). Protein concentration was determined using the BCA protein assay kit (Thermo Fisher Scientific), and 30 µg of protein per sample was used for sample preparation. The samples were further processed using SP3 beads according to Hughes et al.^30^

All data were analyzed and quantified using MaxQuant software (version 1.6.12.0).^31,32^ The false discovery rate (FDR) was set to 1 % for both proteins and peptides, and a minimum peptide length of seven amino acids was specified. The Andromeda search engine was used to search MS/MS spectra against the human database (downloaded from UniProt.org, containing 20,598 entries). Data analysis was carried out by the Perseus 1.6.15.0 software.^33^

### Metabolomic analysis

For metabolomic assays, cells were grown in a 100 mm petri dish, washed with ice-cold 0.9% sodium chloride (NaCl) solution, and metabolites were extracted with 500 µL 80% methanol + 2 µg/ml ribitol (internal standard). Protein quantification of the resulting pellets was performed for normalization. Next, the samples were processed as proteomic samples^34^ with minor modifications available upon request. Data were processed using the Skyline software.

### Mice experiments

NOD scid gamma mice (NSG) and BALB/c mice were injected with 1×10^6^ MDA-MB-231 and 4T1 cells, respectively. C57BL/6 mice were injected with 2×10^5^ B16 cells. When the tumor volume reached 30-50 mm^3^ (tumor size quantified using ultrasound imaging, USI), each group was further divided into two subgroups and treated intraperitoneally (i. p.) with either vehicle (2.5 % DMSO in corn oil, 100 µL per dose) or mitoDFX (1 mg/kg or 0.25 mg/kg in corn oil) twice per week. Moreover, another group of C57BL/6 mice was administered 1.5 µM mitoDFX orally (roughly equivalent to a 0.25 mg/kg dose) continuously in drinking water. The tumor volume was monitored using the USI instrument Vevo3100 (VisualSonics). At the end of the experiment, mice were sacrificed and samples were obtained from the liver, spleen, blood, and tumor and analysed. Maximal tumor size allowed by ethics committee was 1000 mm^3^.

### Statistics

Statistical information for individual experiments can be found in the corresponding figure legends. All results are expressed as mean ± SEM of at least three independent experiments or 5 different animals. The comparison between treated samples and control was performed by one-way ANOVA or two-way ANOVA followed by Tukey’s multiple comparison test or Student’s t-test, using GraphPad Prism software version 10.1.1. The minimum significance level was set at P < 0.05. No statistical methods were used to determine the sample sizes.

## Results

### MitoDFX exhibits anti-proliferative and cytotoxic effects against breast cancer cells in vitro

The iron chelator deferasirox (DFX) inhibits the growth of several types of cancer cells *in vitro* and *in vivo*.^20-22^ To deliver DFX into mitochondria, we synthesized its mitochondrially targeted analogue, mitoDFX, by tagging the chelator with a triphenylphosphonium (TPP^+^) moiety (**Fig. 1A**).

**Figure 1:**
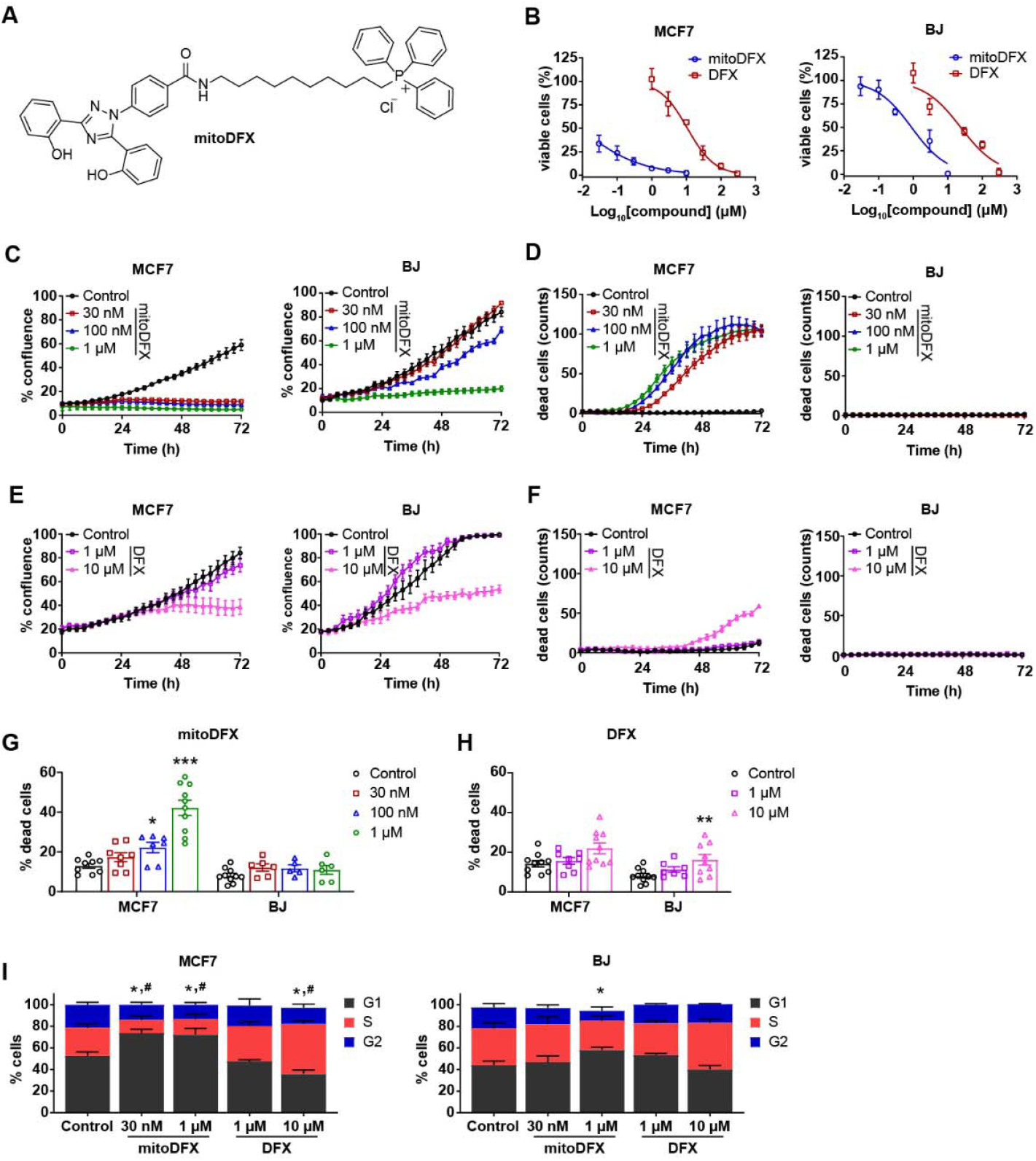
MitoDFX treatment induces cytostatic and cytotoxic effects in breast cancer cells. **(A)** Structure of mitoDFX. **(B)** Viability of MCF7 breast cancer cells and BJ human fibroblasts treated with mitoDFX and DFX for 48 h. Real-time monitoring of **(C)** proliferation and **(D)** cell death of MCF7 and BJ cells treated with mitoDFX. Cell death was measured using Sytox green dye (0.5 µM) and dead cells were counted in one field per well. Proliferation **(E)** and cell death **(F)** of MCF7 and BJ cells treated with DFX for 72 h using real-time LumaScope 720 microscope. Cell death was measured using Sytox green dye (0.5 µM). The percentage of dead MCF7 and BJ cells treated with **(G)** mitoDFX and **(H)** DFX was determined by annexin V and propidium iodide staining after 48 h. **(I)** Cell cycle distribution was assessed in MCF7 and BJ cells treated with indicated concentrations of mitoDFX and DFX and analyzed by FlowJo software. All data represents mean ± SEM of three independent experiments with at least two replicates each. P values were calculated using one-way **(G-H)** or two-way **(I)** ANOVA, followed by Tukey’s multiple comparisons test. \*\**P* < 0.01, \*\*\**P* < 0.001 relative to Control and * *P* ⍰ 0.05 relative to the G_1_ phase in the control; ^#^ P ⍰ 0.05 relative to the S phase in the control **(I)**.

First, to examine the anti-proliferative activity of mitoDFX and compare it with the parental agent, we used a panel of cancer cell lines and non-malignant fibroblasts. MitoDFX significantly reduced the number of viable cancer cells at nM concentrations, showing an almost three-order magnitude increase in efficacy compared to DFX (**Fig. 1B, S1B, Table S1**). We further performed real-time monitoring of cell proliferation and death to study the kinetics of the mitoDFX effect. The results demonstrated a cytostatic and cytotoxic effect on breast cancer cells, not affecting BJ fibroblasts (**Fig. 1C-D, S1C**). (**Fig. 1D**), demonstrating the selectivity against cancer cells (**Table S1**). In contrast, treatment with 10 µM DFX showed markedly lower cytostatic and cytotoxic activity (**Fig. 1E, F, S1D**). Enhanced cytotoxic effect in malignant breast cancer cells was further confirmed by Annexin V/PI staining (**Fig. 1G-H, S1E-F**). Moreover, 30 nM mitoDFX treatment for 72 h arrested proliferation at the G_1_ phase of the cell cycle in MCF7 but not in BJ (**Fig. 1I, S1G**) with DFX (10 µM) unable to elicit a similar response (**Fig. 1I, S1G**). Overall, these results illustrate the significant cytostatic and cytotoxic effects of nM concentrations of mitoDFX against cancer cells while sparing non-malignant cells.

### MitoDFX specifically reduces mitochondrial iron level in malignant cells

As mitoDFX is an iron chelator, we explored whether it alters the level of intracellular Fe^2+^. We demonstrated that treatment with 50 nM mitoDFX for 60 h did not affect the total intracellular Fe^2+^ level. When treated with 1 µM mitoDFX, a significantly reduced level of free intracellular Fe^2+^ was seen in breast cancer cells, whereas no change was observed in BJ cells (**Fig. 2A, B, S2A-D**).

**Figure 2:**
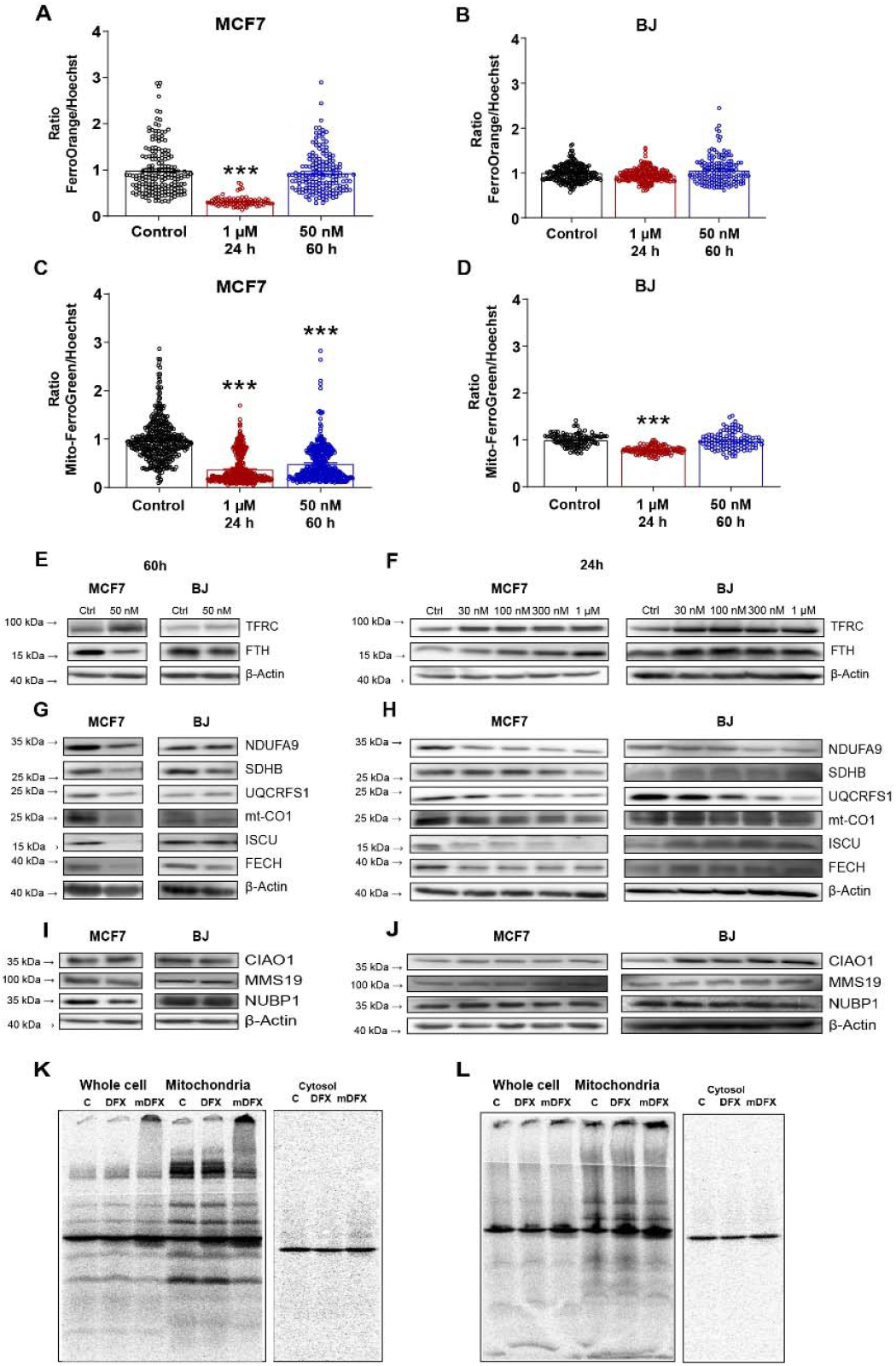
MitoDFX disrupts iron metabolism. **(A, B)** Quantification of the ratio of FerroOrange/Hoechst fluorescence relative to the control in MCF7 and BJ cells. **(C, D)** Quantification of the ratio of Mito-FerroGreen/Hoechst fluorescence relative to control in MCF7 and BJ cells. **(E-F)** Western blot images of proteins related to iron metabolism in MCF7 and BJ cells treated with mitoDFX for long (60 h) and short term (24 h). **(G-H)** Western blot images of mitochondrial [Fe-S] cluster and heme-containing proteins in MCF7 and BJ cells treated with mitoDFX for long (60 h) and short term (24 h). **(I-J)** Western blot images of cytosolic [Fe-S] cluster biogenesis-related proteins obtained from MCF7 and BJ cells exposed to mitoDFX for long (60 h) and short term (24 h). **(K-L)** Autoradiograph of 55Fe loaded cells and also a mitochondrial fractioon treated with either DFX or mitoDFX and solubilized either with digitonin (K) or lauryl maltoside (L).All data represent mean ± SEM of three independent experiments with at least 50 cells analyzed per condition in panels A-D. P values were calculated by one-way ANOVA followed by Tukey’s multiple comparisons test **(A-D)**. \*\*\**P* < 0.001 relative to Control.

Moreover, a considerable decrease in mitochondrial Fe^2+^ level in both malignant cells, after 50 nM 60 h mitoDFX treatment was observed, similarly to the 24 h incubation with 1 µM mitoDFX (**Fig. 2C, S3A-C**). BJ fibroblasts exhibited no significant change in mitochondrial Fe^2+^ level after 50 nM 60 h incubation and only a mild decrease after 24 h treatment with 1 µM mitoDFX (**Fig. 2D, S3D**). These data demonstrate that nM concentration of mitoDFX specifically reduce mitochondrial Fe^2+^ in cancer cells, highlighting its targeted effects on mitochondria and its selectivity for malignant cells.

### MitoDFX induces iron deprivation and destabilizes [Fe-S] cluster and heme-containing proteins in cancer cells

Next, we evaluated the effects of mitoDFX on cellular iron metabolism. Treatment in cancer cells with 50 nM mitoDFX for 60 h upregulated transferrin receptor 1 (TFRC) and decreased ferritin (FTH), while BJ cells remained unchanged (**Fig. 2E, S3E**). A similar trend was observed after 24 h exposure to 1 µM mitoDFX, except for increased FTH levels in MCF7 cells (**Fig. 2F, S3F**).

Furthermore, treatment with 50 nM mitoDFX for 60 h significantly reduced the levels of iron-containing proteins involved in the electron transport chain (ETC) (NDUFA9, SDHB, UQCRFS1, mtCO1) as well as heme and [Fe-S] cluster biogenesis (ISCU, FECH) in malignant cells. In contrast, these changes were absent in BJ fibroblasts (**Fig. 2G, S3G**). Similar changes were observed after 24 h incubation with 1 µM mitoDFX (**Fig. 2H, S3H**). Yet, protein levels of the cytosolic [Fe-S] assembly pathway proteins (CIAO1, MMS19, and NUBP1) remained unaltered in all tested cell lines (**Fig. 2I, J, S3I-J**).

Autoradiography of cells loaded with ^55^Fe has confirmed that iron-containing proteins are reduced after mitoDFX treatment, both in conditions that maintain mitochondrial respiratory supercomplexes (digitonin solubilization; **Fig. 2 K**), as well as in those that disrupt them (lauryl maltoside solubilization; **Fig. 2L**).

These results demonstrate that mitoDFX deprives cells of the available mitochondrial iron pool and reduces the levels of mitochondrial [Fe-S] cluster- and heme-containing proteins. Importantly, the cytosolic [Fe-S] cluster assembly machinery remained intact, indicating a specific effect on mitochondria.

### MitoDFX decreases the amount and assembly of mitochondrial respiratory supercomplexes, causing a reduction in oxygen consumption

Noticing the decrease in [Fe-S]- and heme-containing subunits of the ETC, we further investigated the effect of mitoDFX on its functionality. We have observed a reduced amount and assembly of respiratory supercomplexes in malignant cells treated with mitoDFX (**Fig. 3A, S4A**), which corroborates results from autoradiography. In contrast, these changes were absent in non-malignant BJ with lower mitoDFX concentrations and became apparent only at 1 µM (**Fig. 3A**). Our results showed that mitoDFX inhibited mitochondrial oxygen consumption at 1 µM concentration in malignant cell lines, diminishing both basal and maximal respiration with only a minor impact on BJ cells (**Fig. 3B, S4B**). Moreover, a significantly higher dose of DFX was required to impair the ETC (**Fig. 3C, S4C**).

**Figure 3:**
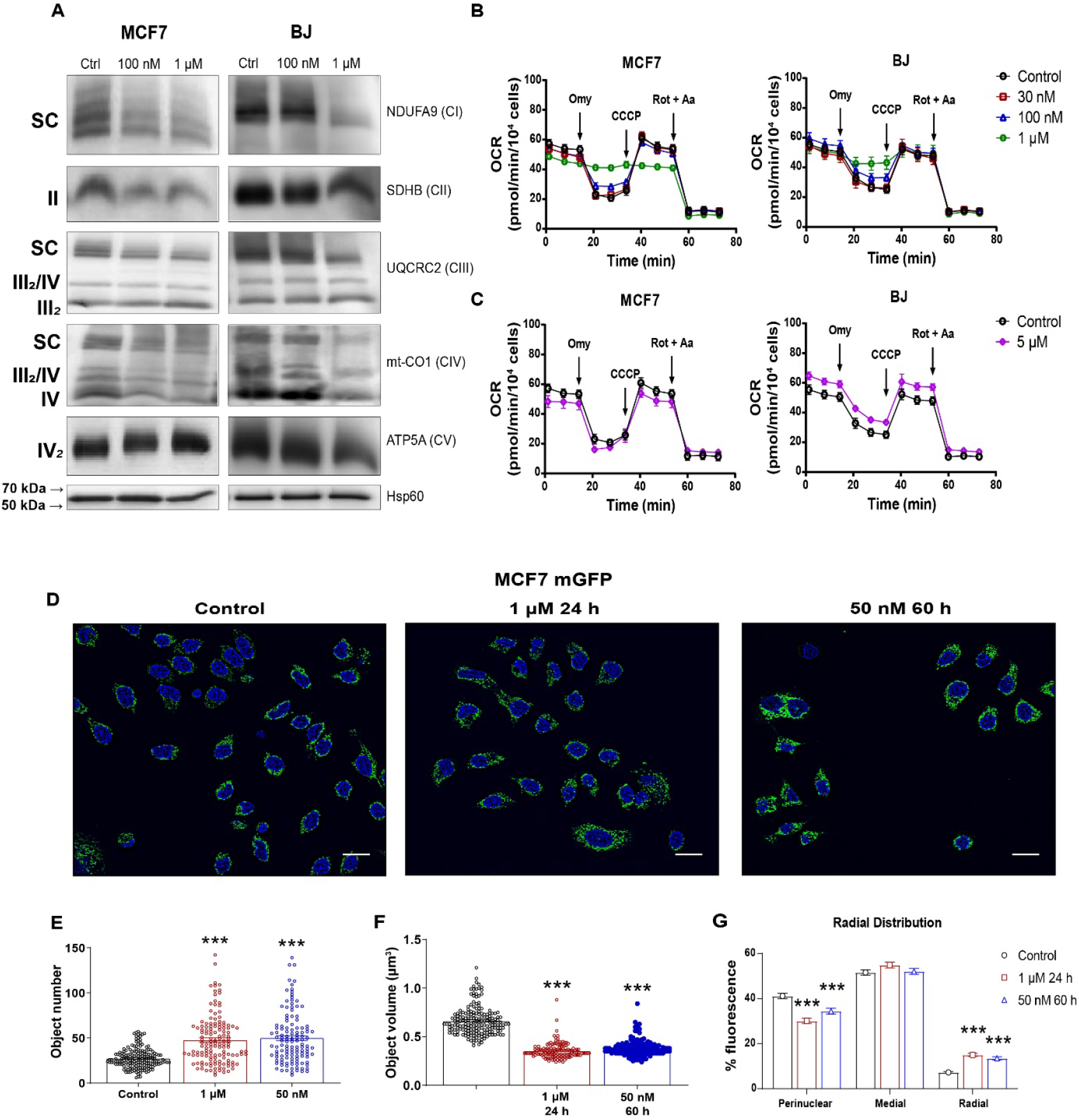
MitoDFX disrupts assembly of mitochondrial respiratory supercomplexes, inhibits mitochondrial oxygen consumption and induces mitochondrial fragmentation and redistribution. **(A)** BN-PAGE of the OXPHOS complexes from isolated mitochondrial samples obtained from MCF7 and BJ cells treated with mitoDFX for 24 h. Hsp60 was used as a loading control. The oxygen consumption rate (OCR) profile of MCF7 and BJ cells treated with mitoDFX **(B)** or DFX **(C)** was obtained using an extracellular flux analyzer (Seahorse XF). **(D)** Confocal images of MCF7-mitoGFP cells subjected to treatment with mitoDFX for 24-60 h. Nuclei were stained with Hoechst 33342. Scale bars = 20 µm. Quantification of mitochondrial number **(E)** and average volume **(F)** in MCF7-mitoGFP transfected cells treated with mitoDFX for 24-60 h. **(G)** Quantification of mitochondrial distribution in MCF7-mitoGFP cells exposed to mitoDFX for 24-60 h. All data represent mean ± SEM of three independent experiments with at least two replicates each. *P* values were calculated by one-way **(E-F)** or two-way ANOVA **(G)** followed by Tukey’s multiple comparison. \*\**P* < 0.01, \*\*\**P* < 0.001 relative to Control.

Additionally, the simultaneous treatment with mitoDFX and 2-deoxyglucose (2-DG) significantly depleted ATP level in malignant cells compared to BJ cells. However, mitoDFX treatment alone or its combination with oligomycin did not alter ATP levels in any cell lines, indicating a compensatory shift to glycolysis to replenish the ATP pool despite the loss of mitochondrial respiration (**Fig. S4D**).

### MitoDFX induces a decrease in mitochondrial membrane potential, mitochondrial fragmentation and redistribution

Mitochondrially targeted drugs containing the TPP^+^ moiety usually affect mitochondrial membrane potential (MMP) and mitochondrial function.^24,35,36^ We have found significantly reduced MMP in all tested cells after treatment with 50 nM mitoDFX for 60 h (**Fig. S5A**); however, 24 h treatment reduced MMP only in malignant cancer cells (**Fig. S5B**). Given the profound changes seen in the ETC and MMP, we next examined the effect of mitoDFX on mitochondria structure and dynamics.^37^ We have demonstrated that mitoDFX treatment significantly increased the number of mitochondria and decreased their mean volume (**Fig. 3D-F**). Furthermore, mitoDFX caused a shift of mitochondria from the perinuclear compartment to the radial regions of the cell (**Fig. 3G**). Together, these results indicate mitochondrial fragmentation and redistribution, accompanied by enhanced mitophagy in malignant cells (**Fig. S5C**) coupled with increased PINK1, BNIP3 and LC3 II (**Fig. S5D**).

Overall, mitoDFX induces mitochondrial dysfunction that involves the loss of MMP, induction of mitochondrial fission, and markedly increased mitophagy in malignant breast cancer cells only.

### MitoDFX alters mitochondrial proteome and impedes other mitochondrial processes, including translation, replication and transcription

We carried out a proteomic analysis and found that 50 nM mitoDFX significantly reduced levels of many mitochondrial proteins, unlike the non-targeted DFX (**Fig. 4A**).

**Figure 4:**
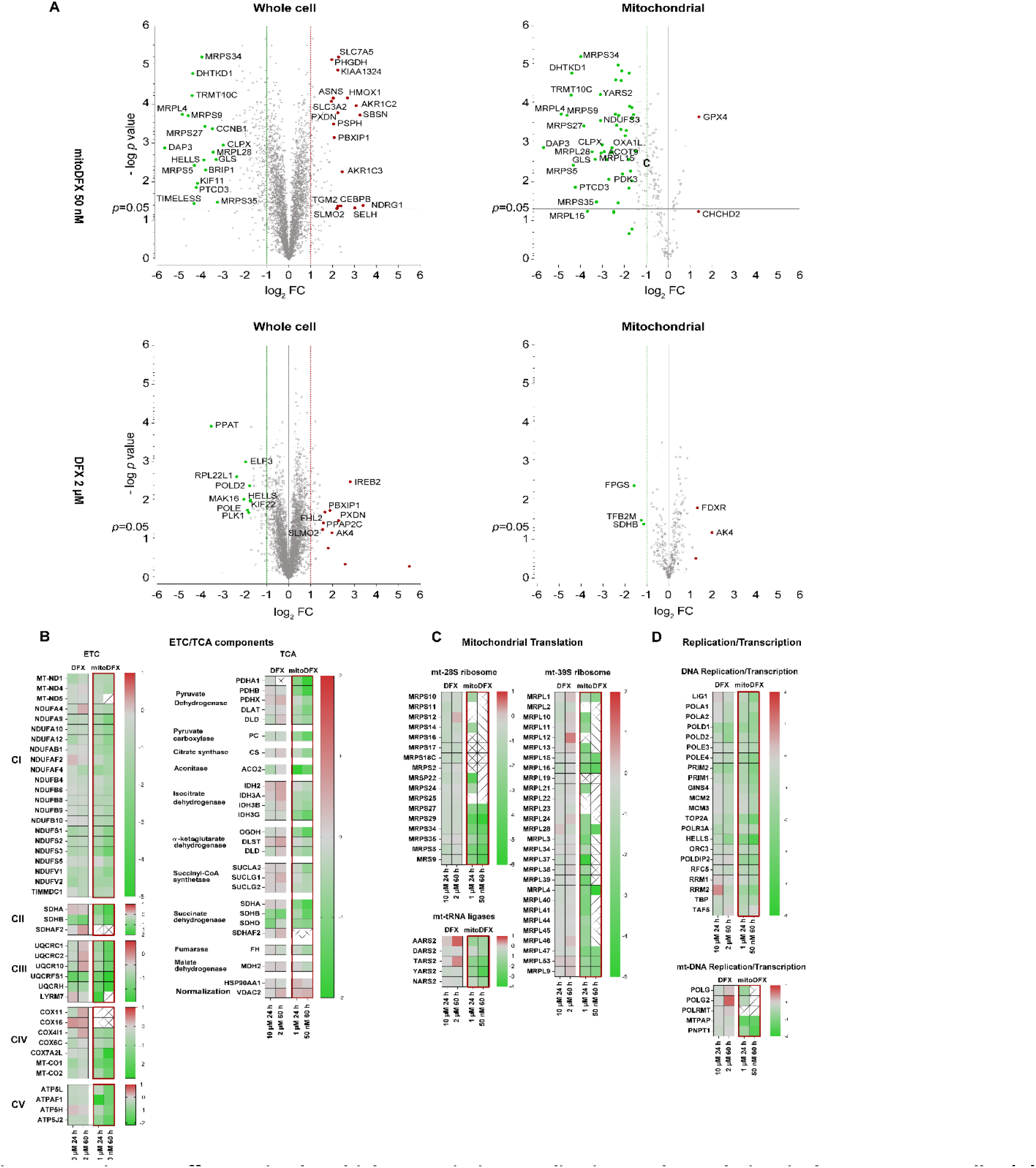
MitoDFX affects mitochondrial transcription, replication and translation in breast cancer cells. **(A)** Volcano plots showing volcano plots of proteins detected by LC-MS in cells treated with **(A)** mitoDFX 50 nM or **(B)** DFX 2 µM for 60 h. The red points represent proteins upregulated by the treatments (p-value < 0.05, |fold change|> 2). The green points represent the downregulated proteins with the treatments (p-value < 0.05, |fold change|> 2). Proteomic data demonstrating dysregulated pathways, including ETC/TCA components **(B)**, mitochondrial translation **(C)**, and DNA replication/transcription **(D)**. All data represent mean ± SEM of three independent experiments with at least three replicates each. *P* values were calculated by two-way ANOVA **(F)**. *** *P* < 0.001 relative to Control.

According to the Reactome pathway knowledgebase^38^, most significantly affected pathways after treatment with 50 nM mitoDFX were mitochondrial translation, mitochondrial electron transport chain (ETC), TCA cycle and DNA replication/transcription machinery especially in mitochondria (**Fig. 4B-D, S6A-C**). In detail, mitoDFX treatment reduced the protein subunits of all five mitochondrial respiratory complexes and enzymes of the TCA cycle (**Fig. 4B**). Furthermore, we detected a dramatic reduction in the subunits of mitochondrial ribosomes and several tRNA ligases involved in mitochondrial translation (**Fig. 4C**) and inhibition of proteins connected with both general and mitochondrial transcription and DNA replication (**Fig. 4D**). In line with reduced components of the mitochondrial transcription machinery, we have detected markedly reduced levels of mitochondrial transcripts (**Fig. S6D**). A detailed analysis of the affected pathways is presented in **Fig. S6C** and the complete analysis is available as **Supplementary File 1**.

### MitoDFX inhibits the tricarboxylic acid cycle (TCA)

Considering the notable deregulation of components of the ETC and TCA cycle identified by Reactome analysis, we examined the impact of mitoDFX on glycolysis and TCA cycle intermediates levels. MitoDFX treatment led to a significant decrease in glycolysis products, consistent with reduction infructose-1,6-bisphosphate in treated cells. Similarly, TCA cycle intermediates, such as citric acid, isocitric acid, succinyl-CoA, fumarate, and malate were drastically decreased in treated samples. Conversely, acetyl-CoA, α-ketoglutarate, and succinate remained unchanged after treatment (**Fig. 5**). These findings suggest that the TCA cycle no longer runs as a canonical cycle, impairing its ability to support associated metabolic processes, such as lipogenesis, urea cycle, or amino acid metabolism.

**Figure 5:**
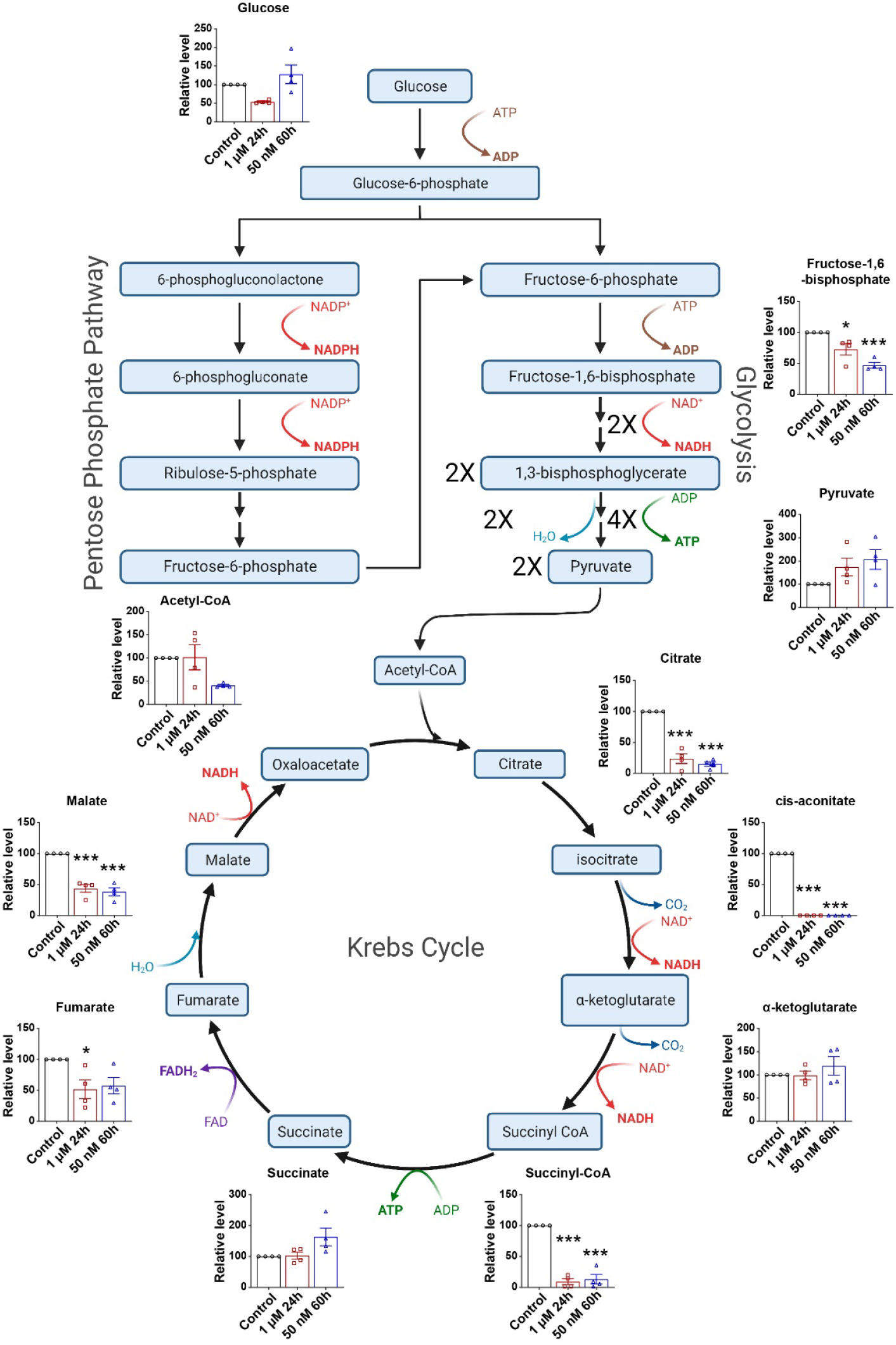
MitoDFX affects glycolysis and TCA cycle. Representative diagram of glycolysis, PPP pathway and TCA cycle showing the metabolite levels detected in MCF7 cells treated with mitoDFX for short (24 h, 1 µM) and long term (60 h, 50 nM). Data is expressed as means ± SEM of three independent samples. *P* values were evaluated by one-way ANOVA followed by Tukey’s multiple comparisons test. \**P* < 0.05, *** *P* < 0.001 relative to Control.

### MitoDFX increases ROS levels and induces lipid peroxidation

Since oxygen consumption and metabolic functions of mitochondria are compromised by mitoDFX treatment, we further assessed the levels of reactive oxygen species (ROS), which can damage biomolecules, including lipids.^39^ Not only did the treatment with 50 nM mitoDFX induce mitochondrial superoxide levels in MCF7 and MDA-MB-231 cells **(Fig. 6A, S7A)**, but it also increased overall cellular ROS levels **(Fig. 6B, S7B)**. Importantly, ROS levels remained unchanged in BJ cells after long-term treatment with mitoDFX **(Fig. 6A, B)**, confirming its selectivity. Conversely, mitochondrial and cellular ROS levels were elevated in all cell lines after treatment with higher concentrations (1 and 5 µM) for 24 h **(Fig. S7C, D)**. We found significantly elevated mitochondrial lipid peroxidation (*via* MitoPeDPP^40^) in both breast cancer cells following 60 h treatment compared to BJ fibroblasts **(Fig. 6C, S7E)**. Similar results were obtained after 1 µM mitoDFX treatment for 1 h with an increase in all cell lines **(Fig. S7F)**. However, at 1 µM, mitoDFX selectively killed cancer cells but not BJ cells **(Fig. 1D, S1C)**, suggesting that these cells have more efficient mechanisms to cope with lipid peroxidation.

**Figure 6:**
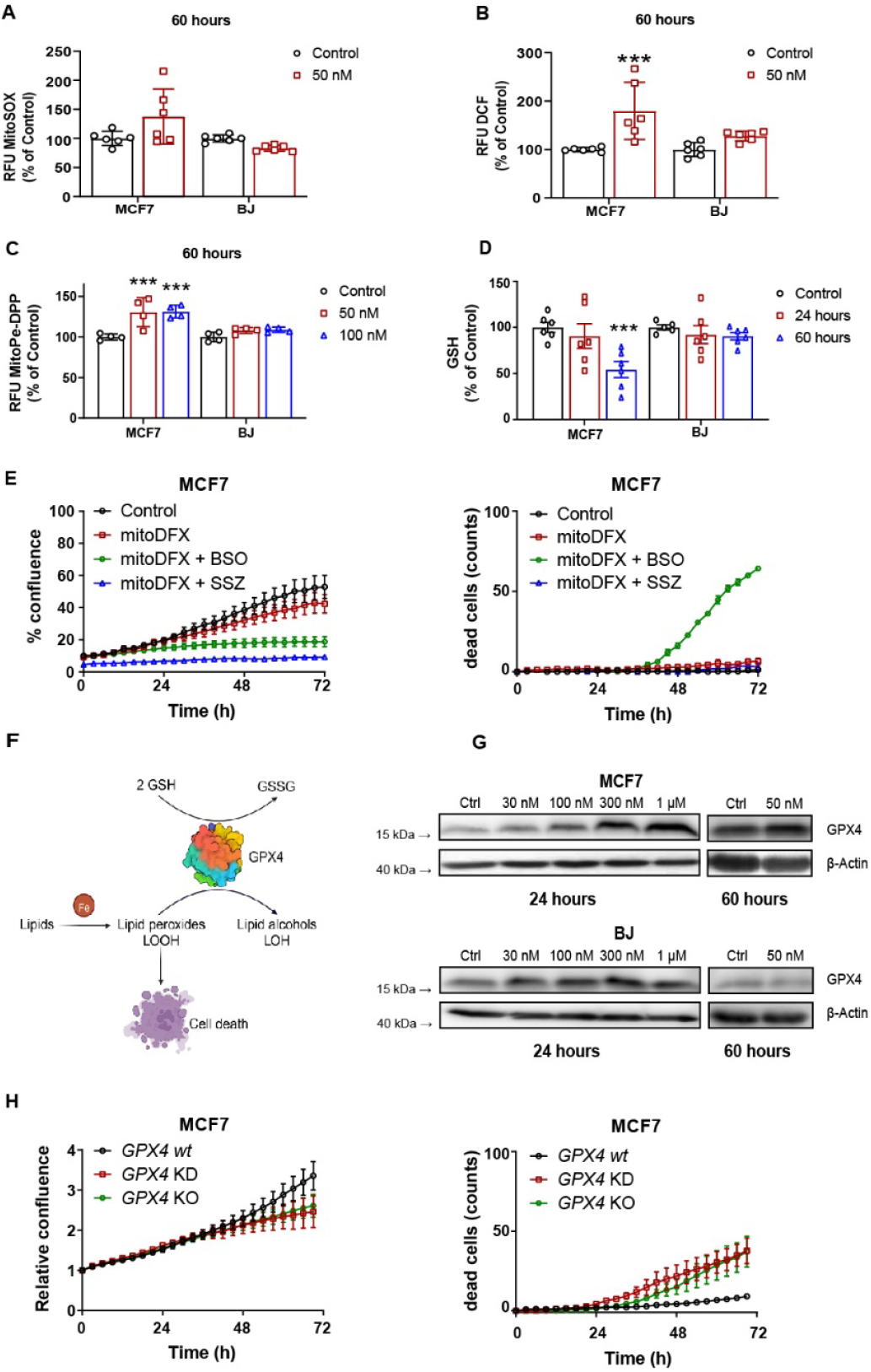
MitoDFX induces ROS levels and mitochondrial lipid peroxidation, decreases reduced glutathione and increases GPX4, deletion of which exacerbates mitoDFX induced cell death. Quantification of **(A)** mitochondrial superoxide and **(B)** total cellular ROS levels after treatment with mitoDFX for 60 h in MCF7 and BJ cells. **(C)** Quantification of lipid peroxidation after mitoDFX treatment for 60 h in MCF7 and BJ cells. **(D)** Quantification of reduced glutathione (GSH) levels in MCF7 and BJ cells incubated with mitoDFX (50 nM) for short and long term. **(E)** Real-time monitoring of proliferation and cell death of MCF7 cells treated with mitoDFX (10 nM) in combination with buthionine sulfoximine (BSO; 100 µM) or sulfasalazine (SSZ; 500 µM) for 72 h. Cell death was measured using Sytox green dye (0.5 µM). **(F)** Schematic representation of the role of GPX4 in lipid peroxide detoxification. **(G)** Western blot images of GPX4 protein levels in MCF7 and BJ cells treated with mitoDFX for 24-60 h. **(H)** Real-time monitoring of proliferation and cell death of MCF7 (10 nM) *GPX4* KD and KO cells incubated with mitoDFX for 72 h. Cell death was measured using Sytox green dye (0.5 µM). All data represent mean ± SEM of three independent experiments with at least two replicates each. *P* values were calculated by two-way ANOVA **(C-D)** followed by Tukey’s multiple comparison, two-tailed unpaired t-test **(A-B)**. \*\**P* < 0.01, \*\*\**P* < 0.001 relative to Control.

### MitoDFX induces exhaustion of reduced glutathione

Cellular protection against oxidative stress relies largely on glutathione (GSH).^41^ We measured GSH levels and observed significant reductions after 60 h mitoDFX exposure in MCF7 malignant cells but not in BJ cells **(Fig. 6D, S7G)**. We evaluated total glutathione levels and observed a significant reduction only in MCF7 and MDA-MB-231 cells treated with 50 nM mitoDFX for 60 h but not in BJ cells **(Fig. S7H)**. Moreover, the GSH/GSSG ratio decreased in MCF7 cells treated with mitoDFX for 60h **(Fig. S7I)**. Interestingly, no significant changes were observed in MDA-MB-231 and BJ cells after mitoDFX treatment for 24-60 h. This suggests that the shift in the GSH/GSSG ratio as a sign of oxidative damage might be a critical factor leading to cell death, as MCF7 cells have IC_50_ values below 50 nM **(Fig. 1C, D)** Conversely, MDA-MB-231 and BJ cells have IC_50_ values above 50 nM, showing either a cytostatic response (MDA-MB-231) or no effect (BJ) **(Fig. 1C, D, S1C)**.

### Depletion of glutathione and loss of GPX4 sensitizes cells to mitoDFX-induced death

To confirm the importance of reduced glutathione in the effect of mitoDFX, we used buthionine sulfoximine (BSO), a potent inhibitor of γ-glutamylcysteine synthetase (γ-GCS).^42^ Co-treatment of BSO and mitoDFX showed enhanced cytostatic and cytotoxic effects in MCF7 cells **(Fig. 6E)**. Similarly, combining sulfasalazine (SSZ), an inhibitor of the xCT-cystine/glutamate antiporter, with mitoDFX resulted in significantly stronger cytostatic effect in MCF7 cells, suggesting synergy of the two **(Fig. 6E)**. We further focused on GPX4, a selenium-containing protein that protects cells from lipid peroxides using GSH as a substrate^41^ **(Fig. 6F)**, as its level was significantly induced by 50 nM mitoDFX exposure for 60 h in MCF7 and MDA-MB-231 cells, but not in BJ cells, suggesting a link between the accumulation of lipid peroxides and the induction of GPX4 **(Fig. 6G, S7J)**. To confirm that GPX4 induction is related to mitoDFX sensitivity, we knocked out (KO) or down (KD) *GPX4* in MCF7 cells **(Fig. S7K)**. We observed enhanced cell death in *GPX4* KO and KD cells after treatment with 10 nM mitoDFX, while wild-type cells remained unaffected, indicating an important role of GPX4 in mitigating mitoDFX-induced toxicity **(Fig. 6H)**. Overall, our results demonstrated that mitoDFX elevates ROS levels, leading to lipid peroxide generation, which is neutralized by GPX4 at the expense of GSH, eventually depleting GSH reserves, leading to the accumulation of toxic lipid peroxides and subsequent ferroptotic cell death.

### MitoDFX increases the NAD^+^ /NADH ratio and synergizes with the inhibitor of the pentose phosphate pathway

We further investigated the status of vital redox cofactors NAD^+^/NADH and NADP^+^/NADPH, which are crucial in regulating cellular redox status, glutathione metabolism, and mitochondrial function. MitoDFX treatment (50 nM, 60 h) led to a significant increase in the NAD^+^/NADH ratio in MCF7 cells. Less sensitive MDA-MB-231 cells showed this increase only after 24 h of 1 µM mitoDFX treatment, with no change in BJ cells **(Fig. S7L)**. However, the NADP^+^/NADPH ratio remained unaltered after mitoDFX treatment across all the tested cell lines **(Fig. S7M)**. It is well documented that cancer cells exhibit increased activity of the pentose phosphate pathway (PPP) that generates NADPH to counteract elevated ROS levels by facilitating the continuous reduction of GSSG ^43^. However, mitoDFX treatment did not alter the NADP^+^/NADPH ratio, and therefore, we assessed the potential protective role of the PPP against mitoDFX using 6-aminonicotinamide (6-AN), a PPP inhibitor. Interestingly, 6-AN markedly enhanced the cytotoxic effect of mitoDFX in MCF7 cells, confirming the active utilization of the PPP pathway to protect cells from oxidative damage **(Fig. S7N)**.

### MitoDFX inhibits tumor growth of syngeneic melanoma and TNBC cells, as well as xenografted human TNBC in vivo

Our *in vitro* results demonstrated a profound suppressive effect of mitoDFX on proliferation and its ability to induce cell death in breast cancer cell lines. Therefore, we assessed mitoDFX efficacy *in vivo* using syngeneic and xenograft mouse models. In the syngeneic mouse models (4T1 TNBC/Balbc mice and melanoma B16 cells/C57/BL6J mice), mitoDFX (intraperitoneal at 1 mg/kg or 0.25 mg/kg) significantly decreased tumor growth **(Fig. 7A)**. Similarly, in a breast cancer xenograft model (MDA-MB-231 cells), mitoDFX treatment (intraperitoneally at 1 mg/kg) markedly suppressed tumor progression **(Fig. 7B)**. Moreover, long-term oral treatment with mitoDFX considerably reduced tumor growth in a melanoma mouse model **(Fig. 7C)**. Hematological parameters in tumor-bearing mice showed no significant differences between control and mitoDFX-treated animals **(Fig. 7D)**. To confirm that mitoDFX targets cancer cells and does not affect systemic iron metabolism, we evaluated the liver, spleen, and tumor tissue iron levels in BALB/c, NOD scid gamma (NSG), and C57BL/6 mouse models. We did not observe any changes in these parameters in the treated animals, except for a decrease in B16 tumor iron level in the mitoDFX-treated group **(Fig. 7E)**. Additionally, the body weight of mitoDFX-treated mice remained unchanged, indicating the sustained well-being of the animals **(Fig. S8)**. We have also demonstrated that B16 tumors treated with mitoDFX show enhanced levels of GPX4 and 4HNE, thus supporting the role of lipid peroxidation in mitoDFX treatment *in vivo* **(Fig. 7 F)**. Likewise, we detected higher levels of TFRC and PINK1 and slightly lower level of NDUFA9 in mitoDFX-treated tumors, confirming our *in vitro* findings **(Fig. 7F**). Together, these results demonstrate that mitoDFX is an extremely effective drug *in vivo*, requiring significantly lower doses compared to other mitochondrially targeted drugs.^35,36,44^

**Figure 7:**
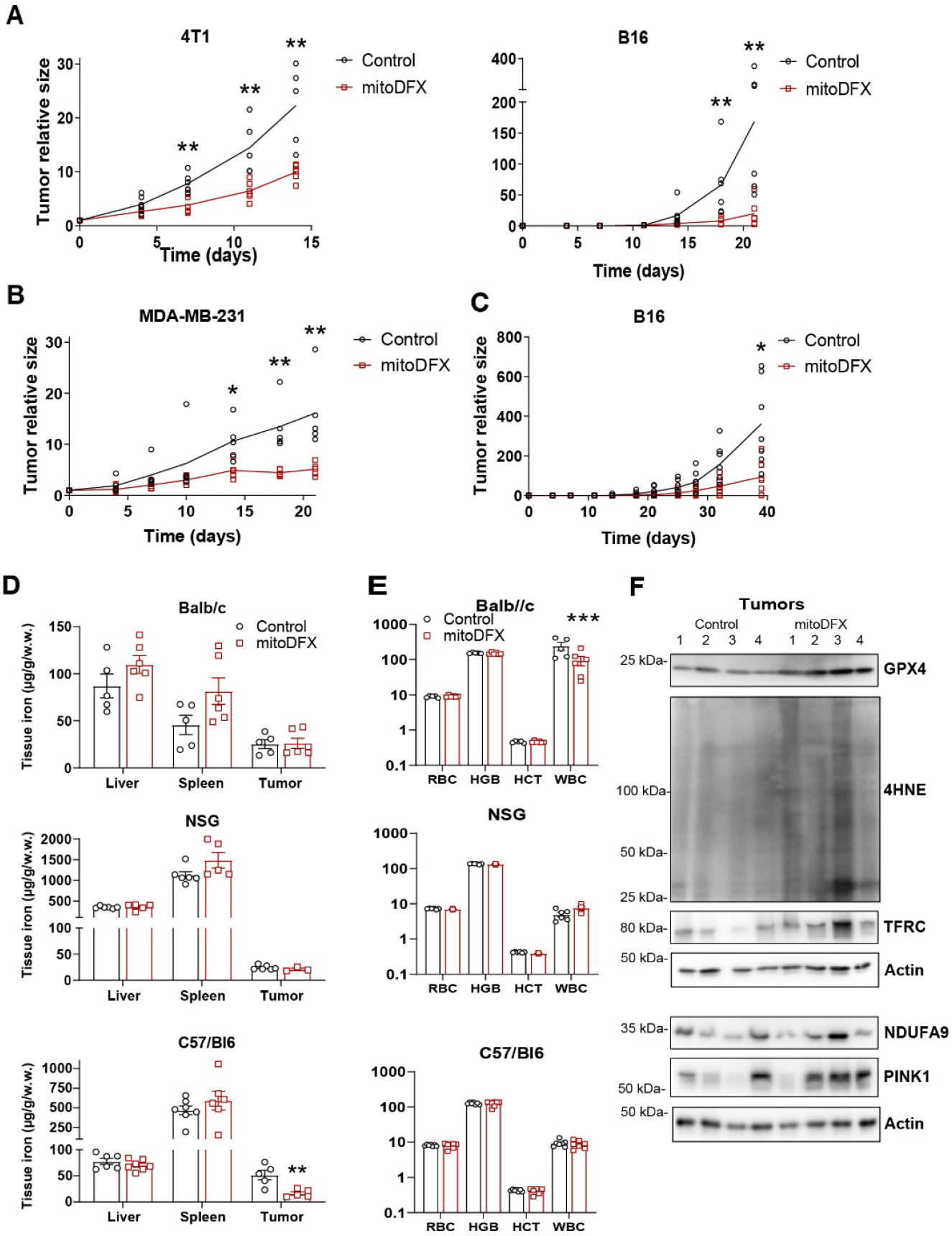
The effects of mitoDFX on tumor growth, hematological and systemic iron parameters in mice models. Tumor growth curves of mice injected subcutaneously with murine syngeneic tumor cells of **(A)** triple negative breast cancer 4T1 and melanoma B16. **(B)** Tumor growth charts of mice injected subcutaneously with murine melanoma B16 cells and continuously given mitoDFX in drinking water (1.5 µM) for 40 days. **(C)** Xenotransplantation of human triple negative breast cancer MDA-MB-231 cells in immunodeficient NSG mice treated with mitoDFX every 2 days. When the tumor volume reached approximately 5-30 mm^3^, the mice received mitoDFX i.p. (1 mg/kg for 4T1 and MDA-MB-231, and 0.25 mg/kg for B16). **D)** Quantification of total red blood cells (RBC), hematocrit (HCT), hemoglobin (HGB), and white blood cells (WBC) in BALB/c, NSG, and C57BL/6 tumor-bearing mice treated i.p. with mitoDFX. **(E)** The iron content of liver, spleen, and tumor in BALB/c, NSG, and C57BL/6 tumor-bearing mice treated i.p. with mitoDFX twice per week. All data represent the mean ± SEM of at least 5 animals. P values were calculated by a two-tailed unpaired t-test. * *P* ⍰ 0.05, ** *P* ⍰ 0.01, *** *P* ⍰ 0.001 relative to Control.

## Discussion

With the continuous development of novel anticancer drugs, selective targeting is one of the perspective options to improve treatment efficacy. In this report, we describe a novel mitochondrially targeted deferasirox (mitoDFX) that proves highly efficient both *in vitro* and *in vivo*. Its IC_50_ values range between 10-100 nM, a 1000-fold increase in efficacy compared to its parental compound. It is approximately 30 times more effective than previously described mitochondrially targeted deferoxamine, mitoDFO,^24^ and surpasses other mitochondrially targeted drugs.^35,36,44^ Its IC_50_ is similar to Dp44mT and DpC developed by the team of Prof. Richardson;^45,46^ however, mitoDFX shows cytotoxicity against most cancer cells at a 100 nM concentration, while the Dp44mT and DpC were mostly used at 5 µM.

Similarly, the salinomycin^47^ derivative ironomycin, which acts by sequestering iron in the lysosome and reducing mitochondrial iron,^48^ decreases cellular proliferation at 50 nM but does not induce cell death. Importantly, the same concentration of mitoDFX not only completely halts proliferation of the cancer cells but also kills them. MitoDFX seems to utilize a different mechanism of action that does not reduce mitochondrial iron *via* the lysosomal iron retention. Instead, mitoDFX utilizes unique mechanism of action that combines iron chelation with lipid peroxidation and depletion of glutathione, both of which are hallmarks of ferroptosis. In addition, mitoDFX is remarkably selective for malignant cells, which might reduce its side effects and increase efficacy in future cancer therapy.

The ability to chelate iron and make it unavailable for biochemical and biosynthetic processes seems to be shared with the previously described mitoDFO.^24^ We could see a similar pattern of increased mitochondrial superoxide production, decreased mitochondrial respiration, and overall mitochondrial dysfunction coupled with their fragmentation and induction of mitophagy. However, we see a higher efficacy of mitoDFX in inhibiting mitochondrial biosynthetic processes (mitochondrial transcription, replication and translation). Contrary to the expected NADH accumulation typically associated with adysfunctional ETC, we see a decrease. This suggests that NADH might be consumed by ETC-independent processes, maintaining NADPH level to support antioxidant defence mechanism and essential metabolic reactions.^49^ Additionally, enhanced cell death was observed when combining mitoDFX with inhibition of the pentose phosphate pathway (PPP), suggesting that the PPP produces reducing equivalents to counteract the oxidative stress induced by mitoDFX.

Importantly, we observed distinctive characteristics after mitoDFX treatment like its ability to upregulate glutathione peroxidase 4 (GPX4), a crucial mitochondrial enzyme that regulates ferroptosis,^41^ and its ability to induce mitochondrial lipid peroxidation both *in vitro* as well as *in vivo*. We found that GPX4/GSH axis plays an important role in the mode of action of mitoDFX as *GPX4* KO cells exhibited enhanced sensitivity to mitoDFX treatment. Furthermore, the inhibition of GSH production *with* BSO or sulfasalazine significantly enhanced the cytostatic/cytotoxic effect of mitoDFX treatment, suggesting that inhibiting the GSH/GPX4 axis could sensitize cancer cells to mitoDFX treatment.

In summary, mitochondrially targeted deferasirox (mitoDFX) has a unique ability to elicit iron deprivation as well as produce toxic lipid peroxides *via* its redox activity, thus harnessing the dual nature of iron in a single molecule to combat cancer **(Fig. 8)**. Importantly, it can effectively kill cancer cells at lower doses compared to other mitochondrially targeted drugs,^35,36,44^ and demonstrates high efficacy in animal models of TNBC and melanoma.

**Figure 8:**
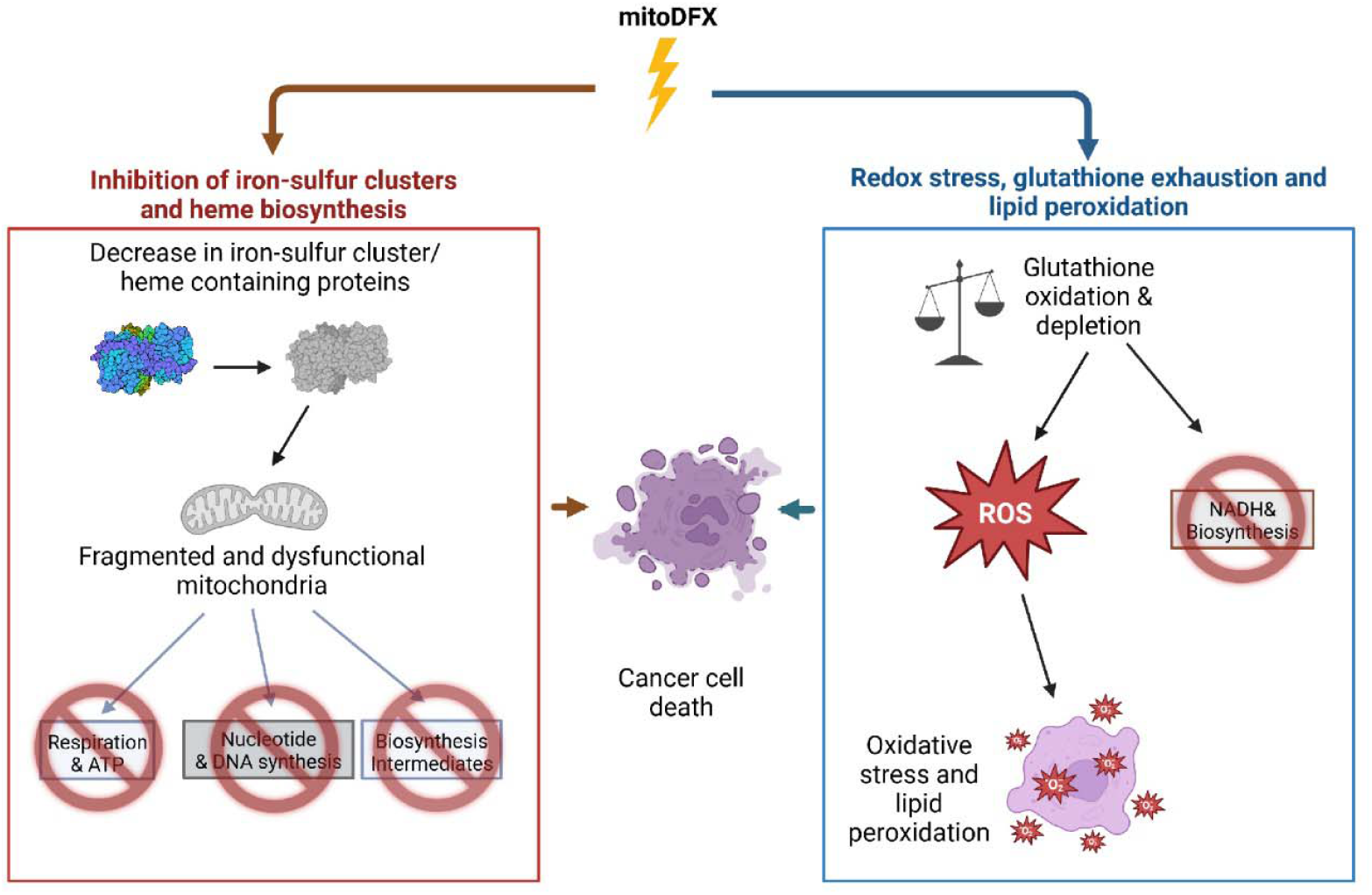
Graphical abstract depicting selective killing of cancer cells by simultaneously inhibiting iron-sulfur cluster and heme biogenesis, depleting glutathione and increasing mitochondrial lipid peroxidation elicited by mitoDFX. Our study shows novel mode of action that is elicited by mitochondrially targeted deferasirox (mitoDFX) to kill cancer cells. MitoDFX has the unique ability to simultaneously induce iron deprivation and deplete reduced glutathione, coupled with production of toxic lipid peroxides, therefore harnessing the dual nature of iron in a single molecule to combat cancer.

## Supporting information

Supplementary File1_Reactome

Supplementary File 2_raw proteomics

## Acknowledgments

We sincerely thank O. Souckova, P. Talacko, and K. Harant for the OMICS analyses (Laboratory of Mass Spectrometry, BIOCEV Research Center).

The project was funded by Czech Academy of Sciences RVO 86652036, National Institute for Cancer Research funded by the European Union Next Generation EU (Program EXCELES LX22NPO5102, MEYS), Czech Science Foundation 25-18052S to J.T. and by the GAUK Project No. 1310420 to S.B.J. We acknowledge support of the Imaging Methods Core Facility at BIOCEV by the MEYS CR (LM2018129 Czech-BioImaging). Animal experiments were supported by the Czech Academy of Sciences RVO 68378050 and grants OPVVVCZ.02.1.01/0.0/0.0/16_013/0001789 and LM2023036 from the Czech Centre for Phenogenomics (RS) provided by the MEYS of the Czech Republic.

## Authors’ contributions

**S.B. J**.: Performed most of the *in vitro* experiments and helped with all *in vivo* experiments, analyzed and interpreted the data, curated the figures, and wrote, reviewed, and edited the manuscript. **C.S.A**.: Helped with *in vitro* experiments and led the *in vivo* experiments, analyzed and interpreted the data, supervised and designed experiments, generated the figures, and wrote, reviewed, and edited the manuscript. **Y.P. and P.P**.: Performed *in vitro* experiments, helped with *in vivo* experiments, and reviewed, and edited the manuscript. **R.S**.: provided NSG mice and funding. **K.K**., **K.B**., **J.S**., **L.W**.: Design and synthesized the mitoDFX and helped with the experimental design. **J.T**.: Conceptualized and obtained funding for the project, supervised the study, established collaborations, helped in designing experiments, analyzed and interpreted the data, and wrote, reviewed, and edited the manuscript.

## Data availability statement

All data generated or analysed during this study are included in this published article and its supplementary information files. Raw proteomics data are provided in **Supplementary File 2**.

