## Supplementary Figures and Tables for "Selective killing of cancer cells via simultaneous destabilization of mitochondrial iron metabolism and induction of ferroptosis elicited by mitochondrially targeted iron chelator deferasirox"

A

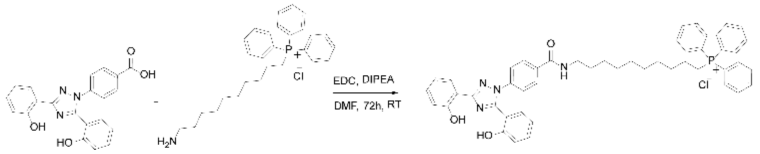


$^1$H NMR (500 MHz, Chloroform-d) δ 10.96 (bs, OH, 1H), 10.16 (bs, OH, 1H), 8.40 (bs, NH, 1H), 8.12 (t, J = 7.9 Hz, 2H), 7.79 – 7.68 (m, 6H), 7.65 (s, 7H), 7.42 (d, J = 6.9 Hz, 2H), 7.33 (t, J = 7.9 Hz, 1H), 7.25 (dd, J = 26.3, 7.7 Hz, 2H), 7.11 (d, J = 7.7 Hz, 1H), 7.00 (m, 2H), 6.74 (t, J = 7.4 Hz, 1H), 3.46 (m, 4H), 1.79 – 1.48 (m, 6H), 1.26 (m, 10H).

$^{13}$C NMR (126 MHz, Chloroform-d) δ 166.48, 159.61, 157.18, 156.65, 152.14, 139.81, 135.84, 135.10, 133.52 (d, J = 9.9 Hz), 132.60, 131.39, 130.52 (d, J = 12.4 Hz), 129.14, 128.79, 127.31, 124.61, 119.65, 19.24, 118.15 (d, J = 85.74 Hz), 117.95, 117.05, 113.65, 111.91, 40.17, 30.19, 30.06, 29.69, 29.00, 28.59, 28.50, 28.46, 26.61, 22.83, 22.51, 22.47.

HRMS calculated for C49H50O3N4P$^+$: 773.36150, found: 773.36128

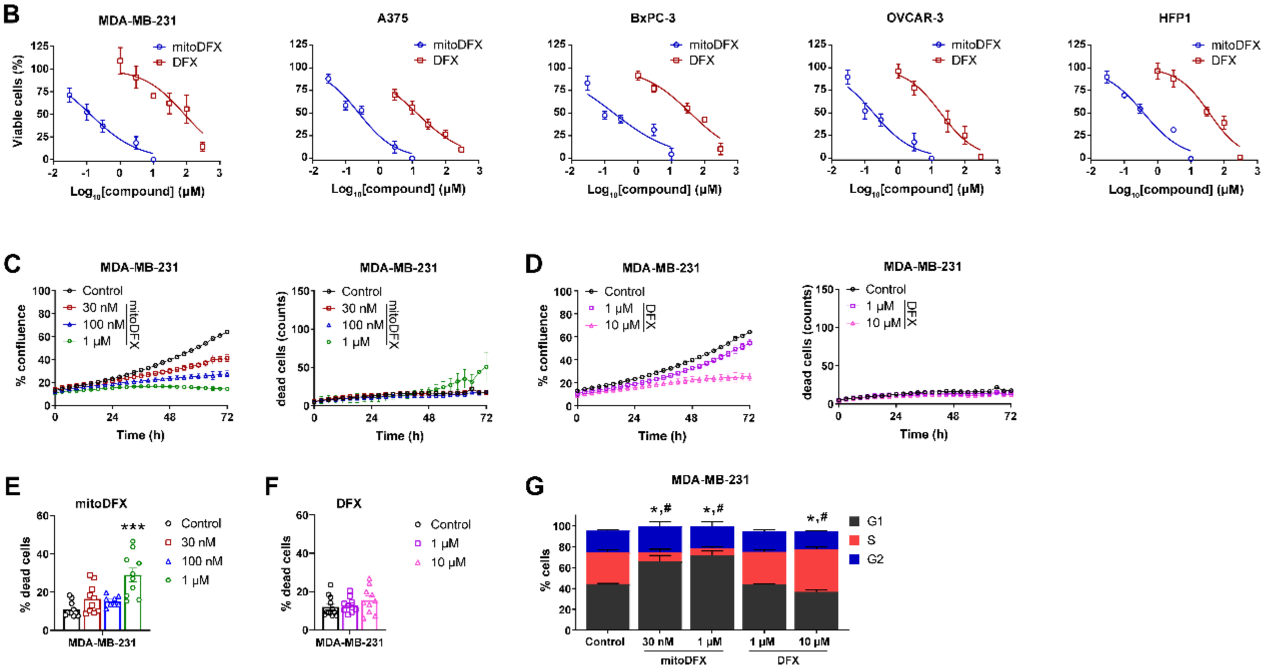


**Figure S1: mitoDFX treatment induces anti-proliferative effects in different cancer cell lines. (A**) Synthesis of (10-(4-(3,5-bis(2-hydroxyphenyl)-1H-1,2,4-triazol-1-yl)benzamido)decyl)triphenylphosphonium chloride (mitoDFX). **(B)** Five different cancer cell lines (MDA-MB-231, A375, BxPC-3, OVCAR-3, and HFP1) were treated with varying concentrations of mitoDFX and DFX for 48 h and cell viability was determined by crystal violet staining. **(C)** Proliferation and cell death of MDA-MB-231 exposed to mitoDFX using real-time LumaScope 720 microscope. Cell death was measured using Sytox green dye (0.5 µM). Proliferation and cell death **(D)** of MDA-MB-231 cells treated with DFX for 72 h using real-time LumaScope 720 microscope. Cell death was measured using Sytox green dye (0.5 µM). Breast cancer cell line MDA-MB-231 was treated with mitoDFX **(E)** and DFX **(F)**, stained with annexin V and propidium iodide, and flow cytometry was performed. **(G)** Cell cycle distribution was assessed in MCF7, MDA-MB-231 and BJ cells treated with indicated concentrations of mitoDFX and DFX and analyzed by FlowJo software. All data represent mean ± SEM of three independent experiments with at least two replicates each. P values were calculated by one-way **(E-F)** or two-way **(G)** ANOVA followed by Tukey's multiple comparisons test. *** $P < 0.001$ relative to the Control **(E-F)** and * $P < 0.05$ relative to the $G_1$ phase in the control; # $P < 0.05$ relative to the S phase in the control **(G)**.

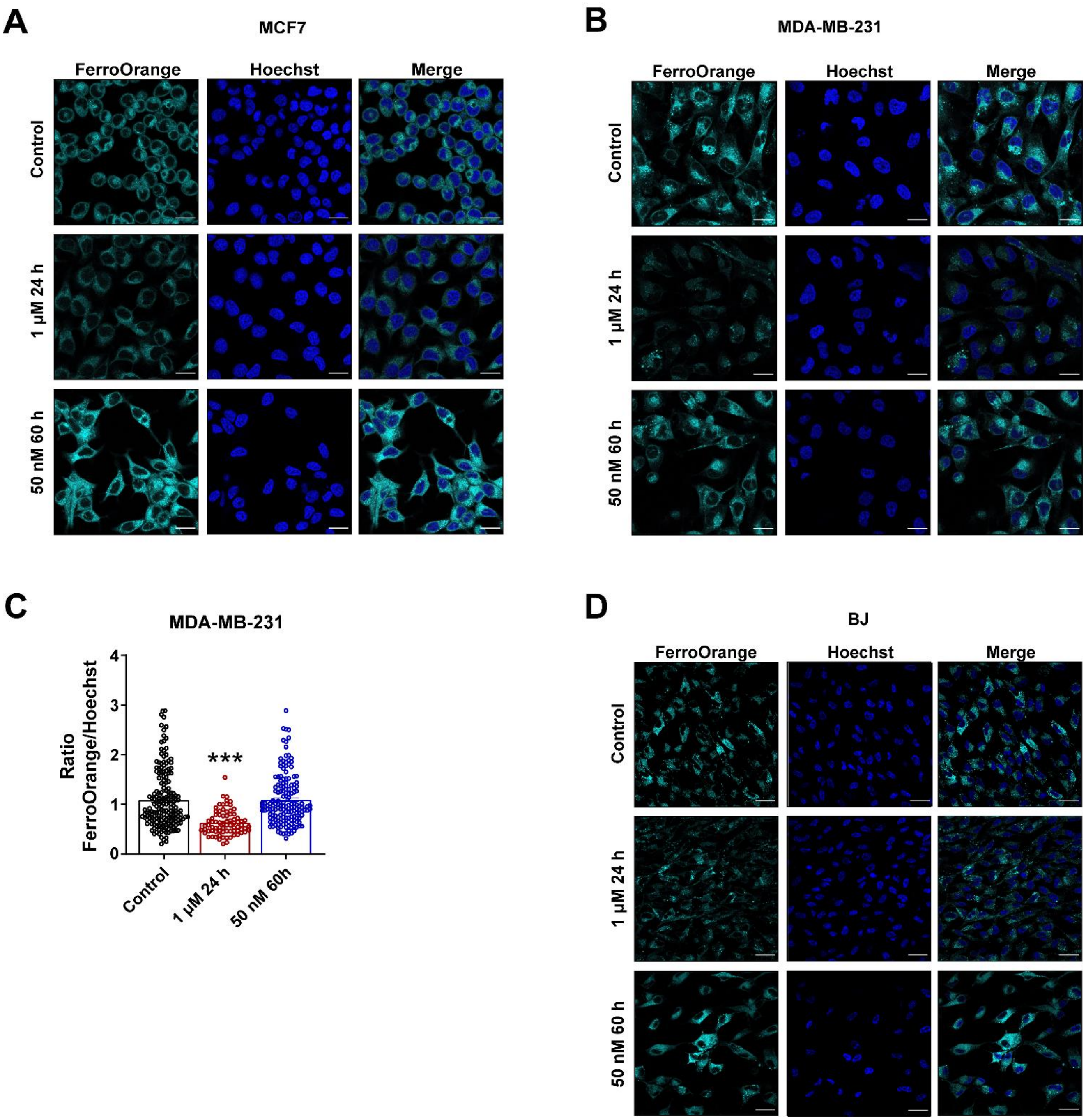


**Figure S2: mitoDFX reduces the levels of cellular ferrous iron at 1µM. (A, B, D)** Confocal images of MCF7, MDA-MB-231 and BJ cells treated with mitoDFX for short (24 h) and long term (60 h) and incubated with FerroOrange (1 µM) for 30 mins to assess the levels of cellular ferrous iron. Nuclei were stained with Hoechst 33342. **(C)** Quantification of the ratio of FerroOrange/Hoechst fluorescence relative to the control in MDA-MB-231 cells. All data represent mean ± SEM of three independent experiments with at least 50 cells each. P values were calculated by one-way ANOVA followed by Tukey's multiple comparisons test **(C)**. *** $P < 0.001$. Scale bars, 20 µm **(A, B, D)**.

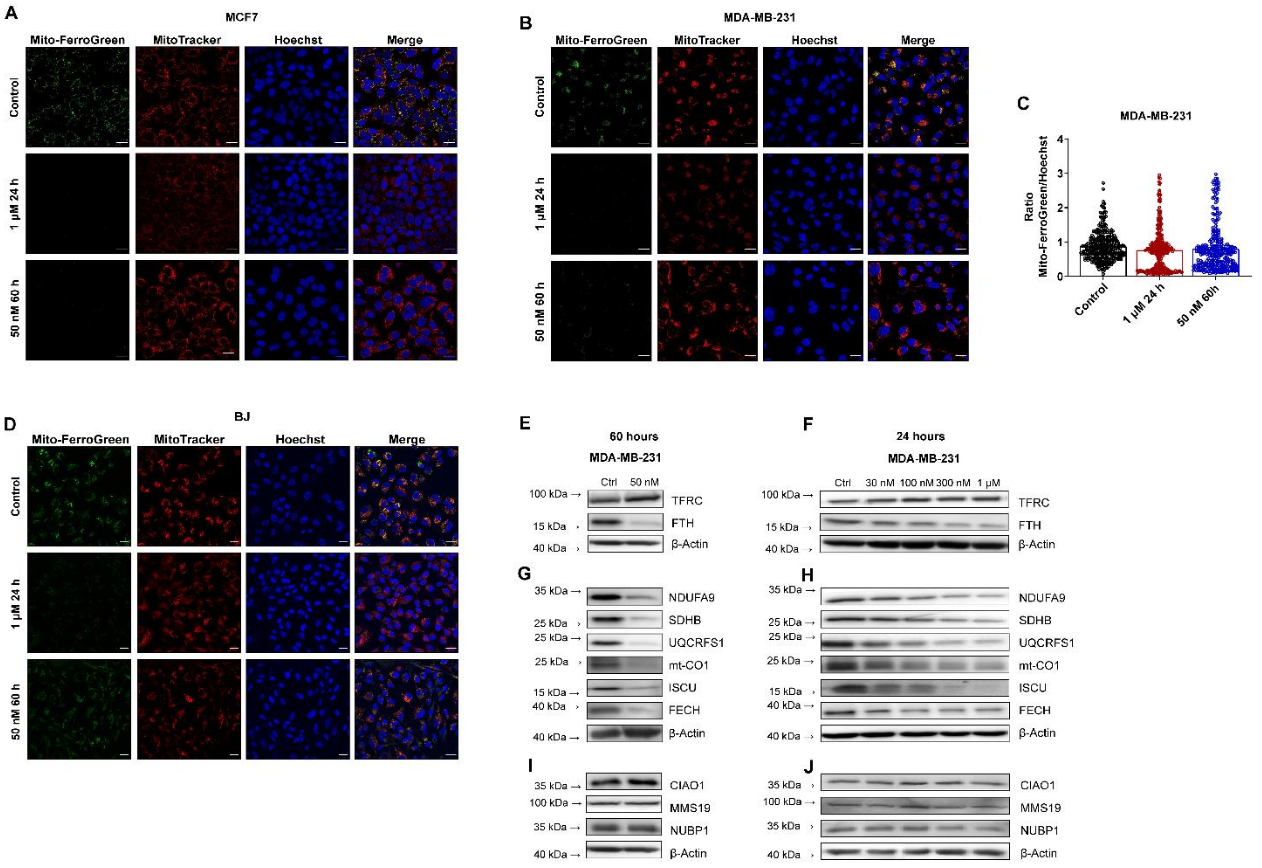


**Figure S3: mitoDFX reduces mitochondrial ferrous iron in malignant cells at 50 nM, induces iron deprivation and diminishes levels of [Fe-S] cluster and heme-containing proteins in breast cancer cells. (A, B, D)** Confocal images of MCF7, MDA-MB-231 and BJ cells treated with mitoDFX for short (24 h) and long term (60 h) and incubated with Mito-FerroGreen (5 μM). Nuclei were stained with Hoechst 33342. **(C)** Quantification of the ratio of Mito-FerroGreen/Hoechst fluorescence relative to control in MDA-MB-231 cells. **(E-F)** Western blot images of proteins related to iron metabolism in MDA-MB-231 cells treated with mitoDFX for long (60 h) and short term (24 h). **(G-H)** Western blot images of mitochondrial [Fe-S] cluster and heme-containing proteins metabolism in MDA-MB-231 cells treated with mitoDFX for long (60 h) and short term (24 h). **(I-J)** Western blot images of cytosolic [Fe-S] cluster biogenesis-related proteins obtained from MDA-MB-231 cells exposed to mitoDFX for long (60 h) and short term (24 h). All data represent mean ± SEM of three independent experiments with more than 50 cells each. P values were calculated by one-way ANOVA followed by Tukey's multiple comparisons test **(C)**. ****P* < 0.001 relative to Control. Scale bars = 20 μm **(A, B, D)**.

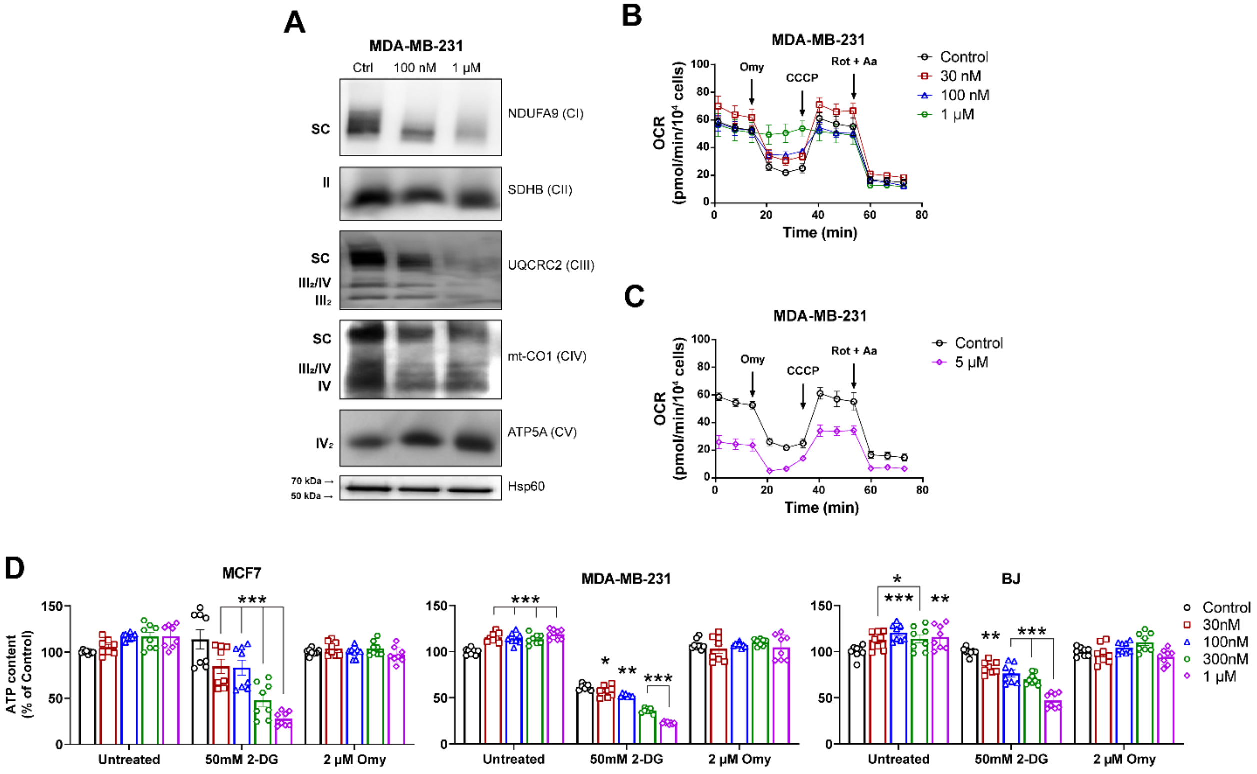


**Figure S4: Iron deficiency induced by mitoDFX disrupts mitochondrial respiratory chain complex assembly and affects mitochondrial respiratory activity. (A)** Mitochondrial lysates were isolated from MDA-MB-231 cells treated with mitoDFX for 24 h. BNE-PAGE was performed using the antibodies indicated. Oxygen consumption rate (OCR) profile of malignant MDA-MB-231 breast cancer cells treated with mitoDFX **(B)** or DFX **(C)** obtained using an extracellular flux analyzer (Seahorse XF). **(D)** The levels of intracellular ATP were determined in MCF7, MDA-MB-231, and BJ cells treated with varying concentrations of mitoDFX for 4 h in the presence or absence of 2-DG (50 mM) or Oligimycin (2 µM). The amounts of cellular ATP were calculated and normalized to control. All data represent three independent experiments with at least three replicates. *P* values were calculated by two-way ANOVA **(D)**. * $P < 0.05$, ** $P < 0.01$, *** $P < 0.001$ relative to Control.

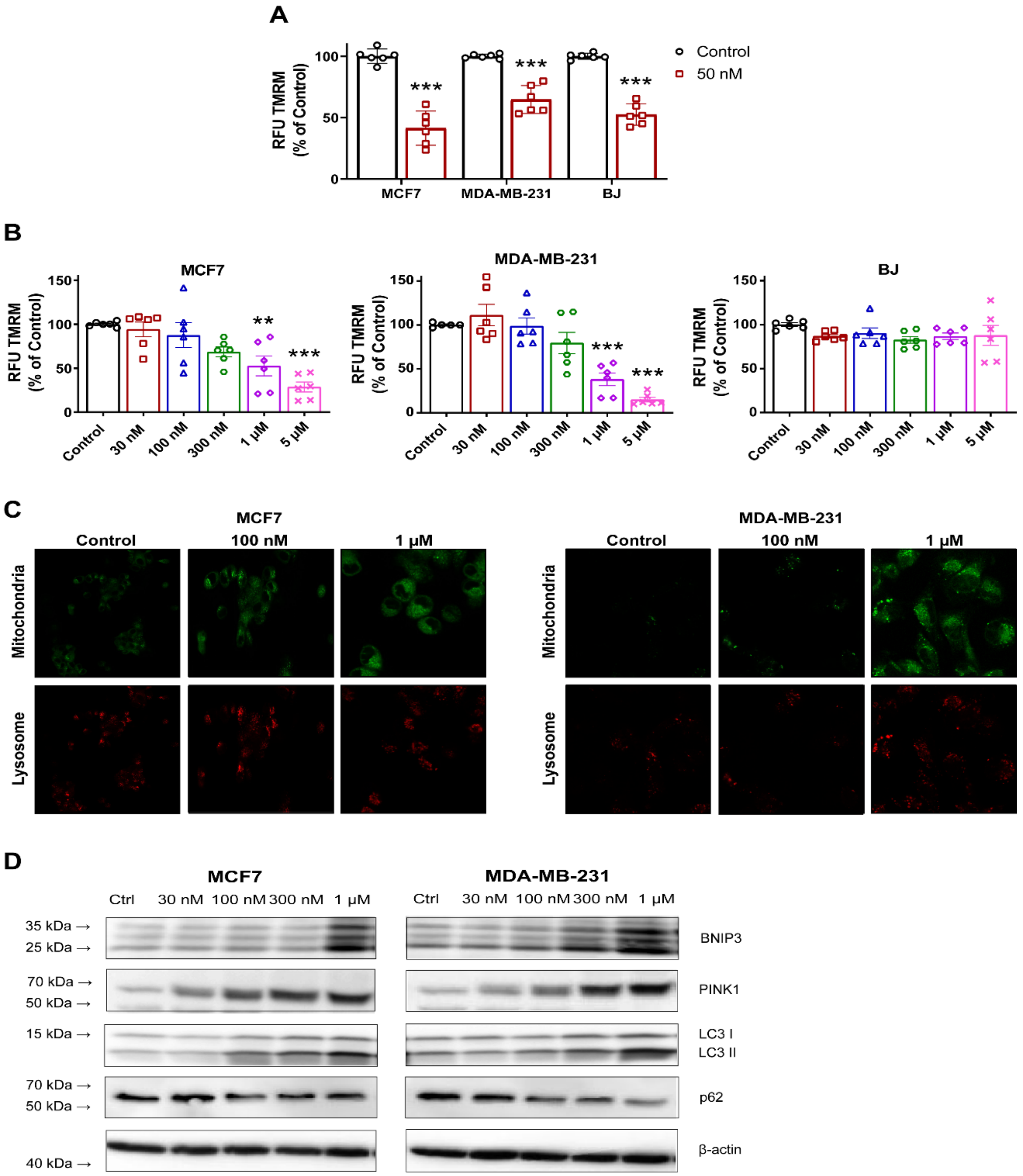


**Figure S5: MitoDFX induces mitochondrial dysfunction and mitophagy:** Quantification of mitochondrial membrane potential in MCF7, MDA-MDA-231 and BJ cells treated with mitoDFX for 60 h (50 nM) **(A)** or 24 h **(B)**. **(C)** Confocal microscopy images of breast cancer cell lines (MCF7 and MDA-MB-231) treated with mitoDFX for 24 h and stained with Mtphagy and Lyso dye for detecting mitochondrial acidification. **(D)** Western blot images of proteins relate to mitophagy induction in MCF7 and MDA-MB-231 breast cancer cells treated with varying concentrations of mitoDFX for 24 h. All data represent the mean ± SEM of three independent experiments with at least two replicates or 50 cells each. *P* values were calculated by t-student test **(A)** or by one-way ANOVA **(B)** followed by Tukey's multiple comparisons test, or by ** $P < 0.01$, *** $P < 0.001$ versus control **(A-B)**.

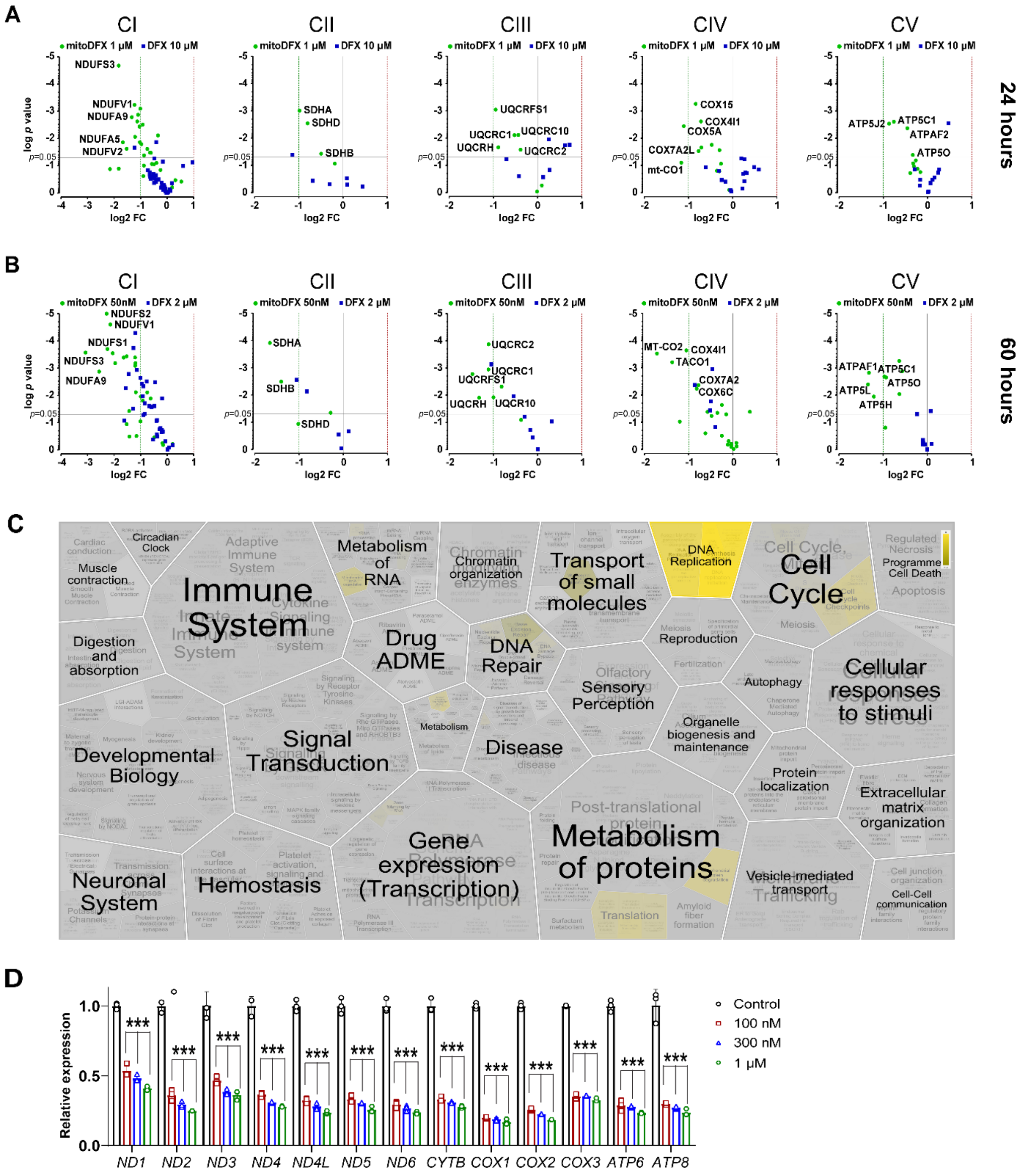


**Figure S6: MitoDFX affects mitochondrial transcription, replication and translation in breast cancer cells. (A-B)** Volcano plot showing proteins of the mitochondrial respiratory complexes detected by LC-MS in cells treated with either mitoDFX or DFX for 24-60 h. The blue points represent proteins after DFX treatment (p-value < 0.05, |fold change|> 2). The green points represent proteins after mitoDFX treatments (p-value < 0.05, |fold change|> 2. **(C)** Voronoi diagram depicting the results of the Reactome Pathway Analysis. **(D)** Real-time PCR analysis of mitochondrial transcripts in RNA samples obtained from MCF7 breast cancer cells after treatment with mitoDFX. Data was normalized to RPLP0. All data represent mean ± SEM of three independent experiments with at least three replicates each. *P* values were calculated by two-way ANOVA **(D)**. *** *P* < 0.001 relative to Control.

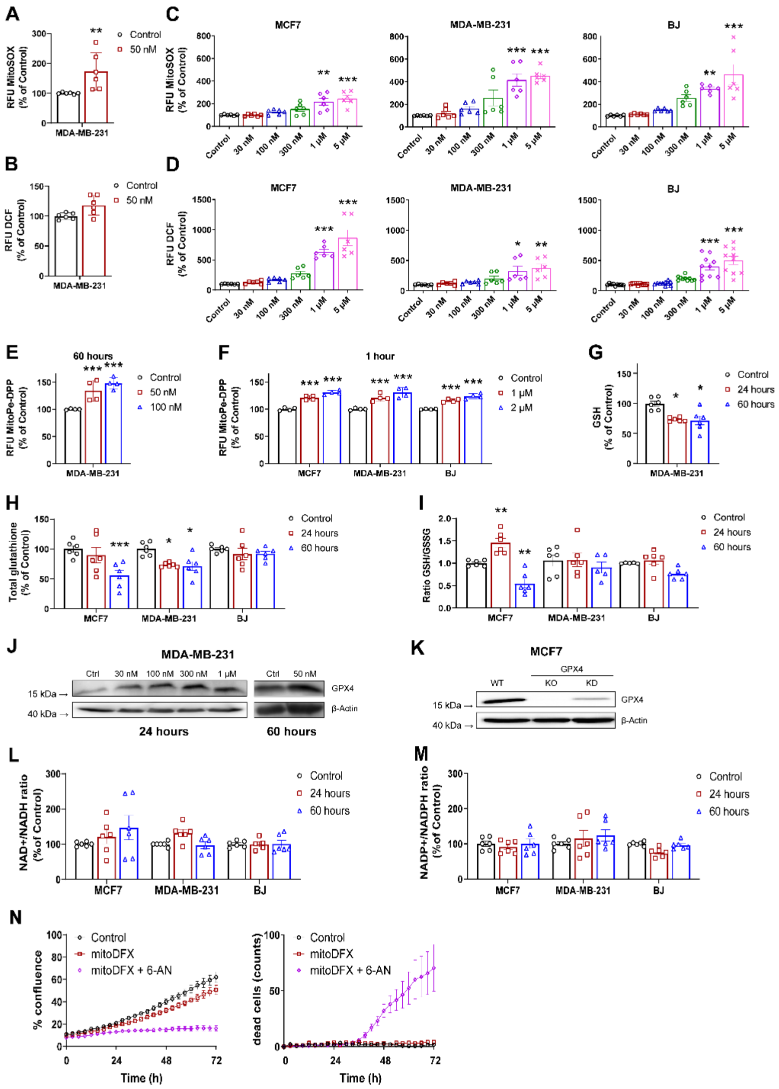


**Figure S7: Mitochondrial targeting induces oxidative stress and antioxidant defense mechanism as well as alters intracellular redox ratio.** Quantification of mitochondrial **(A)** and cellular ROS **(B)** of MDA-MB-231 cells treated with mitoDFX for 60 h (50 nM). Quantification of mitochondrial **(C)** and cellular ROS **(D)** of MCF7, MDA-MB-231 and BJ cells treated with mitoDFX for 24 h. Mitochondrial lipid peroxidation in MDA-MB-231 cells treated with mitoDFX for 60 h **(E)** and MCF7, MDA-MB-231 and BJ cells treated with mitoDFX for 1 h **(F)**. **(G)** Reduced levels of glutathione (GSH) were measured in MDA-MB-231 cells exposed to 50 nM mitoDFX for 24- 60 h. Total glutathione (GSH + GSSG) levels **(H)** and the ratio of GSH/GSSG **(I)** were determined in MCF7, MDA-MB-231 and BJ cells incubated with 50 nM mitoDFX for 24-60 h. **(J)** GPX4 protein levels were evaluated by western blot in MDA-MB-231 cells following treatment with varying concentrations of mitoDFX for 24-60 h. **(K)** Western blot analysis of GPX4 expression in WT and *GPX4* KO and KD MCF7 breast cancer cells. Ratio of cellular NAD$^+$/NADH **(L)** and NADP$^+$/NADPH **(M)** in MCF7, MDA-MB-231 and BJ cells treated with 50 nM mitoDFX for 24-60 h. **(N)** Proliferation assay of MCF7 cells treated with mitoDFX (10 nM) alone or in combination with 6-AN (10 µM) for 72 h. The growth curve was monitored using a real-time LumaScope 720 microscope and cell death was measured using Sytox green dye (0.5 µM). All data represent mean ± SEM of three independent experiments with at least two replicates each. *P* values were calculated by t-student test **(A-B)** or one-way ANOVA **(C-I)** followed by Tukey's multiple comparisons test. * $P < 0.05$, ** $P < 0.01$, *** $P < 0.001$ relative to Control.

# A

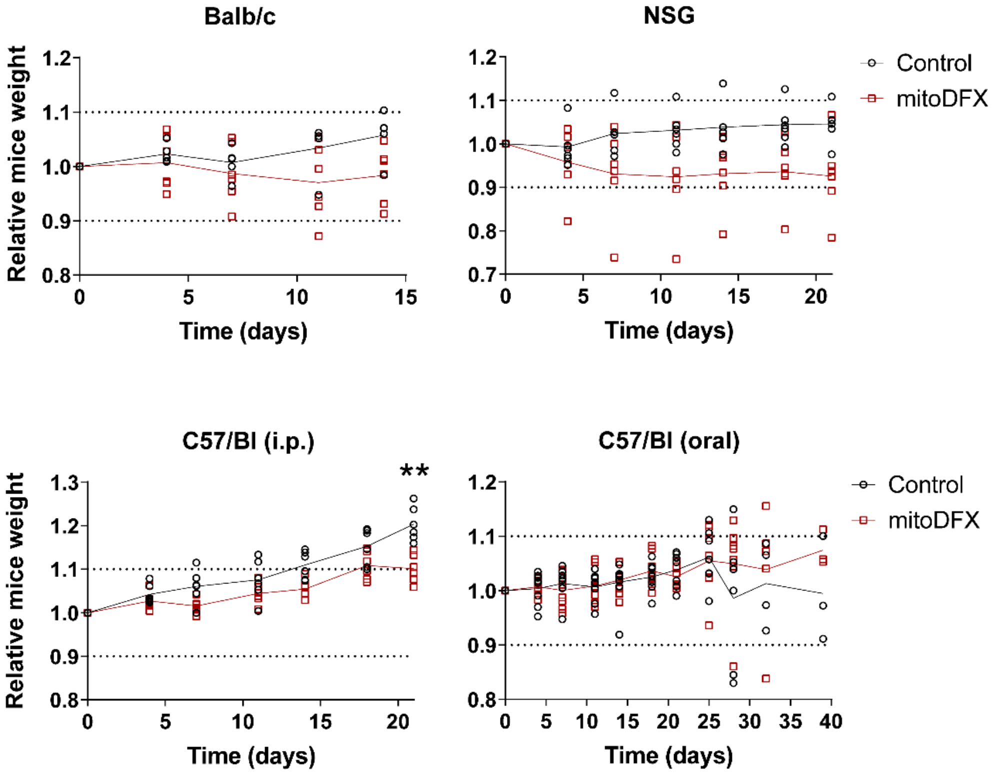


**Figure S8: mitoDFX does not affect mice body weight. (A)** Graph depicting the relative body weights of Balb/c, NSG, and C57Bl/6 tumor-bearing mice that were given mitoDFX i.p. or orally. Data represent mean ± SEM of at least 5 individual animals.

**Table S1. $IC_{50}$ values of iron-chelators in different cell lines.**

| $IC_{50}$ (μM) | mitoDFX | DFX |
|---|---|---|
| | **Breast cancer** | |
| **MCF7** | 0.01 ± 0.09 | 10.8 ± 0.3 |
| **MDA-MB-231** | 0.13 ± 0.04 | 85.6 ± 0.7 |
| **T47D** | 0.03 ± 0.04 | 59.0 ± 0.3 |
| | **Melanoma** | |
| **A375** | 0.35 ± 0.07 | 31.8 ± 0.7 |
| **B16** | 0.37 ± 0.09 | 23.9 ± 0.1 |
| **BLM** | 0.79 ± 0.05 | 274.3 ± 1.3 |
| **G361** | 0.19 ± 0.06 | 34.1 ± 0.1 |
| **Malme-3M** | 0.72 ± 0.08 | > 500 |
| | **Lung cancer** | |
| **A549** | 0.94 ± 0.09 | >500 |
| **Calu-1** | 1.11 ± 0.08 | > 500 |
| **LLC1** | 0.39 ± 0.07 | 23.3 ± 0.4 |
| | **Other cancers** | |
| **BxPC-3** | 0.19 ± 0.07 | 37.6 ± 0.5 |
| **OVCAR3** | 0.18 ± 0.05 | 18.6 ± 0.4 |
| | **Non-malignant cells** | |
| **BJ** | 0.90 ± 0.06 | 23.6 ± 0.6 |
| **HFP1** | 0.58 ± 0.05 | 35.9 ± 0.5 |

**Table S2. List of antibodies used for western blot experiments in SDS-PAGE and BNE gels throughout the paper.**

| | Target | Supplier | Catalog no. |
|---|---|---|---|
| SDS-PAGE | BNIP3 | Santa Cruz Biotechnology | sc-56167 |
| | CIAO1 | Santa Cruz Biotechnology | sc-374498 |
| | FECH | Santa Cruz Biotechnology | sc-377377 |
| | FPN | Bioss | bs-4906R |
| | FTH | Abcam | ab75973 |
| | GPX-4 | Abcam | ab 125066 |
| | ISCU | Abcam | ab180532 |
| | LC3B | Cell Signaling Technology | 3868S |
| | MMS19 | Santa Cruz Biotechnology | sc-390658 |
| | Mt-CO1 | Abcam | ab14705 |
| | NDUFA9 | Abcam | ab14713 |
| | NUBP1 | Santa Cruz Biotechnology | sc-398368 |
| | PINK1 | Santa Cruz Biotechnology | sc-517353 |
| | SDHB | Abcam | ab14714 |
| | TFRC | Life Technologies | 13-6800 |
| | SQSTM1/p62 | Santa Cruz Biotechnology | Sc-48402 |
| | UQCRFS1 | Abcam | ab14746 |
| | β-Actin | Santa Cruz Biotechnology | sc-47778 |
| | 4-HNE | ABclonal | A24456 |
| BNE | ATP5A (complex V) | Abcam | ab110273 |
| | HSP60 | Sigma | PLA0269 |
| | mtCO1 (complex IV) | Abcam | ab14705 |
| | NDUFA9 (complex I) | Abcam | ab14713 |
| | SDHB (complex II) | Abcam | ab14714 |
| | UQCRC2 (complex III) | Abcam | ab14745 |

**Table S3. List of primers and their sequences used for qPCR experiments throughout the paper.**

| Target | Supplier | Sequence |
|---|---|---|
| *MT-ND1* | Metabion | GGGCCTTTGCGTAGTTGTAT |
| | | ATACCCATGGCCAACCTCCT |
| *MT-ND2* | Metabion | GATGCGGTTGCTTGCGTGAG |
| | | GGCCCAACCCGTCATCTACT |
| *MT-ND3* | Metabion | CCGCGTCCCTTTCTCCATAAB |
| | | GGTAGGGGTAAAAGGAGGGC |
| *MT-ND4* | Metabion | ACTACTCACTCTCACTGCCC |
| | | AGTGGAGTCCGTAAAGAGGT |
| *MT-ND4L* | Metabion | TAGGCCCACCGCTGCTTCGC |
| | | AACCCTCAACACCCACTCCC |
| *MT-ND5* | Metabion | AACAGAGTGGTGATAGCGCC |
| | | CCCTACTCCACTCAAGCACT |
| *MT-ND6* | Metabion | AGGGGGAATGATGGTTGTCT |
| | | CCTACCTCCATCGCTAACCC |
| *MT-CO1* | Metabion | GCCTCCGTAGACCTAACCAT |
| | | GTTATGGCAGGGGGTTTTAT |
| *MT-CO2* | Metabion | AGTCCTGTATGCCCTTTTCC |
| | | GCGATGAGGACTAGGATGAT |
| *MT-CO3* | Metabion | CCCACCAATCACATCCTAT |
| | | TAGGCCGGAGGTCATTAGGA |
| *MT-CYB* | Metabion | GGCGATTGATGAAAAGGCGG |
| | | GAAACTTCGGCTCACTCCTT |
| *MT-ATP6* | Metabion | GATTAGTCATTGTTGGGTGG |
| | | CTGTTCGCTTCATTCATTGC |
| *MT-ATP8* | Metabion | CTTTGGTGAGGGAGGTAGGT |
| | | TGCCCCAACTAAATACTACC |
| *RPLP0* | Invitrogen | ATCACAGAGGAAACTCTGCATTCTCG |
| | | GATAGAATGGGGTACTGATGCAACAGTT |
