## Supplementary File1_Reactome for "Selective killing of cancer cells via simultaneous destabilization of mitochondrial iron metabolism and induction of ferroptosis elicited by mitochondrially targeted iron chelator deferasirox"

### Pathway Analysis Report

This report contains the pathway analysis results for the submitted sample ''. Analysis was performed against Reactome version 86 on 21/11/2023. The web link to these results is:

https://reactome.org/PathwayBrowser/#/ANALYSIS=MjAyMzExMTYxNzIyNTRfMjg1MjE%3D

Please keep in mind that analysis results are temporarily stored on our server. The storage period depends on usage of the service but is at least 7 days. As a result, please note that this URL is only valid for a limited time period and it might have expired.

### Table of Contents


1. Introduction
2. Properties
3. Genome-wide overview
4. Most significant pathways
5. Pathways details
6. Identifiers found
7. Identifiers not found

### 1. Introduction

Reactome is a curated database of pathways and reactions in human biology. Reactions can be considered as pathway 'steps'. Reactome defines a 'reaction' as any event in biology that changes the state of a biological molecule. Binding, activation, translocation, degradation and classical biochemical events involving a catalyst are all reactions. Information in the database is authored by expert biologists, entered and maintained by Reactome's team of curators and editorial staff. Reactome content frequently cross-references other resources e.g. NCBI, Ensembl, UniProt, KEGG (Gene and Compound), ChEBI, PubMed and GO. Orthologous reactions inferred from annotation for Homo sapiens are available for 14 non-human species including mouse, rat, chicken, puffer fish, worm, fly and yeast. Pathways are represented by simple diagrams following an SBGN-like format.

Reactome's annotated data describe reactions possible if all annotated proteins and small molecules were present and active simultaneously in a cell. By overlaying an experimental dataset on these annotations, a user can perform a pathway over-representation analysis. By overlaying quantitative expression data or time series, a user can visualize the extent of change in affected pathways and its progression. A binomial test is used to calculate the probability shown for each result, and the p-values are corrected for the multiple testing (Benjamini–Hochberg procedure) that arises from evaluating the submitted list of identifiers against every pathway.

To learn more about our Pathway Analysis, please have a look at our relevant publications:

Fabregat A, Sidiropoulos K, Garapati P, Gillespie M, Hausmann K, Haw R, … D'Eustachio P (2016). The reactome pathway knowledgebase. Nucleic Acids Research, 44(D1), D481–D487. https://doi.org/10.1093/nar/gkv1351.

Fabregat A, Sidiropoulos K, Viteri G, Forner O, Marin-Garcia P, Arnau V, … Hermjakob H (2017). Reactome pathway analysis: a high-performance in-memory approach. BMC Bioinformatics, 18.

### 2. Properties

- This is an **overrepresentation** analysis: A statistical (hypergeometric distribution) test that determines whether certain Reactome pathways are over-represented (enriched) in the submitted data. It answers the question 'Does my list contain more proteins for pathway X than would be expected by chance?' This test produces a probability score, which is corrected for false discovery rate using the Benjamani-Hochberg method.
- 533 out of 592 identifiers in the sample were found in Reactome, where 1765 pathways were hit by at least one of them.
- All non-human identifiers have been converted to their human equivalent.
- IntAct interactors were included to increase the analysis background. This greatly increases the size of Reactome pathways, which maximises the chances of matching your submitted identifiers to the expanded pathway, but will include interactors that have not undergone manual curation by Reactome and may include interactors that have no biological significance, or unexplained relevance.
- This report is filtered to show only results for species 'Homo sapiens' and resource 'all resources'.
- The unique ID for this analysis (token) is MjAyMzExMTYxNzIyNTRfMjg1MjE%3D. This ID is valid for at least 7 days in Reactome's server. Use it to access Reactome services with your data.

### 3. Genome-wide overview

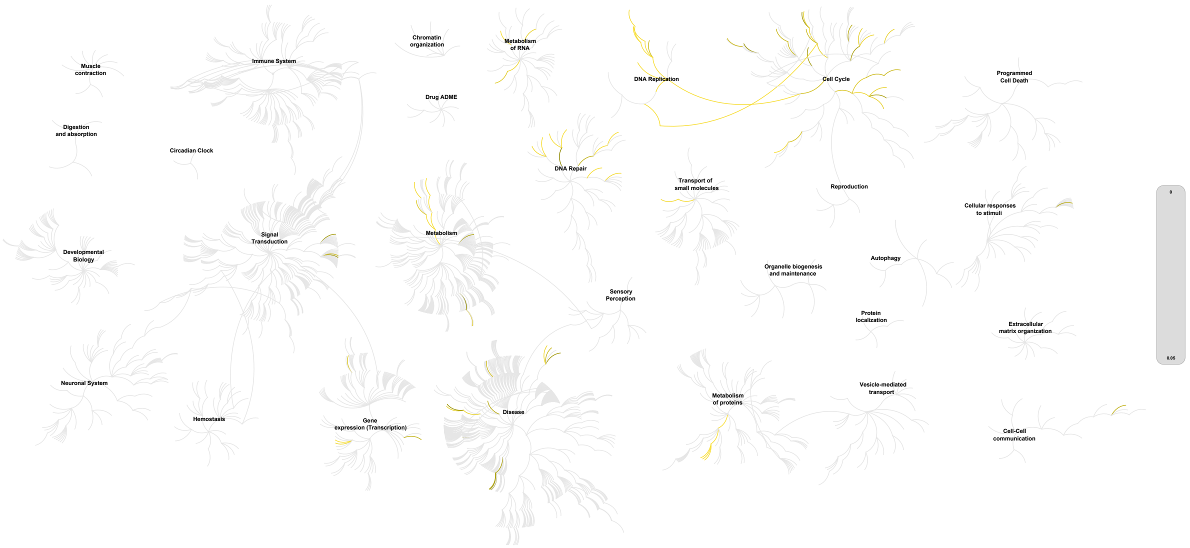


This figure shows a genome-wide overview of the results of your pathway analysis. Reactome pathways are arranged in a hierarchy. The center of each of the circular "bursts" is the root of one top-level pathway, for example "DNA Repair". Each step away from the center represents the next level lower in the pathway hierarchy. The color code denotes over-representation of that pathway in your input dataset. Light grey signifies pathways which are not significantly over-represented.

### 4. Most significant pathways

The following table shows the 25 most relevant pathways sorted by p-value.

| Pathway name | Entities | | | | Reactions | |
|---|---|---|---|---|---|---|
| | found | ratio | p-value | FDR* | found | ratio |
| Mitochondrial translation termination | 79 / 104 | 0.005 | 1.11e-16 | 4.23e-14 | 5 / 5 | 3.45e-04 |
| Mitochondrial translation | 86 / 199 | 0.009 | 1.11e-16 | 4.23e-14 | 13 / 17 | 0.001 |
| Mitochondrial translation initiation | 83 / 176 | 0.008 | 1.11e-16 | 4.23e-14 | 3 / 4 | 2.76e-04 |
| Mitochondrial translation elongation | 78 / 102 | 0.004 | 1.11e-16 | 4.23e-14 | 5 / 8 | 5.52e-04 |
| Translation | 114 / 732 | 0.032 | 1.11e-16 | 4.23e-14 | 57 / 99 | 0.007 |
| The citric acid (TCA) cycle and respiratory electron transport | 74 / 522 | 0.023 | 5.88e-13 | 1.86e-10 | 39 / 67 | 0.005 |
| Respiratory electron transport, ATP synthesis by chemiosmotic coupling, and heat production by uncoupling proteins. | 55 / 385 | 0.017 | 2.29e-11 | 6.24e-09 | 19 / 31 | 0.002 |
| Respiratory electron transport | 52 / 350 | 0.015 | 2.84e-11 | 6.75e-09 | 16 / 19 | 0.001 |
| Polymerase switching | 9 / 16 | 7.05e-04 | 7.18e-08 | 9.77e-06 | 4 / 4 | 2.76e-04 |
| Leading Strand Synthesis | 9 / 16 | 7.05e-04 | 7.18e-08 | 9.77e-06 | 4 / 4 | 2.76e-04 |
| DNA strand elongation | 19 / 77 | 0.003 | 7.34e-07 | 8.74e-05 | 14 / 15 | 0.001 |
| Lagging Strand Synthesis | 10 / 31 | 0.001 | 2.08e-06 | 2.18e-04 | 11 / 11 | 7.60e-04 |
| DNA replication initiation | 7 / 13 | 5.73e-04 | 2.71e-06 | 2.58e-04 | 2 / 2 | 1.38e-04 |
| Complex I biogenesis | 37 / 287 | 0.013 | 4.41e-06 | 3.97e-04 | 12 / 13 | 8.98e-04 |
| Gap-filling DNA repair synthesis and ligation in GG-NER | 9 / 27 | 0.001 | 5.15e-06 | 4.22e-04 | 2 / 2 | 1.38e-04 |
| Synthesis of DNA | 35 / 209 | 0.009 | 5.35e-06 | 4.22e-04 | 25 / 27 | 0.002 |
| Gap-filling DNA repair synthesis and ligation in TC-NER | 13 / 66 | 0.003 | 1.37e-05 | 0.001 | 2 / 2 | 1.38e-04 |
| Activation of the pre-replicative complex | 21 / 67 | 0.003 | 1.59e-05 | 0.001 | 9 / 9 | 6.21e-04 |
| Resolution of AP sites via the multiple-nucleotide patch replacement pathway | 11 / 55 | 0.002 | 5.38e-05 | 0.004 | 12 / 19 | 0.001 |
| PCNA-Dependent Long Patch Base Excision Repair | 9 / 39 | 0.002 | 8.77e-05 | 0.006 | 6 / 6 | 4.14e-04 |
| Processive synthesis on the lagging strand | 7 / 24 | 0.001 | 1.30e-04 | 0.008 | 7 / 7 | 4.83e-04 |
| tRNA processing in the mitochondrion | 9 / 45 | 0.002 | 2.51e-04 | 0.015 | 2 / 3 | 2.07e-04 |
| Polymerase switching on the C-strand of the telomere | 11 / 67 | 0.003 | 2.92e-04 | 0.017 | 6 / 6 | 4.14e-04 |

| Pathway name | Entities | | | | Reactions | |
|---|---|---|---|---|---|---|
| | found | ratio | p-value | FDR* | found | ratio |
| Recognition of DNA damage by PCNA-containing replication complex | 8 / 40 | 0.002 | 5.46e-04 | 0.031 | 5 / 6 | 4.14e-04 |
| Dual incision in TC-NER | 14 / 86 | 0.004 | 6.61e-04 | 0.035 | 7 / 7 | 4.83e-04 |

* False Discovery Rate

### 5. Pathways details

For every pathway of the most significant pathways, we present its diagram, as well as a short summary, its bibliography and the list of inputs found in it.

#### 1. Mitochondrial translation termination (R-HSA-5419276)

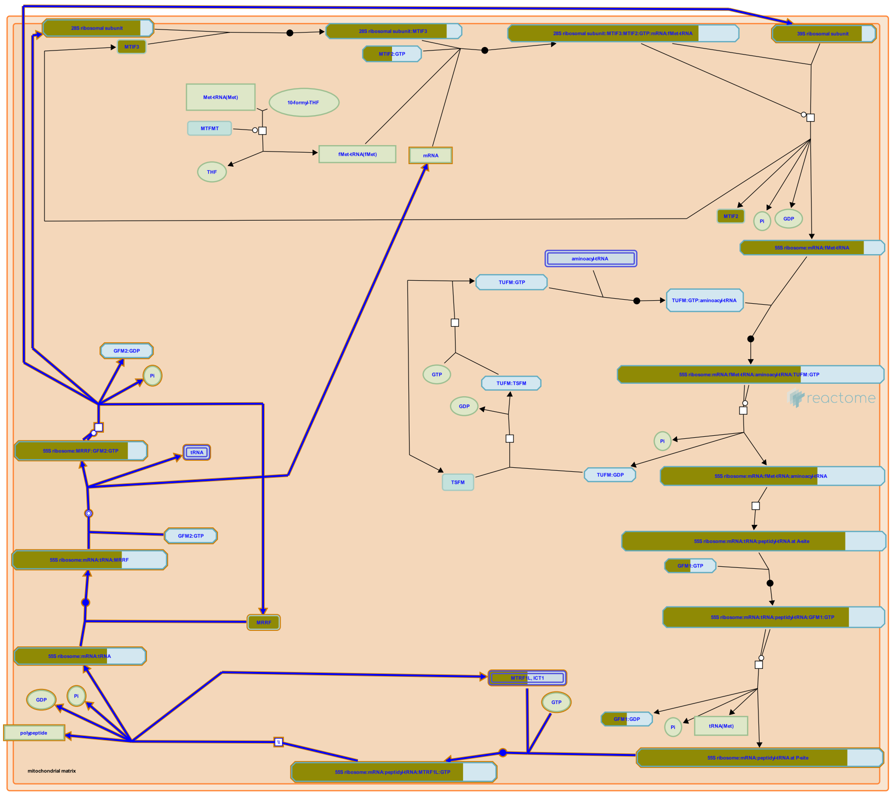

**Cellular compartments:** mitochondrial inner membrane, mitochondrial matrix.

Translation is terminated when MTRF1L:GTP (MTRF1a:GTP) recognizes a UAA or UAG termination codon in the mRNA at the A site of the ribosome (Soleimanpour-Lichaei et al. 2007, reviewed in Richter et al. 2010, Chrzanowska-Lightowlers et al. 2011. Christian and Spremulli 2012). GTP is hydrolyzed, and the aminoacyl bond between the translated polypeptide and the final tRNA at the P site is hydrolyzed by the 39S ribosomal subunit, releasing the translated polypeptide. MRRF (RRF) and GFM2:GTP (EF-G2mt:GTP) then act to release the remaining tRNA and mRNA from the ribosome and dissociate the 55S ribosome into 28S and 39S subunits.

#### Edit history

| Date | Action | Author |
|---|---|---|
| 2014-04-30 | Edited | May B |
| 2014-04-30 | Authored | May B |
| 2014-05-03 | Created | May B |
| 2014-08-29 | Reviewed | Chrzanowska-Lightowlers ZM |
| 2014-09-20 | Reviewed | Spremulli LL |
| 2023-08-26 | Modified | Wright A |

#### 77 submitted entities found in this pathway, mapping to 84 Reactome entities

| Input | UniProt Id | Input | UniProt Id | Input | UniProt Id |
|---|---|---|---|---|---|
| BRIP1 | Q9P0J6 | DAP3 | P51398 | ERAL1 | O75616 |
| GADD45GIP1 | Q8TAE8 | ICT1 | Q14197 | MRPL1 | Q9BYD6 |
| MRPL10 | Q7Z7H8 | MRPL11 | Q9Y3B7 | MRPL12 | P52815 |
| MRPL13 | Q9BYD1 | MRPL14 | Q6P1L8 | MRPL15 | P49406, Q9P015 |
| MRPL16 | Q9NX20 | MRPL17 | Q9NRX2 | MRPL18 | Q9H0U6 |
| MRPL19 | P49406 | MRPL2 | Q5T653, Q9BZE1 | MRPL20 | Q9BYC9 |
| MRPL21 | Q7Z2W9 | MRPL22 | Q9NWU5 | MRPL23 | Q16540 |
| MRPL24 | Q96A35 | MRPL27 | Q8IXM3, Q9P0M9 | MRPL28 | Q13084, Q8TCC3 |
| MRPL3 | P09001 | MRPL32 | Q6P1L8, Q9BYC8 | MRPL34 | Q9BQ48 |
| MRPL37 | Q9BZE1 | MRPL38 | Q96DV4 | MRPL39 | Q9NYK5 |
| MRPL4 | Q9BYD3 | MRPL40 | Q9NQ50 | MRPL41 | Q8IXM3 |
| MRPL42 | Q9Y6G3 | MRPL44 | Q9H9J2 | MRPL45 | Q9BRJ2 |
| MRPL46 | Q9H2W6 | MRPL47 | Q9HD33 | MRPL48 | Q96GC5 |
| MRPL49 | Q13405 | MRPL51 | Q4U2R6 | MRPL52 | Q86TS9 |
| MRPL53 | Q96EL3 | MRPL54 | Q6P161 | MRPL55 | Q7Z7F7 |
| MRPL9 | Q9BYD2 | MRPS10 | P82664 | MRPS11 | P82912 |
| MRPS12 | O15235 | MRPS14 | O60783 | MRPS15 | P82914 |
| MRPS16 | Q9Y3D3 | MRPS17 | Q9Y2R5 | MRPS18A | Q9NVS2 |
| MRPS18B | Q9Y676 | MRPS18C | Q9Y3D5 | MRPS2 | Q9Y399 |
| MRPS21 | P82921 | MRPS22 | P82650 | MRPS23 | Q9Y3D9 |
| MRPS24 | Q96EL2 | MRPS25 | P82663 | MRPS26 | Q9BYN8 |
| MRPS27 | Q92552 | MRPS28 | P82673, Q9Y2Q9 | MRPS30 | Q9NP92 |
| MRPS31 | Q92665 | MRPS33 | Q9Y291 | MRPS34 | P82930 |
| MRPS35 | P82673, Q9Y2Q9 | MRPS5 | P82675 | MRPS6 | P82932 |
| MRPS7 | Q9Y2R9 | MRPS9 | P82933 | MRRF | Q96E11 |
| OXA1L | Q15070 | PTCD3 | Q96EY7 | | |

#### Interactors found in this pathway (1)

| Input | UniProt Id | Interacts with | Input | UniProt Id | Interacts with |
|---|---|---|---|---|---|
| TRIM27 | P14373 | Q96E11 | | | |

#### 2. Mitochondrial translation (R-HSA-5368287)

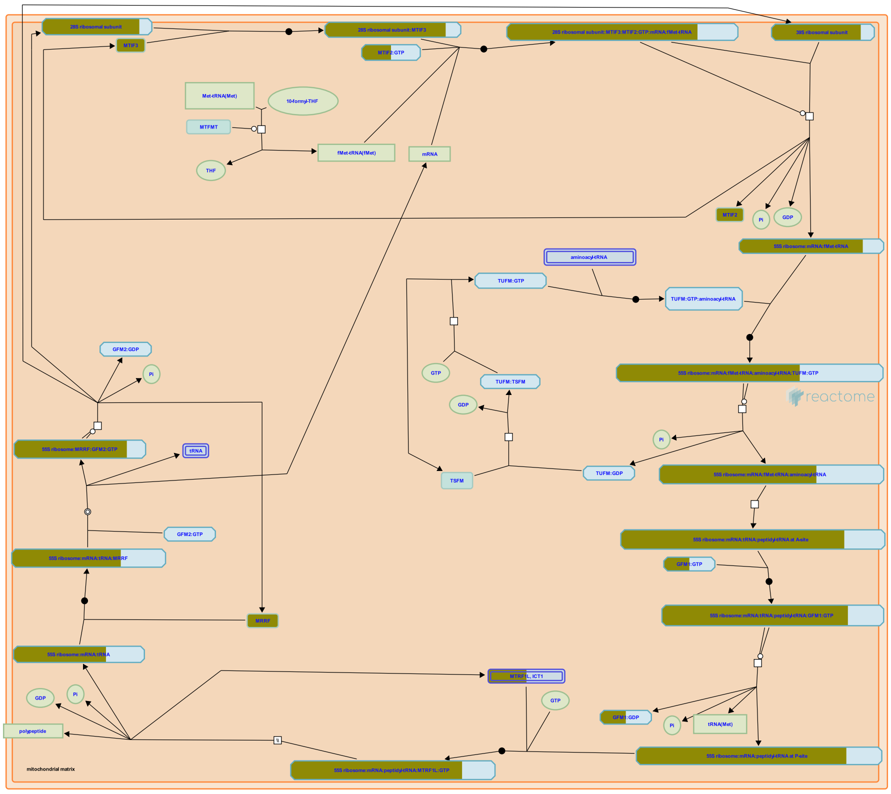

**Cellular compartments:** mitochondrial inner membrane, mitochondrial matrix.

Of the roughly 1000 human mitochondrial proteins only 13 proteins, all of them hydrophobic inner membrane proteins that are components of the oxidative phosphorylation apparatus, are encoded in the mitochondrial genome and translated by mitoribosomes at the matrix face of the inner membrane (reviewed in Herrmann et al. 2012, Hallberg and Larsson 2014, Lightowlers et al. 2014). The remainder, including all proteins of the mitochondrial translation system, are encoded in the nucleus and imported from the cytosol into the mitochondrion. Translation in the mitochondrion reflects both the bacterial origin of the organelle and subsequent divergent evolution during symbiosis (reviewed in Huot et al. 2014, Richman et al. 2014). Human mitochondrial ribosomes have a low sedimentation coefficient of only 55S, but at 2.71 MDa they retain a similar mass to E. coli 70S particles. The 55S particles are protein-rich compared to both cytosolic ribosomes and eubacterial ribosomes. This is due to shorter mt-rRNAs, mitochondria-specific proteins, and numerous rearrangements in individual protein positions within the two ribosome subunits (inferred from bovine ribosomes in Sharma et al. 2003, Greber et al. 2014, Kaushal et al. 2014, reviewed in Agrawal and Sharma 2012).

Mitochondrial mRNAs have either no untranslated leader or short leaders of 1-3 nucleotides, with the exception of the 2 bicistronic transcripts, RNA7 and RNA14, which have overlapping orfs that encode ND4L/ND4 and ATP8/ATP6 respectively. Translation is believed to initiate with the mRNA binding the 28S subunit:MTIF3 (28S subunit:IF-3Mt, 28S subunit:IF2mt) complex together with MTIF2:GTP (IF-2Mt:GTP, IF2mt:GTP) at the matrix face of the inner membrane (reviewed in Christian and Spremulli 2012). MTIF3 can dissociate 55S particles in preparation for initiation, enhances formation of initiation complexes, and inhibits N-formylmethionine-tRNA (fMet-tRNA) binding to 28S subunits in the absence of mRNA. Binding of fMet-tRNA to the start codon of the mRNA results in a stable complex while absence of a start codon at the 5' end of the mRNA causes eventual dissociation of the mRNA from the 28S subunit. After recognition of a start codon, the 39S subunit then binds the stable complex, GTP is hydrolyzed, and the initiation factors MTIF3 and MTIF2:GDP dissociate.

Translation elongation then proceeds by cycles of aminoacyl-tRNAs binding, peptide bond formation, and displacement of deacylated tRNAs. In each cycle an aminoacyl-tRNA in a complex with TUFM:GTP (EF-Tu:GTP) binds at the A-site of the ribosome, GTP is hydrolyzed, and TUFM:GDP dissociates. The elongating polypeptide bonded to the tRNA at the P-site is transferred to the aminoacyl group at the A-site by peptide bond formation at the peptidyl transferase center, leaving a deacylated tRNA at the P-site and the elongating polypeptide attached to the tRNA at the A-site. The polypeptide is co-translationally inserted into the inner mitochondrial membrane via an interaction with OXA1L (Haque et al. 2010, reviewed in Ott and Hermann 2010). After peptide bond formation, GFM1:GTP (EF-Gmt:GTP) then binds the ribosome complex, GTP is hydrolyzed, GFM1:GDP dissociates, and the ribosome translocates 3 nucleotides in the 3' direction along the mRNA, relocating the polypeptide-tRNA to the P-site and allowing another cycle to begin. TUFM:GDP is regenerated to TUFM:GTP by the guanine nucleotide exchange factor TSFM (EF-Ts, EF-TsMt).

Translation is terminated when MTRF1L:GTP (MTRF1a:GTP) recognizes an UAA or UAG termination codon at the A-site of the ribosome (Tsuboi et al. 2009). GTP hydrolysis does not appear to be required. The tRNA-aminoacyl bond between the translated polypeptide and the final tRNA at the P-site is hydrolyzed by the 39S subunit, facilitating release of the polypeptide. MRRF (RRF) and GFM2:GTP (EF-G2mt:GTP) then act to release the remaining tRNA and mRNA from the ribosome and dissociate the 55S ribosome into 28S and 39S subunits.

Mutations have been identified in genes encoding mitochondrial ribosomal proteins and translation factors. These have been shown to be pathogenic, causing neurological and other diseases (reviewed in Koopman et al. 2013, Pearce et al. 2013).

#### Edit history

| Date | Action | Author |
|---|---|---|
| 2014-04-26 | Edited | May B |
| 2014-04-26 | Authored | May B |
| 2014-04-26 | Created | May B |
| 2014-08-29 | Reviewed | Chrzanowska-Lightowlers ZM |
| 2014-09-20 | Reviewed | Spremulli LL |
| 2023-08-26 | Modified | Wright A |

#### 80 submitted entities found in this pathway, mapping to 87 Reactome entities

| Input | UniProt Id | Input | UniProt Id | Input | UniProt Id |
|---|---|---|---|---|---|
| BRIP1 | Q9P0J6 | DAP3 | P51398 | ERAL1 | O75616 |
| GADD45GIP1 | Q8TAE8 | GFM1 | Q96RP9 | ICT1 | Q14197 |
| MRPL1 | Q9BYD6 | MRPL10 | Q7Z7H8 | MRPL11 | Q9Y3B7 |
| MRPL12 | P52815 | MRPL13 | Q9BYD1 | MRPL14 | Q6P1L8 |
| MRPL15 | P49406, Q9P015 | MRPL16 | Q9NX20 | MRPL17 | Q9NRX2 |
| MRPL18 | Q9H0U6 | MRPL19 | P49406 | MRPL2 | Q5T653, Q9BZE1 |
| MRPL20 | Q9BYC9 | MRPL21 | Q7Z2W9 | MRPL22 | Q9NWU5 |
| MRPL23 | Q16540 | MRPL24 | Q96A35 | MRPL27 | Q8IXM3, Q9P0M9 |
| MRPL28 | Q13084, Q8TCC3 | MRPL3 | P09001 | MRPL32 | Q6P1L8, Q9BYC8 |
| MRPL34 | Q9BQ48 | MRPL37 | Q9BZE1 | MRPL38 | Q96DV4 |
| MRPL39 | Q9NYK5 | MRPL4 | Q9BYD3 | MRPL40 | Q9NQ50 |
| MRPL41 | Q8IXM3 | MRPL42 | Q9Y6G3 | MRPL44 | Q9H9J2 |
| MRPL45 | Q9BRJ2 | MRPL46 | Q9H2W6 | MRPL47 | Q9HD33 |
| MRPL48 | Q96GC5 | MRPL49 | Q13405 | MRPL51 | Q4U2R6 |
| MRPL52 | Q86TS9 | MRPL53 | Q96EL3 | MRPL54 | Q6P161 |
| MRPL55 | Q7Z7F7 | MRPL9 | Q9BYD2 | MRPS10 | P82664 |
| MRPS11 | P82912 | MRPS12 | O15235 | MRPS14 | O60783 |
| MRPS15 | P82914 | MRPS16 | Q9Y3D3 | MRPS17 | Q9Y2R5 |
| MRPS18A | Q9NVS2 | MRPS18B | Q9Y676 | MRPS18C | Q9Y3D5 |
| MRPS2 | Q9Y399 | MRPS21 | P82921 | MRPS22 | P82650 |
| MRPS23 | Q9Y3D9 | MRPS24 | Q96EL2 | MRPS25 | P82663 |
| MRPS26 | Q9BYN8 | MRPS27 | Q92552 | MRPS28 | P82673, Q9Y2Q9 |
| MRPS30 | Q9NP92 | MRPS31 | Q92665 | MRPS33 | Q9Y291 |
| MRPS34 | P82930 | MRPS35 | P82673, Q9Y2Q9 | MRPS5 | P82675 |
| MRPS6 | P82932 | MRPS7 | Q9Y2R9 | MRPS9 | P82933 |
| MRRF | Q96E11 | MTIF2 | P46199 | MTIF3 | Q9H2K0 |
| OXA1L | Q15070 | PTCD3 | Q96EY7 | | |

#### Interactors found in this pathway (5)

| Input | UniProt Id | Interacts with |
| --- | --- | --- |
| CD47 | Q08722-3 | Q9H2K0 |
| LDLR | P01130 | Q9H2K0 |
| TRIM27 | P14373 | Q96E11 |
| FXYD3 | Q14802-3 | Q9H2K0 |
| MRFAP1 | Q9Y605 | Q9H2K0 |

#### 3. Mitochondrial translation initiation (R-HSA-5368286)

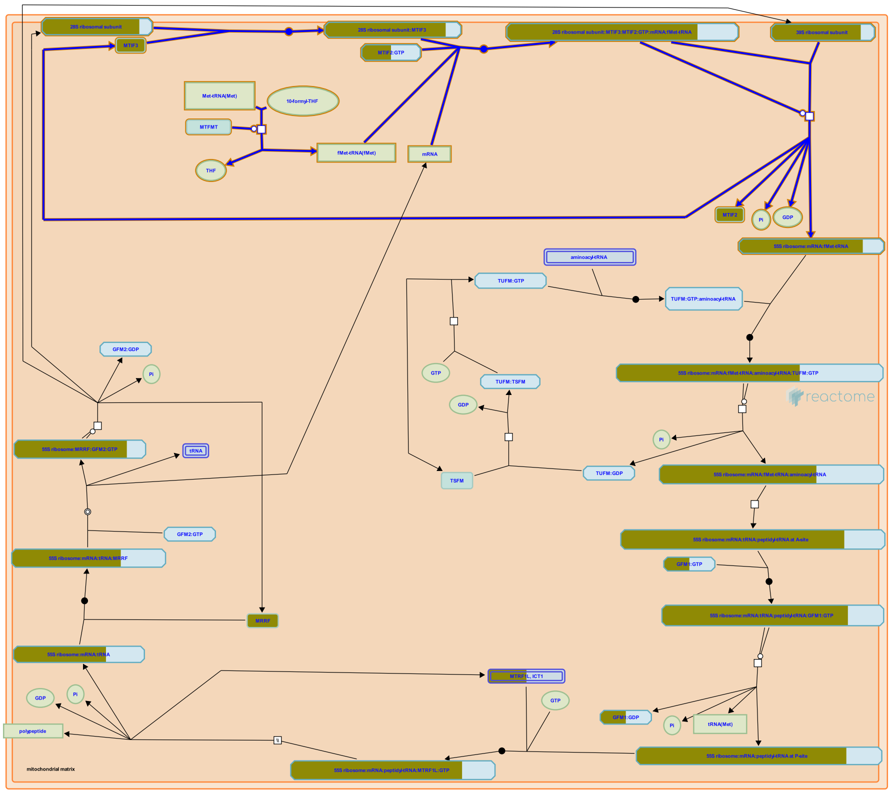

**Cellular compartments:** mitochondrial inner membrane, mitochondrial matrix.

Translation initiates with the mitochondrial mRNA binding the 28S subunit:MTIF3 (28S subunit:IF-3Mt, 28S subunit:IF3mt) complex together with MTIF2:GTP (IF-2Mt:GTP, IF2mt:GTP) (reviewed in Christian and Spremulli 2012, Kuzmenko et al. 2014). As inferred from bovine homologs, the 28S subunit, 39S subunit, and 55S holoribosome associate with the matrix-side face of the inner membrane and the translation products are inserted into the inner membrane as translation occurs (Liu and Spremulli 2000). Mitochondrial mRNAs have either no untranslated leader or short leaders of 1-3 nucleotides, with the exception of the 2 bicistronic transcripts, RNA7 and RNA14, which have overlapping orfs that encode ND4L/ND4 and ATP8/ATP6 respectively.. Binding of N-formylmethionine-tRNA to the start codon results in a stable complex between the mRNA and the 28S subunit while absence of a start codon at the 5' end of the mRNA causes the mRNA to slide though the 28S subunit and eventually dissociate. The 39S subunit then binds the 28S subunit:mRNA complex, GTP is hydrolyzed, and the initiation factors MTIF3 and MTIF2:GDP dissociate.


##### Edit history

| Date | Action | Author |
|---|---|---|
| 2014-04-26 | Edited | May B |
| 2014-04-26 | Authored | May B |
| 2014-04-26 | Created | May B |
| 2014-08-29 | Reviewed | Chrzanowska-Lightowlers ZM |
| 2014-09-20 | Reviewed | Spremulli LL |
| 2023-03-08 | Modified | Matthews L |

#### 78 submitted entities found in this pathway, mapping to 85 Reactome entities

| Input | UniProt Id | Input | UniProt Id | Input | UniProt Id |
|---|---|---|---|---|---|
| BRIP1 | Q9P0J6 | DAP3 | P51398 | ERAL1 | O75616 |
| GADD45GIP1 | Q8TAE8 | ICT1 | Q14197 | MRPL1 | Q9BYD6 |
| MRPL10 | Q7Z7H8 | MRPL11 | Q9Y3B7 | MRPL12 | P52815 |
| MRPL13 | Q9BYD1 | MRPL14 | Q6P1L8 | MRPL15 | P49406, Q9P015 |
| MRPL16 | Q9NX20 | MRPL17 | Q9NRX2 | MRPL18 | Q9H0U6 |
| MRPL19 | P49406 | MRPL2 | Q5T653, Q9BZE1 | MRPL20 | Q9BYC9 |
| MRPL21 | Q7Z2W9 | MRPL22 | Q9NWU5 | MRPL23 | Q16540 |
| MRPL24 | Q96A35 | MRPL27 | Q8IXM3, Q9P0M9 | MRPL28 | Q13084, Q8TCC3 |
| MRPL3 | P09001 | MRPL32 | Q6P1L8, Q9BYC8 | MRPL34 | Q9BQ48 |
| MRPL37 | Q9BZE1 | MRPL38 | Q96DV4 | MRPL39 | Q9NYK5 |
| MRPL4 | Q9BYD3 | MRPL40 | Q9NQ50 | MRPL41 | Q8IXM3 |
| MRPL42 | Q9Y6G3 | MRPL44 | Q9H9J2 | MRPL45 | Q9BRJ2 |
| MRPL46 | Q9H2W6 | MRPL47 | Q9HD33 | MRPL48 | Q96GC5 |
| MRPL49 | Q13405 | MRPL51 | Q4U2R6 | MRPL52 | Q86TS9 |
| MRPL53 | Q96EL3 | MRPL54 | Q6P161 | MRPL55 | Q7Z7F7 |
| MRPL9 | Q9BYD2 | MRPS10 | P82664 | MRPS11 | P82912 |
| MRPS12 | O15235 | MRPS14 | O60783 | MRPS15 | P82914 |
| MRPS16 | Q9Y3D3 | MRPS17 | Q9Y2R5 | MRPS18A | Q9NVS2 |
| MRPS18B | Q9Y676 | MRPS18C | Q9Y3D5 | MRPS2 | Q9Y399 |
| MRPS21 | P82921 | MRPS22 | P82650 | MRPS23 | Q9Y3D9 |
| MRPS24 | Q96EL2 | MRPS25 | P82663 | MRPS26 | Q9BYN8 |
| MRPS27 | Q92552 | MRPS28 | P82673, Q9Y2Q9 | MRPS30 | Q9NP92 |
| MRPS31 | Q92665 | MRPS33 | Q9Y291 | MRPS34 | P82930 |
| MRPS35 | P82673, Q9Y2Q9 | MRPS5 | P82675 | MRPS6 | P82932 |
| MRPS7 | Q9Y2R9 | MRPS9 | P82933 | MTIF2 | P46199 |
| MTIF3 | Q9H2K0 | OXA1L | Q15070 | PTCD3 | Q96EY7 |

#### Interactors found in this pathway (4)

| Input | UniProt Id | Interacts with | Input | UniProt Id | Interacts with |
|---|---|---|---|---|---|
| CD47 | Q08722-3 | Q9H2K0 | FXYD3 | Q14802-3 | Q9H2K0 |
| LDLR | P01130 | Q9H2K0 | MRFAP1 | Q9Y605 | Q9H2K0 |

#### 4. Mitochondrial translation elongation (R-HSA-5389840)

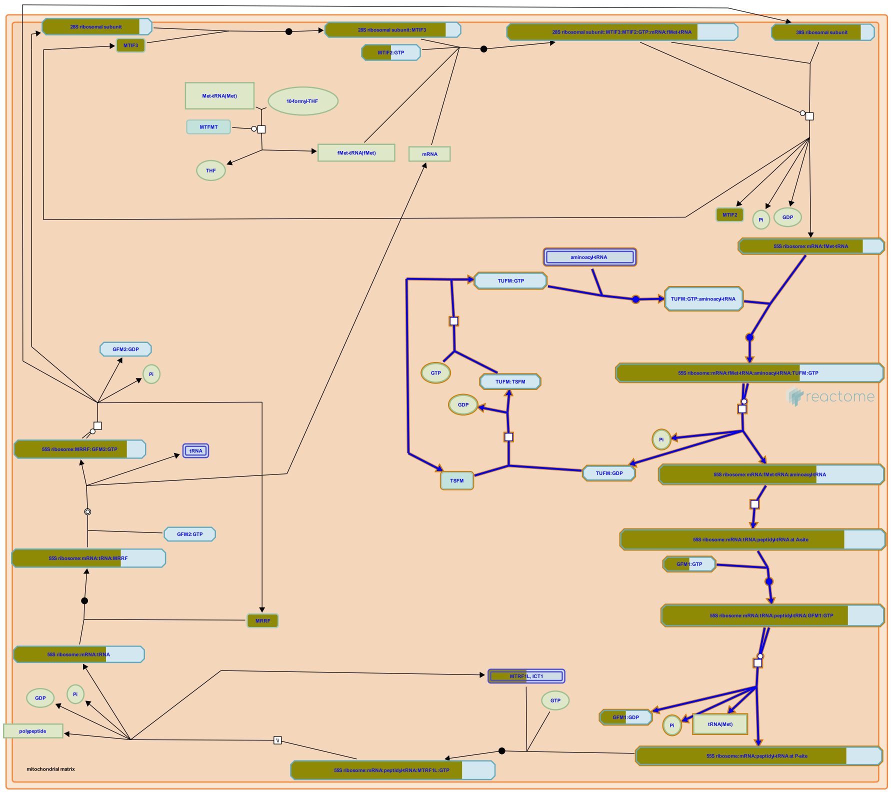


**Cellular compartments:** mitochondrial inner membrane, mitochondrial matrix.

Translation elongation proceeds by cycles of aminoacyl-tRNAs binding, peptide bond formation, and displacement of deacylated tRNAs (reviewed in Christian and Spremulli 2012). In each cycle an aminoacyl-tRNA in a complex with TUFM:GTP (EF-Tu:GTP) binds a cognate codon at the A-site of the ribosome, GTP is hydrolyzed, and TUFM:GDP dissociates. The elongating polypeptide bonded to the tRNA at the P-site is transferred to the aminoacyl group at the A-site by peptide bond formation, leaving a deacylated tRNA at the P-site and the elongating polypeptide attached to the tRNA at the A-site. GFM1:GTP (EF-Gmt:GTP) binds, GTP is hydrolyzed, GFM1:GDP dissociates, and the ribosome translocates 3 nucleotides in the 3' direction, relocating the peptidyl-tRNA to the P-site and allowing another cycle to begin. Mitochondrial ribosomes associate with the inner membrane and polypeptides are co-translationally inserted into the membrane (reviewed in Ott and Herrmann 2010, Agrawal and Sharma 2012). TUFM:GDP is regenerated to TUFM:GTP by the guanine nucleotide exchange factor TSFM (EF-Ts, EF-TsMt).

##### Edit history

| Date | Action | Author |
|---|---|---|
| 2014-04-26 | Edited | May B |
| 2014-04-26 | Authored | May B |
| 2014-04-30 | Created | May B |
| 2014-08-29 | Reviewed | Chrzanowska-Lightowlers ZM |
| 2014-09-20 | Reviewed | Spremulli LL |
| 2023-08-26 | Modified | Wright A |

#### 77 submitted entities found in this pathway, mapping to 84 Reactome entities

| Input | UniProt Id | Input | UniProt Id | Input | UniProt Id |
|---|---|---|---|---|---|
| BRIP1 | Q9P0J6 | DAP3 | P51398 | ERAL1 | O75616 |
| GADD45GIP1 | Q8TAE8 | GFM1 | Q96RP9 | ICT1 | Q14197 |
| MRPL1 | Q9BYD6 | MRPL10 | Q7Z7H8 | MRPL11 | Q9Y3B7 |
| MRPL12 | P52815 | MRPL13 | Q9BYD1 | MRPL14 | Q6P1L8 |
| MRPL15 | P49406, Q9P015 | MRPL16 | Q9NX20 | MRPL17 | Q9NRX2 |
| MRPL18 | Q9H0U6 | MRPL19 | P49406 | MRPL2 | Q5T653, Q9BZE1 |
| MRPL20 | Q9BYC9 | MRPL21 | Q7Z2W9 | MRPL22 | Q9NWU5 |
| MRPL23 | Q16540 | MRPL24 | Q96A35 | MRPL27 | Q8IXM3, Q9P0M9 |
| MRPL28 | Q13084, Q8TCC3 | MRPL3 | P09001 | MRPL32 | Q6P1L8, Q9BYC8 |
| MRPL34 | Q9BQ48 | MRPL37 | Q9BZE1 | MRPL38 | Q96DV4 |
| MRPL39 | Q9NYK5 | MRPL4 | Q9BYD3 | MRPL40 | Q9NQ50 |
| MRPL41 | Q8IXM3 | MRPL42 | Q9Y6G3 | MRPL44 | Q9H9J2 |
| MRPL45 | Q9BRJ2 | MRPL46 | Q9H2W6 | MRPL47 | Q9HD33 |
| MRPL48 | Q96GC5 | MRPL49 | Q13405 | MRPL51 | Q4U2R6 |
| MRPL52 | Q86TS9 | MRPL53 | Q96EL3 | MRPL54 | Q6P161 |
| MRPL55 | Q7Z7F7 | MRPL9 | Q9BYD2 | MRPS10 | P82664 |
| MRPS11 | P82912 | MRPS12 | O15235 | MRPS14 | O60783 |
| MRPS15 | P82914 | MRPS16 | Q9Y3D3 | MRPS17 | Q9Y2R5 |
| MRPS18A | Q9NVS2 | MRPS18B | Q9Y676 | MRPS18C | Q9Y3D5 |
| MRPS2 | Q9Y399 | MRPS21 | P82921 | MRPS22 | P82650 |
| MRPS23 | Q9Y3D9 | MRPS24 | Q96EL2 | MRPS25 | P82663 |
| MRPS26 | Q9BYN8 | MRPS27 | Q92552 | MRPS28 | P82673, Q9Y2Q9 |
| MRPS30 | Q9NP92 | MRPS31 | Q92665 | MRPS33 | Q9Y291 |
| MRPS34 | P82930 | MRPS35 | P82673, Q9Y2Q9 | MRPS5 | P82675 |
| MRPS6 | P82932 | MRPS7 | Q9Y2R9 | MRPS9 | P82933 |
| OXA1L | Q15070 | PTCD3 | Q96EY7 | | |

#### 5. Translation (R-HSA-72766)

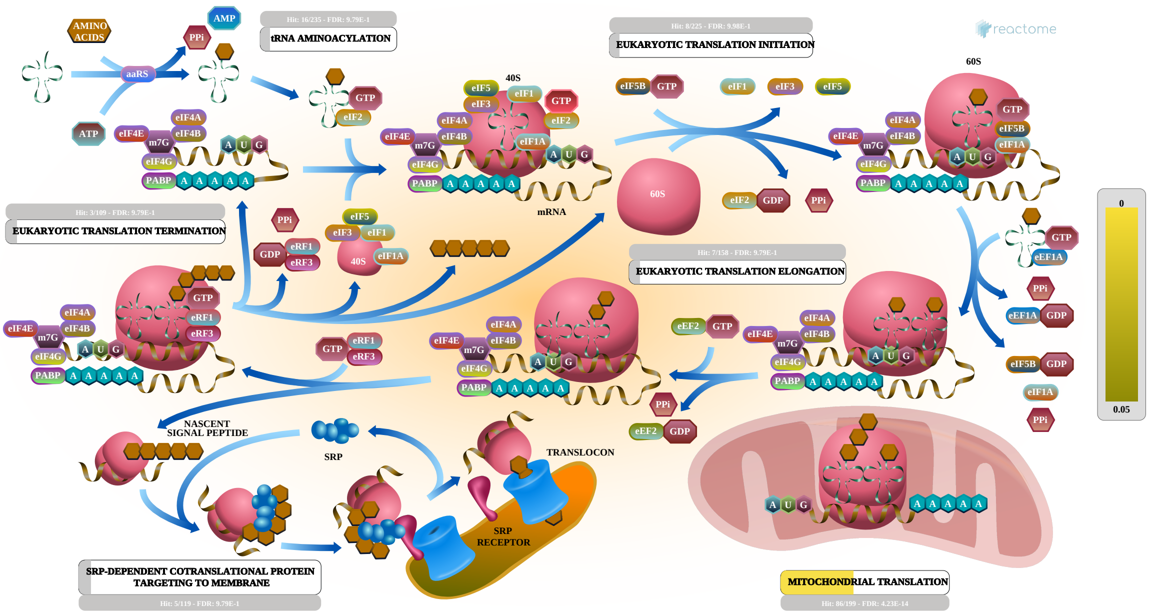


Protein synthesis is accomplished through the process of translation of an mRNA sequence into a polypeptide chain. This process can be divided into three distinct stages: initiation, elongation and termination. During the initiation phase, the two subunits of the ribosome are brought together to the translation start site on the mRNA where the polypeptide chain is to begin. Extension of the polypeptide chain occurs when a specific aminoacyl-tRNA, as determined by the template mRNA, binds an elongating ribosome. The protein chain is released from the ribosome when any one of three stop codons in the relevant reading frame on the mRNA is reached. Individual reactions at each one of these stages are catalyzed by a number of initiation, elongation and release factors, respectively.

Proteins destined for the endoplasmic reticulum (ER) contain a short sequence of hydrophobic amino acid residues (approximately 20 residues) at their N-termini. Upon protrusion of the signal sequence from the translating ribosome, the signal sequence is bound by the cytosolic signal recognition particle (SRP), translation is temporarily halted, and the SRP:nascent peptide:ribosome complex then docks with a SRP receptor complex on the ER membrane. There the nascent peptide:ribosome complex is transferred from the SRP complex to a translocon complex embedded in the ER membrane and reoriented so that the nascent polypeptide protrudes through a pore in the translocon into the ER lumen. Translation now resumes, the signal peptide is cleaved from the polypeptide by signal peptidase as the signal peptide emerges into the ER, and elongation proceeds with the growing polypeptide oriented into the ER lumen.

The 13 proteins encoded by the mitochondrial genome are translated within the mitochondrion by mitochondrial ribosomes (mitoribosomes) at the matrix face of the inner mitochondrial membrane. Mitochondrial translation reflects both the bacterial origin of the organelle and subsequent divergent evolution during symbiosis. Mitoribosomes have shorter rRNAs, mitochondria-specific proteins, and rearranged protein positions. Mitochondrial mRNAs have either no untranslated leaders or very short untranslated leaders of 1-3 nucleotides. Translation begins with N-formylmethionine, as in bacteria, and continues with cycles of aminoacyl-tRNA:TUFM:GTP binding, GTP hydrolysis and dissociation of TUFM:GDP. All 13 proteins encoded by the mitochondrial genome are hydrophobic inner membrane proteins which are inserted cotranslationally into the membrane by an interaction with OXA1L. Translation is terminated when MTRF1L:GTP recognizes a UAA or UAG codon at the A-site of the mitoribosome. The translated polypeptide is released and MRRF and GFM2:GTP act to dissociate the 55S ribosome into 28S and 39S subunits.

#### References

#### Edit history

| Date | Action | Author |
|---|---|---|
| 2004-09-08 | Revised | Kinzy TG |
| 2005-03-29 | Authored | Gebauer F, Balar B, Ulloque R, Ortiz PA, Pittman YR et al. |
| 2005-03-29 | Created | Gebauer F, Balar B, Ulloque R, Ortiz PA, Pittman YR et al. |
| 2013-11-25 | Edited | Tello-Ruiz MK, Matthews L, Gopinathrao G, Gillespie ME |
| 2023-08-24 | Reviewed | Hinnebusch AG, Sonenberg N, Hershey JW |
| 2023-08-26 | Modified | Wright A |

#### 97 submitted entities found in this pathway, mapping to 104 Reactome entities

| Input | UniProt Id | Input | UniProt Id | Input | UniProt Id |
|---|---|---|---|---|---|
| AARS2 | Q5JTZ9 | BRIP1 | Q9P0J6 | CARS | P49589 |
| DAP3 | P51398 | DARS2 | Q6PI48 | EARS2 | Q5JPH6 |
| ERAL1 | O75616 | GADD45GIP1 | Q8TAE8 | GARS | P41250 |
| GFM1 | Q96RP9 | ICT1 | Q14197 | LARS2 | Q15031 |
| MRPL1 | Q9BYD6 | MRPL10 | Q7Z7H8 | MRPL11 | Q9Y3B7 |
| MRPL12 | P52815 | MRPL13 | Q9BYD1 | MRPL14 | Q6P1L8 |
| MRPL15 | P49406, Q9P015 | MRPL16 | Q9NX20 | MRPL17 | Q9NRX2 |
| MRPL18 | Q9H0U6 | MRPL19 | P49406 | MRPL2 | Q5T653, Q9BZE1 |
| MRPL20 | Q9BYC9 | MRPL21 | Q7Z2W9 | MRPL22 | Q9NWU5 |
| MRPL23 | Q16540 | MRPL24 | Q96A35 | MRPL27 | Q8IXM3, Q9P0M9 |
| MRPL28 | Q13084, Q8TCC3 | MRPL3 | P09001 | MRPL32 | Q6P1L8, Q9BYC8 |
| MRPL34 | Q9BQ48 | MRPL37 | Q9BZE1 | MRPL38 | Q96DV4 |
| MRPL39 | Q9NYK5 | MRPL4 | Q9BYD3 | MRPL40 | Q9NQ50 |
| MRPL41 | Q8IXM3 | MRPL42 | Q9Y6G3 | MRPL44 | Q9H9J2 |
| MRPL45 | Q9BRJ2 | MRPL46 | Q9H2W6 | MRPL47 | Q9HD33 |
| MRPL48 | Q96GC5 | MRPL49 | Q13405 | MRPL51 | Q4U2R6 |
| MRPL52 | Q86TS9 | MRPL53 | Q96EL3 | MRPL54 | Q6P161 |
| MRPL55 | Q7Z7F7 | MRPL9 | Q9BYD2 | MRPS10 | P82664 |
| MRPS11 | P82912 | MRPS12 | O15235 | MRPS14 | O60783 |
| MRPS15 | P82914 | MRPS16 | Q9Y3D3 | MRPS17 | Q9Y2R5 |

| Input | UniProt Id | Input | UniProt Id | Input | UniProt Id |
|---|---|---|---|---|---|
| MRPS18A | Q9NVS2 | MRPS18B | Q9Y676 | MRPS18C | Q9Y3D5 |
| MRPS2 | Q9Y399 | MRPS21 | P82921 | MRPS22 | P82650 |
| MRPS23 | Q9Y3D9 | MRPS24 | Q96EL2 | MRPS25 | P82663 |
| MRPS26 | Q9BYN8 | MRPS27 | Q92552 | MRPS28 | P82673, Q9Y2Q9 |
| MRPS30 | Q9NP92 | MRPS31 | Q92665 | MRPS33 | Q9Y291 |
| MRPS34 | P82930 | MRPS35 | P82673, Q9Y2Q9 | MRPS5 | P82675 |
| MRPS6 | P82932 | MRPS7 | Q9Y2R9 | MRPS9 | P82933 |
| MRRF | Q96E11 | MTIF2 | P46199 | MTIF3 | Q9H2K0 |
| NARS2 | Q96I59 | OXA1L | Q15070 | PTCD3 | Q96EY7 |
| RARS2 | Q5T160 | RPL26L1 | Q9UNX3 | RPLP1 | P05386 |
| RPS27L | Q71UM5 | SPC25 | Q15005 | TARS2 | Q9BW92 |
| TRAM1 | Q15629 | WARS | P23381 | YARS | P54577 |
| YARS2 | Q9Y2Z4 | | | | |

#### Interactors found in this pathway (16)

| Input | UniProt Id | Interacts with | Input | UniProt Id | Interacts with |
|---|---|---|---|---|---|
| AGO1 | Q9UL18 | P11940 | C11orf68 | Q9H3H3 | Q15056 |
| CD47 | Q08722-3 | Q9H2K0 | DPH2 | Q9BQC3 | O95363 |
| ECH1 | Q13011 | P26641 | FXYD3 | Q14802-3 | Q9H2K0 |
| HSD17B10 | Q99714 | P13639 | KANK2 | Q63ZY3 | P06730 |
| LDLR | P01130 | Q9H2K0 | MRFAP1 | Q9Y605 | Q9H2K0 |
| PLK1 | P53350 | P23588 | SYPL1 | Q16563 | Q9BW92 |
| TGM2 | P21980-2 | Q5T160 | TPT1 | P13693 | P29692 |
| TRIM27 | P14373 | Q96E11, P47897, P06730, O95363 | UQCRC1 | P31930 | P26641 |

#### 6. The citric acid (TCA) cycle and respiratory electron transport (R-HSA-1428517)

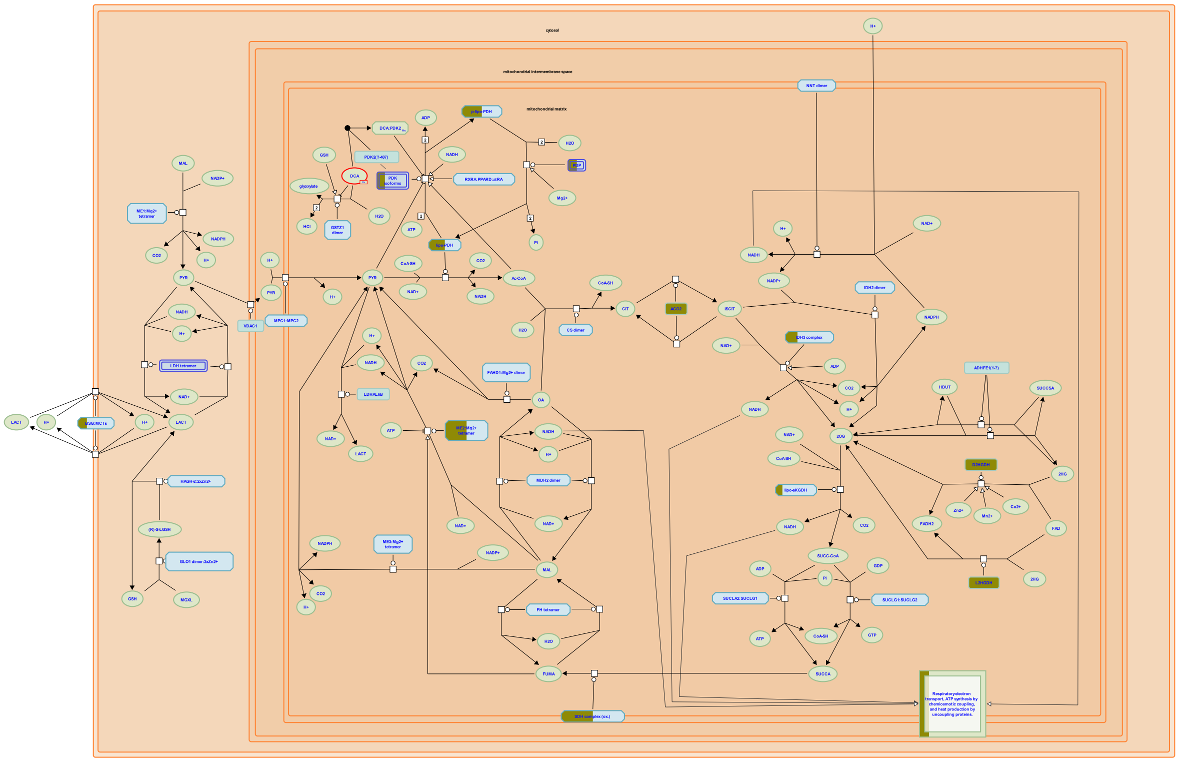

The metabolism of pyruvate provides one source of acetyl-CoA which enters the citric acid (TCA, tricarboxylic acid) cycle to generate energy and the reducing equivalent NADH. These reducing equivalents are re-oxidized back to NAD+ in the electron transport chain (ETC), coupling this process with the export of protons across the inner mitochondrial membrane. The chemiosmotic gradient created is used to drive ATP synthesis.

##### References

##### Edit history

| Date | Action | Author |
|---|---|---|
| 2003-11-03 | Authored | Birney E, Schmidt EE, D'Eustachio P |
| 2011-07-07 | Edited | Jassal B |
| 2011-07-07 | Created | Jassal B |
| 2023-08-26 | Modified | Wright A |

##### 65 submitted entities found in this pathway, mapping to 66 Reactome entities

| Input | UniProt Id | Input | UniProt Id | Input | UniProt Id |
|---|---|---|---|---|---|
| ACAD9 | Q9H845 | ACO2 | Q99798 | ATP5H | O75947 |
| ATP5J2 | P56134 | ATP5L | O75964 | COX11 | Q9Y6N1 |
| COX16 | Q9P0S2 | COX4I1 | P13073 | COX7A2L | O14548 |
| D2HGDH | Q8N465 | DLAT | P10515 | ECSIT | Q9BQ95 |
| IDH1 | P51553 | IDH3G | P51553 | L2HGDH | Q9H9P8 |
| ME2 | P23368 | MT-CO1 | P00395 | MT-CO2 | P00403 |
| MT-ND1 | P03886 | MT-ND4 | P03905 | MT-ND5 | P03915 |

| Input | UniProt Id | Input | UniProt Id | Input | UniProt Id |
|---|---|---|---|---|---|
| NDUFA10 | O95299 | NDUFA12 | Q9UI09 | NDUFA13 | Q9P0J0 |
| NDUFA2 | O43678 | NDUFA3 | O95167 | NDUFA4 | O00483 |
| NDUFA5 | Q16718 | NDUFA6 | P56556 | NDUFA9 | Q16795 |
| NDUFAB1 | O14561 | NDUFAF1 | Q9Y375 | NDUFAF2 | Q8N183 |
| NDUFAF4 | Q9P032 | NDUFB10 | O96000 | NDUFB11 | Q9NX14 |
| NDUFB4 | O95168 | NDUFB6 | O95139 | NDUFB9 | Q9Y6M9 |
| NDUFS1 | P28331 | NDUFS2 | O75306 | NDUFS3 | O75489 |
| NDUFS4 | O43181 | NDUFS5 | O43920 | NDUFS7 | O75251 |
| NDUFS8 | O00217 | NDUFV1 | P49821 | NDUFV2 | P19404 |
| OGDH | Q02218 | PDHA1 | P08559, P29803 | PDHB | P11177 |
| PDK3 | Q15120 | PDP1 | Q9P0J1 | PDPR | Q8NCN5 |
| SDHA | P31040 | SDHB | P21912 | SDHC | Q99643 |
| SLC16A1 | P53985 | SURF1 | Q15526 | TACO1 | Q9BSH4 |
| TMEM126A | Q8IUX1 | UQCRC1 | P31930 | UQCRC2 | P22695 |
| UQCRFS1 | P47985 | UQCRH | P07919 | | |

#### Interactors found in this pathway (39)

| Input | UniProt Id | Interacts with | Input | UniProt Id | Interacts with |
|---|---|---|---|---|---|
| ACAD9 | Q9H845 | O75489, Q9NPL8 | ARL6IP6 | Q8N6S5 | Q9NPL8 |
| CDK1 | P06493 | P21796 | COX4I1 | P13073 | O75489 |
| ECSIT | Q9BQ95 | O75489, Q9NPL8, P03905 | FXYD3 | Q14802-3 | Q9NPL8 |
| HSD17B10 | Q99714 | Q12931 | MT-ND1 | P03886 | O75489 |
| MT-ND4 | P03905 | O75380 | MT-ND5 | P03915 | O75489 |
| NDRG1 | Q92597 | Q9BYZ2 | NDUFA12 | Q9UI09 | O75380, O75489 |
| NDUFA13 | Q9P0J0 | Q9UI09, O75489, Q9P032, O43181 | NDUFA2 | O43678 | O75380, Q9UI09, O75489, P49821, O43920, O43181 |
| NDUFA3 | O95167 | O75489 | NDUFA4 | O00483 | Q9UI09 |
| NDUFA5 | Q16718 | O75489 | NDUFA6 | P56556 | O75380, Q9UI09, O75489, O43920 |
| NDUFA9 | Q16795 | P19404, O75380, O75489 | NDUFAF1 | Q9Y375 | O75489, Q9NPL8, O43920 |
| NDUFAF4 | Q9P032 | O75489, Q9NPL8, O43920, Q9BU61 | NDUFB10 | O96000 | O75489 |
| NDUFB11 | Q9NX14 | O75489, Q9NPL8 | NDUFB4 | O95168 | O75489 |
| NDUFB6 | O95139 | O75489 | NDUFB9 | Q9Y6M9 | O75489 |
| NDUFS1 | P28331 | O75380, O75489 | NDUFS2 | O75306 | P19404, O75380, O75489, Q9BU61 |
| NDUFS3 | O75489 | P03886, O95139, Q9UI09, P03923, Q9P032, P03915, O43920, Q9BU61, P19404, O75380, P49821, P56181, O43181 | NDUFS4 | O43181 | O75489 |
| NDUFS5 | O43920 | O75380, O75489, Q9P032 | NDUFS7 | O75251 | Q9UI09, O75489, Q9P032, Q5TEU4 |
| NDUFS8 | O00217 | O75380, O75489 | NDUFV1 | P49821 | O75489, P56181 |
| NDUFV2 | P19404 | O75380, O75489 | PDK3 | Q15120 | Q15119 |
| PDZK1 | Q5T2W1 | P19404 | | | |

| Input | ChEBI Id | Interacts with | Input | ChEBI Id | Interacts with |
|---|---|---|---|---|---|
| IDH1 | O75874 | 16810 | MMADHC | Q9H3L0, Q99LS1 | 16856 |

#### 7. Respiratory electron transport, ATP synthesis by chemiosmotic coupling, and heat production by uncoupling proteins. (R-HSA-163200)

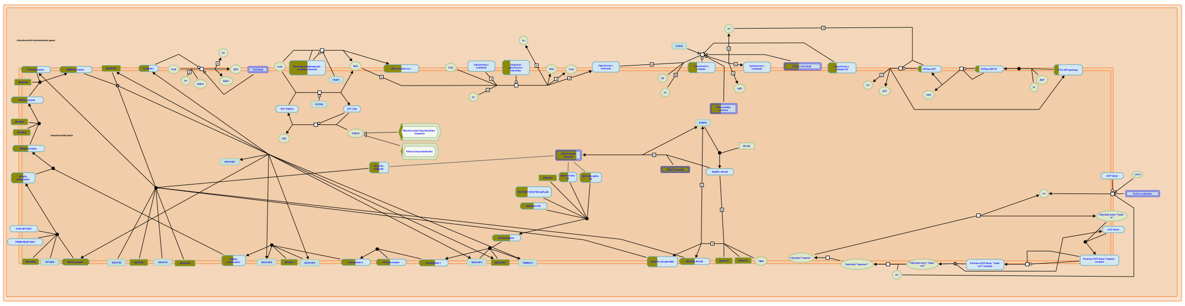

Oxidation of fatty acids and pyruvate in the mitochondrial matrix yield large amounts of NADH. The respiratory electron transport chain couples the re-oxidation of this NADH to NAD+ to the export of protons from the mitochonrial matrix, generating a chemiosmotic gradient across the inner mitochondrial membrane. This gradient is used to drive the synthesis of ATP; it can also be bypassed by uncoupling proteins to generate heat, a reaction in brown fat that may be important in regulation of body temperature in newborn children.

##### References

##### Edit history

| Date | Action | Author |
|---|---|---|
| 2005-04-21 | Authored | Jassal B |
| 2005-04-21 | Created | Jassal B |
| 2005-05-12 | Reviewed | Ferguson SJ |
| 2023-08-26 | Modified | Wright A |

##### 51 submitted entities found in this pathway, mapping to 51 Reactome entities

| Input | UniProt Id | Input | UniProt Id | Input | UniProt Id |
|---|---|---|---|---|---|
| ACAD9 | Q9H845 | ATP5H | O75947 | ATP5J2 | P56134 |
| ATP5L | O75964 | COX11 | Q9Y6N1 | COX16 | Q9P0S2 |
| COX4I1 | P13073 | COX7A2L | O14548 | ECSIT | Q9BQ95 |
| MT-CO1 | P00395 | MT-CO2 | P00403 | MT-ND1 | P03886 |
| MT-ND4 | P03905 | MT-ND5 | P03915 | NDUFA10 | O95299 |
| NDUFA12 | Q9UI09 | NDUFA13 | Q9P0J0 | NDUFA2 | O43678 |
| NDUFA3 | O95167 | NDUFA4 | O00483 | NDUFA5 | Q16718 |
| NDUFA6 | P56556 | NDUFA9 | Q16795 | NDUFAB1 | O14561 |
| NDUFAF1 | Q9Y375 | NDUFAF2 | Q8N183 | NDUFAF4 | Q9P032 |
| NDUFB10 | O96000 | NDUFB11 | Q9NX14 | NDUFB4 | O95168 |
| NDUFB6 | O95139 | NDUFB9 | Q9Y6M9 | NDUFS1 | P28331 |
| NDUFS2 | O75306 | NDUFS3 | O75489 | NDUFS4 | O43181 |
| NDUFS5 | O43920 | NDUFS7 | O75251 | NDUFS8 | O00217 |
| NDUFV1 | P49821 | NDUFV2 | P19404 | SDHA | P31040 |
| SDHB | P21912 | SDHC | Q99643 | SURF1 | Q15526 |
| TACO1 | Q9BSH4 | TMEM126A | Q8IUX1 | UQCRC1 | P31930 |
| UQCRC2 | P22695 | UQCRFS1 | P47985 | UQCRH | P07919 |

#### Interactors found in this pathway (34)

| Input | UniProt Id | Interacts with |
|---|---|---|
| ACAD9 | Q9H845 | O75489, Q9NPL8 |
| COX4I1 | P13073 | O75489 |
| FXYD3 | Q14802-3 | Q9NPL8 |
| MT-ND1 | P03886 | O75489 |
| MT-ND5 | P03915 | O75489 |
| NDUFA13 | Q9P0J0 | Q9UI09, O75489, Q9P032, O43181 |
| NDUFA3 | O95167 | O75489 |
| NDUFA5 | Q16718 | O75489 |
| NDUFA9 | Q16795 | P19404, O75380, O75489 |
| NDUFAF4 | Q9P032 | O75489, Q9NPL8, O43920, Q9BU61 |
| NDUFB11 | Q9NX14 | O75489, Q9NPL8 |
| NDUFB6 | O95139 | O75489 |
| NDUFS1 | P28331 | O75380, O75489 |
| NDUFS3 | O75489 | P03886, O95139, Q9UI09, P03923, Q9P032, P03915, O43920, Q9BU61, P19404, O75380, P49821, P56181, O43181 |
| NDUFS5 | O43920 | O75380, O75489, Q9P032 |
| NDUFS8 | O00217 | O75380, O75489 |
| NDUFV2 | P19404 | O75380, O75489 |

| Input | UniProt Id | Interacts with |
|---|---|---|
| ARL6IP6 | Q8N6S5 | Q9NPL8 |
| ECSIT | Q9BQ95 | O75489, Q9NPL8, P03905 |
| HSD17B10 | Q99714 | Q12931 |
| MT-ND4 | P03905 | O75380 |
| NDUFA12 | Q9UI09 | O75380, O75489 |
| NDUFA2 | O43678 | O75380, Q9UI09, O75489, P49821, O43920, O43181 |
| NDUFA4 | O00483 | Q9UI09 |
| NDUFA6 | P56556 | O75380, Q9UI09, O75489, O43920 |
| NDUFAF1 | Q9Y375 | O75489, Q9NPL8, O43920 |
| NDUFB10 | O96000 | O75489 |
| NDUFB4 | O95168 | O75489 |
| NDUFB9 | Q9Y6M9 | O75489 |
| NDUFS2 | O75306 | P19404, O75380, O75489, Q9BU61 |
| NDUFS4 | O43181 | O75489 |
| NDUFS7 | O75251 | Q9UI09, O75489, Q9P032, Q5TEU4 |
| NDUFV1 | P49821 | O75489, P56181 |
| PDZK1 | Q5T2W1 | P19404 |

#### 8. Respiratory electron transport (R-HSA-611105)

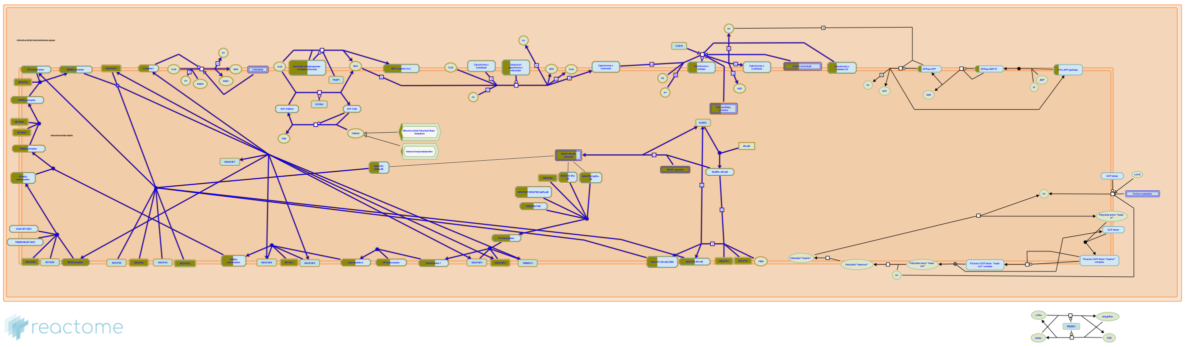

Mitochondria are often described as the "powerhouse" of a cell as it is here that energy is largely released from the oxidation of food. Reducing equivalents generated from beta-oxidation of fatty acids and from the Krebs cycle enter the electron transport chain (also called the respiratory chain). During a series of redox reactions, electrons travel down the chain releasing their energy in controlled steps. These reactions drive the active transport of protons from the mitochondrial matrix , through the inner membrane to the intermembrane space. The respiratory chain consists of five main types of carrier; flavins, iron-sulfur centres, quinones, cytochromes (heme proteins) and copper. The two main reducing equivalents entering the respiratory chain are NADH and FADH2. NADH is linked through the NADH-specific dehydrogenase whereas FADH2 is reoxidised within succinate dehydrogenase and a ubiquinone reductase of the fatty acid oxidation pathway. Oxygen is the final acceptor of electrons and with protons, is converted to form water, the end product of aerobic cellular respiration. A proton electrochemical gradient (often called protonmotive force) is established across the inner membrane, with positive charge in the intermembrane space relative to the matrix. Protons driven by the proton-motive force, can enter ATP synthase thus returning to the mitochondrial matrix. ATP synthases use this exergonic flow to form ATP in the matrix, a process called chemiosmotic coupling. A by-product of this process is heat generation.

An antiport, ATP-ADP translocase, preferentially exports ATP from the matrix thereby maintaining a high ADP:ATP ratio in the matrix. The tight coupling of electron flow to ATP synthesis means oxygen consumption is dependent on ADP availability (termed respiratory control). High ADP (low ATP) increases electron flow thereby increasing oxygen consumption and low ADP (high ATP) decreases electron flow and thereby decreases oxygen consumption. There are many inhibitors of mitochondrial ATP synthesis. Most act by either blocking the flow of electrons (eg cyanide, carbon monoxide, rotenone) or uncoupling electron flow from ATP synthesis (eg dinitrophenol). Thermogenin is a natural protein found in brown fat. Newborn babies have a large amount of brown fat and the heat generated by thermogenin is an alternative to ATP synthesis (and thus electron flow only produces heat) and allows the maintenance of body temperature in newborns.

The electron transport chain is located in the inner mitochondrial membrane and comprises some 80 proteins organized in four enzymatic complexes (I-IV). Complex V generates ATP but has no electron transfer activity. In addition to these 5 complexes, there are also two electron shuttle molecules; Coenzyme Q (also known as ubiquinone, CoQ) and Cytochrome c (Cytc). These two molecules shuttle electrons between the large complexes in the chain.

How many ATPs are generated by this process? Theoretically, for each glucose molecule, 32 ATPs can be produced. As electrons drop from NADH to oxygen in the chain, the number of protons pumped out and returning through ATP synthase can produce 2.5 ATPs per electron pair. For each pair donated by FADH2, only 1.5 ATPs can be formed. Twelve pairs of electrons are removed from each glucose molecule;

10 by NAD+ = 25 ATPs

2 by FADH2 = 3 ATPs.

Making a total of 28 ATPs. However, 2 ATPs are formed during the Krebs' cycle and 2 ATPs formed during glycolysis for each glucose molecule therefore making a total ATP yield of 32 ATPs. In reality, the energy from the respiratory chain is used for other processes (such as active transport of important ions and molecules) so under conditions of normal respiration, the actual ATP yield probably does not reach 32 ATPs.

The reducing equivalents that fuel the electron transport chain, namely NADH and FADH2, are produced by the Krebs cycle (TCA cycle) and the beta-oxidation of fatty acids. At three steps in the Krebs cycle (isocitrate conversion to oxoglutarate; oxoglutarate conversion to succinyl-CoA; Malate conversion to oxaloacetate), a pair of electrons (2e-) are removed and transferred to NAD+, forming NADH and H+. At a single step, a pair of electrons are removed from succinate, reducing FAD to FADH2. From the beta-oxidation of fatty acids, one step in the process forms NADH and H+ and another step forms FADH2.

Cytoplasmic NADH, generated from glycolysis, has to be oxidized to reform NAD+, essential for glycolysis, otherwise glycolysis would cease to function. There is no carrier that transports NADH directly into the mitochondrial matrix and the inner mitochondrial membrane is impermeable to NADH so the cell uses two shuttle systems to move reducing equivalents into the mitochondrion and regenerate cytosolic NAD+.

The first is the glycerol phosphate shuttle, which uses electrons from cytosolic NADH to produce FADH2 within the inner membrane. These electrons then flow to Coenzyme Q. Complex I is bypassed so only 1.5 ATPs can be formed per NADH via this route. The overall balanced equation, summing all the reactions in this system, is

NADH (cytosol) + H+ (cytosol) + NAD+ (mito.) = NAD+ (cytosol) + NADH (mito.) + H+ (mito.)

The malate-aspartate shuttle uses the oxidation of malate to generate NADH in the mitochondrial matrix. This NADH can then be fed directly to complex I and thus can form 3 ATPs via the respiratory chain. The overall balanced equation is

NADH (cytosol) + H+ (cytosol) + FAD (inner memb.) = NAD+ (cytosol) + FADH2 (inner memb.)

Both of these shuttle systems regenerate cytosolic NAD+.

The entry point for NADH is complex I (NADH dehydrogenase) and the entry point for FADH2 is Coenzyme Q. The input of electrons from fatty acid oxidation via ubiquinone is complicated and not shown in the diagram.

#### Edit history

| Date | Action | Author |
|---|---|---|
| 2005-04-21 | Authored | Jassal B |
| 2005-05-12 | Reviewed | Ferguson SJ |
| 2010-04-19 | Created | D'Eustachio P |
| 2015-04-27 | Revised | Jassal B |
| 2023-08-26 | Modified | Wright A |

#### 48 submitted entities found in this pathway, mapping to 48 Reactome entities

| Input | UniProt Id | Input | UniProt Id | Input | UniProt Id |
|---|---|---|---|---|---|
| ACAD9 | Q9H845 | COX11 | Q9Y6N1 | COX16 | Q9P0S2 |
| COX4I1 | P13073 | COX7A2L | O14548 | ECSIT | Q9BQ95 |
| MT-CO1 | P00395 | MT-CO2 | P00403 | MT-ND1 | P03886 |
| MT-ND4 | P03905 | MT-ND5 | P03915 | NDUFA10 | O95299 |
| NDUFA12 | Q9UI09 | NDUFA13 | Q9P0J0 | NDUFA2 | O43678 |
| NDUFA3 | O95167 | NDUFA4 | O00483 | NDUFA5 | Q16718 |
| NDUFA6 | P56556 | NDUFA9 | Q16795 | NDUFAB1 | O14561 |
| NDUFAF1 | Q9Y375 | NDUFAF2 | Q8N183 | NDUFAF4 | Q9P032 |
| NDUFB10 | O96000 | NDUFB11 | Q9NX14 | NDUFB4 | O95168 |
| NDUFB6 | O95139 | NDUFB9 | Q9Y6M9 | NDUFS1 | P28331 |
| NDUFS2 | O75306 | NDUFS3 | O75489 | NDUFS4 | O43181 |
| NDUFS5 | O43920 | NDUFS7 | O75251 | NDUFS8 | O00217 |
| NDUFV1 | P49821 | NDUFV2 | P19404 | SDHA | P31040 |
| SDHB | P21912 | SDHC | Q99643 | SURF1 | Q15526 |
| TACO1 | Q9BSH4 | TMEM126A | Q8IUX1 | UQCRC1 | P31930 |
| UQCRC2 | P22695 | UQCRFS1 | P47985 | UQCRH | P07919 |

#### Interactors found in this pathway (34)

| Input | UniProt Id | Interacts with | Input | UniProt Id | Interacts with |
|---|---|---|---|---|---|
| ACAD9 | Q9H845 | O75489, Q9NPL8 | ARL6IP6 | Q8N6S5 | Q9NPL8 |
| COX4I1 | P13073 | O75489 | ECSIT | Q9BQ95 | O75489, Q9NPL8, P03905 |
| FXYD3 | Q14802-3 | Q9NPL8 | HSD17B10 | Q99714 | Q12931 |
| MT-ND1 | P03886 | O75489 | MT-ND4 | P03905 | O75380 |
| MT-ND5 | P03915 | O75489 | NDUFA12 | Q9UI09 | O75380, O75489 |
| NDUFA13 | Q9P0J0 | Q9UI09, O75489, Q9P032, O43181 | NDUFA2 | O43678 | O75380, Q9UI09, O75489, P49821, O43920, O43181 |
| NDUFA3 | O95167 | O75489 | NDUFA4 | O00483 | Q9UI09 |

| Input | UniProt Id | Interacts with | Input | UniProt Id | Interacts with |
|---|---|---|---|---|---|
| NDUFA5 | Q16718 | O75489 | NDUFA6 | P56556 | O75380, Q9UI09, O75489, O43920 |
| NDUFA9 | Q16795 | P19404, O75380, O75489 | NDUFAF1 | Q9Y375 | O75489, Q9NPL8, O43920 |
| NDUFAF4 | Q9P032 | O75489, Q9NPL8, O43920, Q9BU61 | NDUFB10 | O96000 | O75489 |
| NDUFB11 | Q9NX14 | O75489, Q9NPL8 | NDUFB4 | O95168 | O75489 |
| NDUFB6 | O95139 | O75489 | NDUFB9 | Q9Y6M9 | O75489 |
| NDUFS1 | P28331 | O75380, O75489 | NDUFS2 | O75306 | P19404, O75380, O75489, Q9BU61 |
| NDUFS3 | O75489 | P03886, O95139, Q9UI09, P03923, Q9P032, P03915, O43920, Q9BU61, P19404, O75380, P49821, P56181, O43181 | NDUFS4 | O43181 | O75489 |
| NDUFS5 | O43920 | O75380, O75489, Q9P032 | NDUFS7 | O75251 | Q9UI09, O75489, Q9P032, Q5TEU4 |
| NDUFS8 | O00217 | O75380, O75489 | NDUFV1 | P49821 | O75489, P56181 |
| NDUFV2 | P19404 | O75380, O75489 | PDZK1 | Q5T2W1 | P19404 |

#### 9. Polymerase switching (R-HSA-69091)

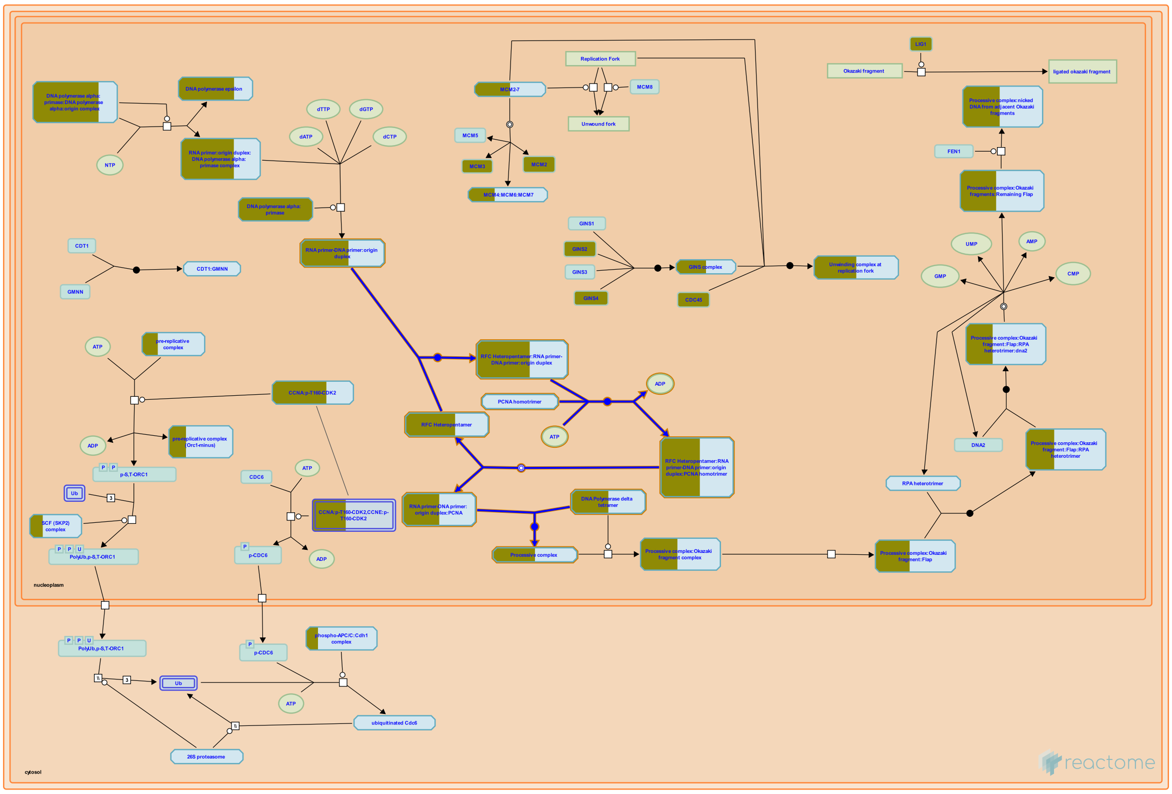

**Cellular compartments:** nucleoplasm.

After the primers are synthesized, Replication Factor C binds to the 3'-end of the initiator DNA to trigger polymerase switching. The non-processive nature of pol alpha catalytic activity and the tight binding of Replication Factor C to the primer-template junction presumably lead to the turnover of the pol alpha:primase complex. After the Pol alpha-primase primase complex is displaced from the primer, the proliferating cell nuclear antigen (PCNA) binds to form a "sliding clamp" structure. Replication Factor C then dissociates, and DNA polymerase delta binds and catalyzes the processive synthesis of DNA.


##### Edit history

| Date | Action | Author |
| --- | --- | --- |
| 2023-08-26 | Modified | Wright A |

#### 8 submitted entities found in this pathway, mapping to 9 Reactome entities

| Input | UniProt Id | Input | UniProt Id | Input | UniProt Id |
|---|---|---|---|---|---|
| POLA1 | P09884 | POLA2 | Q14181 | POLD1 | P28340 |
| POLD2 | P49005 | PRIM1 | P49642 | PRIM2 | P49643 |
| RFC1 | P35251 | RFC5 | P40937, P40938 | | |

#### 10. Leading Strand Synthesis (R-HSA-69109)

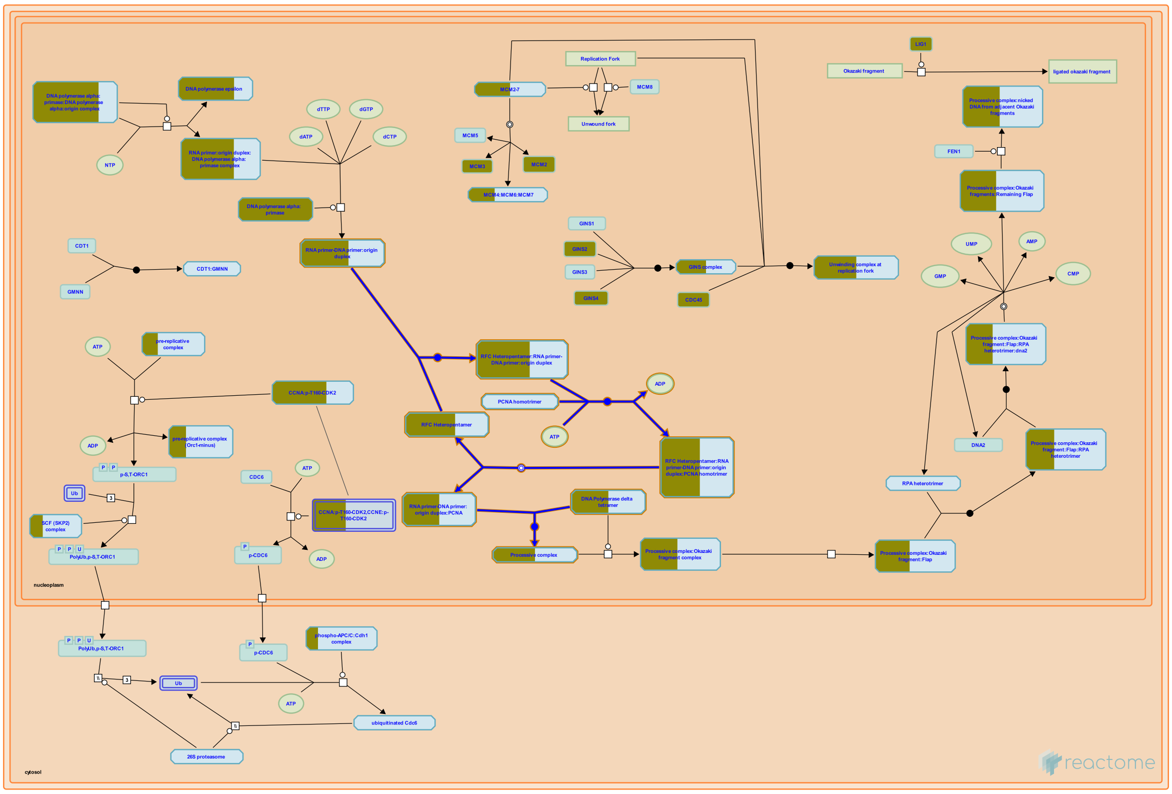

**Cellular compartments:** nucleoplasm.

The processive complex is responsible for synthesizing at least 5-10 kb of DNA in a continuous manner during leading strand synthesis. The incorporation of nucleotides by pol delta is quite accurate. However, incorporation of an incorrect nucleotide does occur occasionally. Misincorporated nucleotides are removed by the 3' to 5' exonucleolytic proofreading capability of pol delta.


##### Edit history

| Date | Action | Author |
|---|---|---|
| 2023-08-26 | Modified | Wright A |

##### 8 submitted entities found in this pathway, mapping to 9 Reactome entities

| Input | UniProt Id | Input | UniProt Id | Input | UniProt Id |
|---|---|---|---|---|---|
| POLA1 | P09884 | POLA2 | Q14181 | POLD1 | P28340 |
| POLD2 | P49005 | PRIM1 | P49642 | PRIM2 | P49643 |

| Input | UniProt Id | Input | UniProt Id | Input | UniProt Id |
|---|---|---|---|---|---|
| RFC1 | P35251 | RFC5 | P40937, P40938 | | |

#### 11. DNA strand elongation (R-HSA-69190)

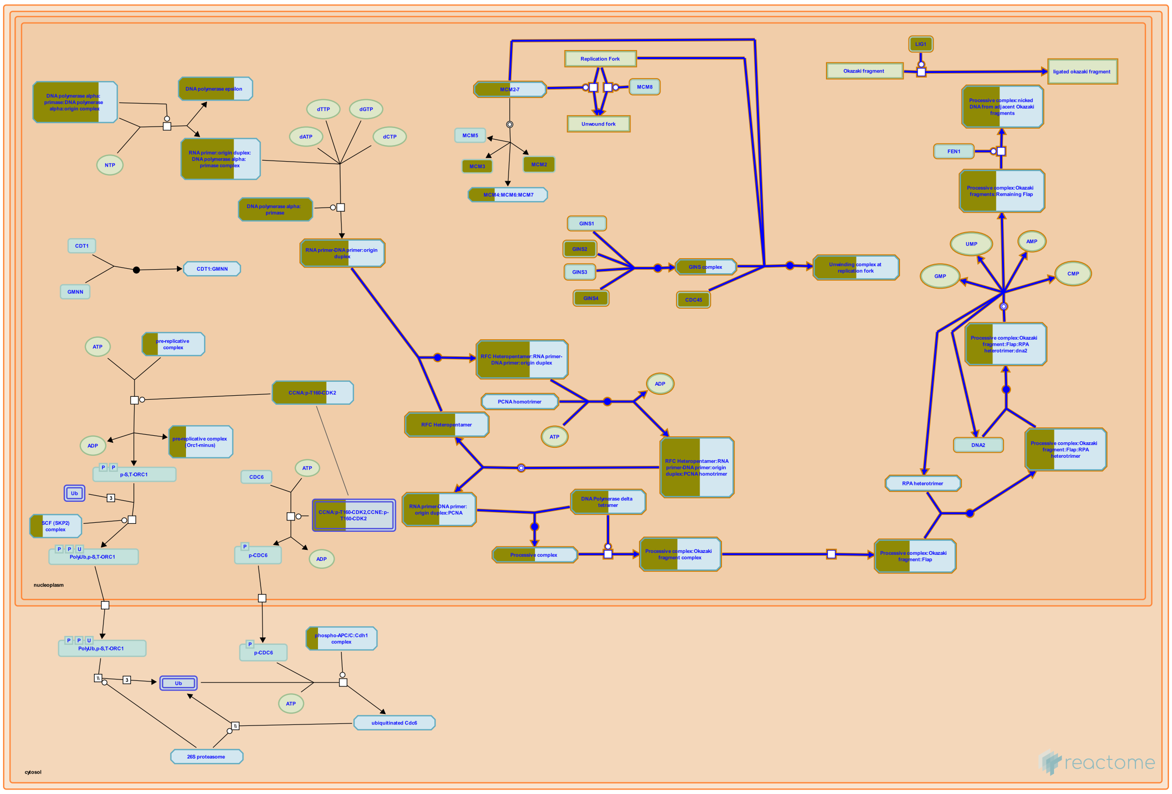

**Cellular compartments:** nucleoplasm.

Accurate and efficient genome duplication requires coordinated processes to replicate two template strands at eucaryotic replication forks. Knowledge of the fundamental reactions involved in replication fork progression is derived largely from biochemical studies of the replication of simian virus and from yeast genetic studies. Since duplex DNA forms an anti-parallel structure, and DNA polymerases are unidirectional, one of the new strands is synthesized continuously in the direction of fork movement. This strand is designated as the leading strand. The other strand grows in the direction away from fork movement, and is called the lagging strand. Several specific interactions among the various proteins involved in DNA replication underlie the mechanism of DNA synthesis, on both the leading and lagging strands, at a DNA replication fork. These interactions allow the replication enzymes to cooperate in the replication process (Hurwitz et al 1990; Brush et al 1996; Ayyagari et al 1995; Budd & Campbell 1997; Bambara et al 1997).

#### Edit history

| Date | Action | Author |
|---|---|---|
| 2003-06-05 | Authored | Tom S, Bambara RA |
| 2003-06-05 | Created | Tom S, Bambara RA |
| 2023-08-26 | Modified | Wright A |

#### 14 submitted entities found in this pathway, mapping to 16 Reactome entities

| Input | UniProt Id | Input | UniProt Id | Input | UniProt Id |
|---|---|---|---|---|---|
| CDC45 | O75419 | GINS2 | Q9Y248 | GINS4 | Q9BRT9 |
| LIG1 | P18858 | MCM2 | P33993, P49736 | MCM3 | P25205 |
| POLA1 | P09884 | POLA2 | Q14181 | POLD1 | P28340 |
| POLD2 | P49005 | PRIM1 | P49642 | PRIM2 | P49643 |
| RFC1 | P35251 | RFC5 | P40937, P40938 | | |

#### Interactors found in this pathway (7)

| Input | UniProt Id | Interacts with | Input | UniProt Id | Interacts with |
|---|---|---|---|---|---|
| CDC45 | O75419 | Q14691 | CEP55 | Q53EZ4 | O75419 |
| GINS2 | Q9Y248 | Q9BRT9, Q14691 | GINS4 | Q9BRT9 | Q9Y248, Q14691 |
| MCM2 | P49736 | O75419 | RAB4B | P61018 | O75419 |
| TOPBP1 | Q92547 | O75419 | | | |

#### 12. Lagging Strand Synthesis (R-HSA-69186)

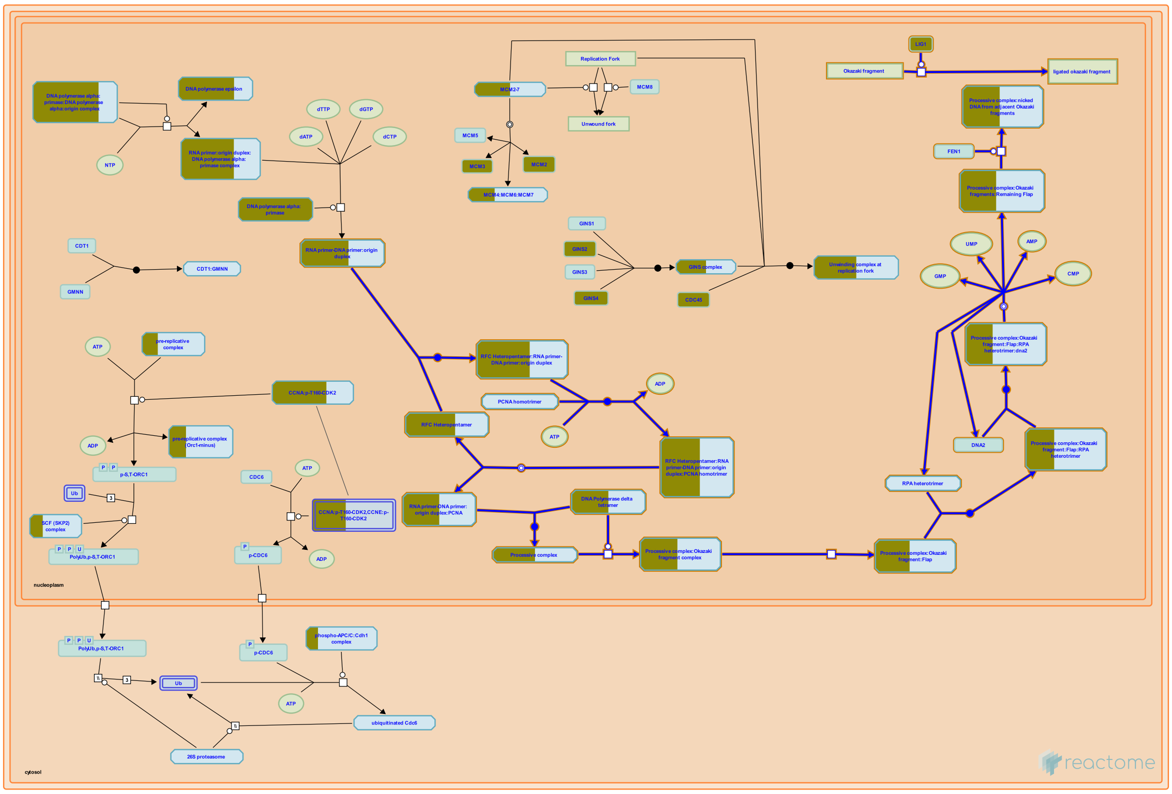

**Cellular compartments:** nucleoplasm.

Due to the antiparallel nature of DNA, DNA polymerization is unidirectional, and one strand is synthesized discontinuously. This strand is called the lagging strand. Although the polymerase switching on the lagging strand is very similar to that on the leading strand, the processive synthesis on the two strands proceeds quite differently. Short DNA fragments, about 100 bases long, called Okazaki fragments are synthesized on the RNA-DNA primers first. Strand-displacement synthesis occurs, whereby the primer-containing 5'-terminus of the adjacent Okazaki fragment is folded into a single-stranded flap structure. This flap structure is removed by endonucleases, and the adjacent Okazaki fragments are joined by DNA ligase.


##### Edit history

| Date | Action | Author |
|---|---|---|
| 2023-08-26 | Modified | Wright A |

##### 9 submitted entities found in this pathway, mapping to 10 Reactome entities

| Input | UniProt Id | Input | UniProt Id | Input | UniProt Id |
|---|---|---|---|---|---|
| LIG1 | P18858 | POLA1 | P09884 | POLA2 | Q14181 |
| POLD1 | P28340 | POLD2 | P49005 | PRIM1 | P49642 |
| PRIM2 | P49643 | RFC1 | P35251 | RFC5 | P40937, P40938 |

#### 13. DNA replication initiation (R-HSA-68952)

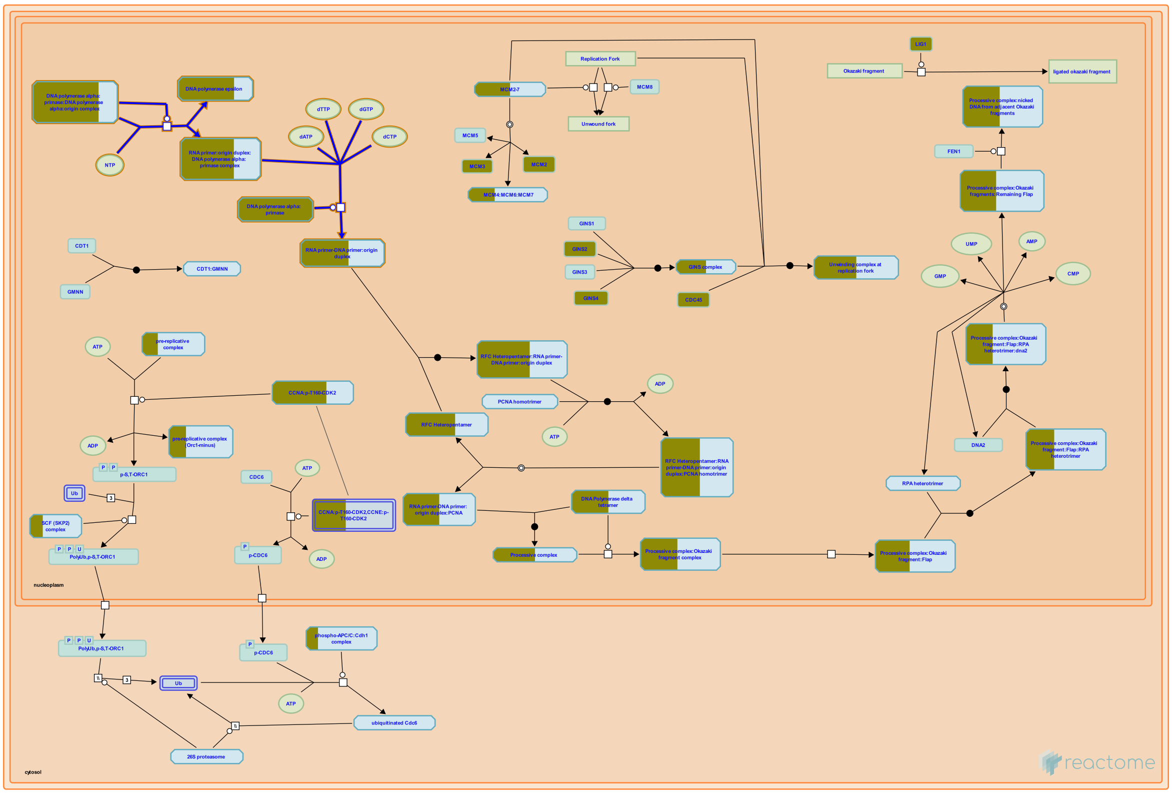


**Cellular compartments:** nucleoplasm.

DNA polymerases are not capable of de novo DNA synthesis and require synthesis of a primer, usually by a DNA-dependent RNA polymerase (primase) to begin DNA synthesis. In eukaryotic cells, the primer is synthesized by DNA polymerase alpha:primase. First, the DNA primase portion of this complex synthesizes approximately 6-10 nucleotides of RNA primer and then the DNA polymerase portion synthesizes an additional 20 nucleotides of DNA (Frick & Richardson 2002; Wang et al 1984).


##### Edit history

| Date | Action | Author |
|---|---|---|
| 2003-06-05 | Created | Davey MJ, O'Donnell M |
| 2023-08-26 | Modified | Wright A |

##### 7 submitted entities found in this pathway, mapping to 7 Reactome entities

| Input | UniProt Id | Input | UniProt Id | Input | UniProt Id |
|---|---|---|---|---|---|
| POLA1 | P09884 | POLA2 | Q14181 | POLE | Q07864 |
| POLE3 | Q9NRF9 | POLE4 | Q9NR33 | PRIM1 | P49642 |
| PRIM2 | P49643 | | | | |

#### 14. Complex I biogenesis (R-HSA-6799198)

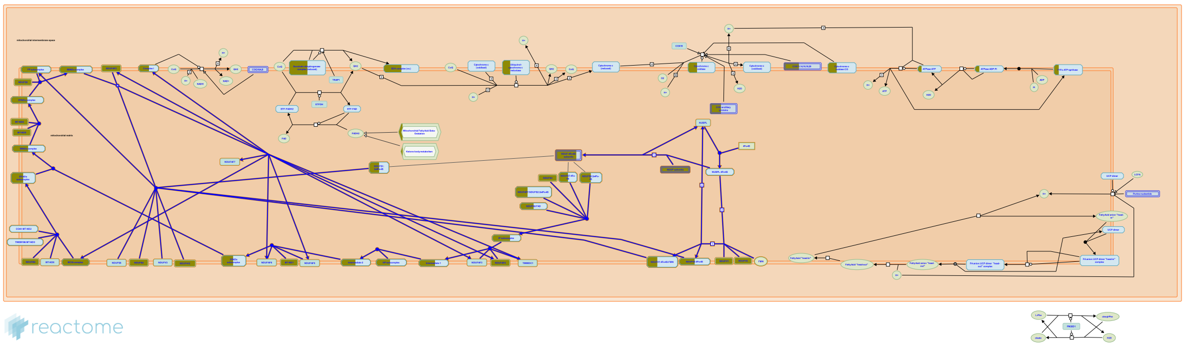

Complex I (NADH:ubiquinone oxidoreductase or NADH dehydrogenase) utilises NADH formed from glycolysis and the TCA cycle to pump protons out of the mitochondrial matrix. It is the largest enzyme complex in the electron transport chain, containing 45 subunits. Seven subunits (ND1-6, ND4L) are encoded by mitochondrial DNA, the remainder encoded in the nucleus. The enzyme has a FMN prosthetic group and 8 Iron-Sulfur (Fe-S) clusters. The subunits are assembled together in a coordinated manner via preassembled subcomplexes to form the mature holoenzyme. The so-called "assembly factor" proteins, acting intrinsically or transiently, are required for constructing complex I although their exact roles in the biogenesis are not fully understood (Fernandez-Vizarra et al. 2009, Mckenzie & Ryan 2010, Mimaki et al. 2012, Andrews et al. 2013).


##### Edit history

| Date | Action | Author |
|---|---|---|
| 2015-09-22 | Edited | Jassal B |
| 2015-09-22 | Authored | Jassal B |
| 2015-09-22 | Created | Jassal B |
| 2015-10-01 | Reviewed | Meldal BH |
| 2023-08-26 | Modified | Wright A |

##### 32 submitted entities found in this pathway, mapping to 32 Reactome entities

| Input | UniProt Id | Input | UniProt Id | Input | UniProt Id |
|---|---|---|---|---|---|
| ACAD9 | Q9H845 | ECSIT | Q9BQ95 | MT-ND1 | P03886 |
| MT-ND4 | P03905 | MT-ND5 | P03915 | NDUFA10 | O95299 |

| Input | UniProt Id | Input | UniProt Id | Input | UniProt Id |
|---|---|---|---|---|---|
| NDUFA12 | Q9UI09 | NDUFA13 | Q9P0J0 | NDUFA2 | O43678 |
| NDUFA3 | O95167 | NDUFA5 | Q16718 | NDUFA6 | P56556 |
| NDUFA9 | Q16795 | NDUFAB1 | O14561 | NDUFAF1 | Q9Y375 |
| NDUFAF2 | Q8N183 | NDUFAF4 | Q9P032 | NDUFB10 | O96000 |
| NDUFB11 | Q9NX14 | NDUFB4 | O95168 | NDUFB6 | O95139 |
| NDUFB9 | Q9Y6M9 | NDUFS1 | P28331 | NDUFS2 | O75306 |
| NDUFS3 | O75489 | NDUFS4 | O43181 | NDUFS5 | O43920 |
| NDUFS7 | O75251 | NDUFS8 | O00217 | NDUFV1 | P49821 |
| NDUFV2 | P19404 | TMEM126A | Q8IUX1 | | |

#### Interactors found in this pathway (33)

| Input | UniProt Id | Interacts with | Input | UniProt Id | Interacts with |
|---|---|---|---|---|---|
| ACAD9 | Q9H845 | O75489, Q9NPL8 | ARL6IP6 | Q8N6S5 | Q9NPL8 |
| COX4I1 | P13073 | O75489 | ECSIT | Q9BQ95 | O75489, Q9NPL8, P03905 |
| FXYD3 | Q14802-3 | Q9NPL8 | MT-ND1 | P03886 | O75489 |
| MT-ND4 | P03905 | O75380 | MT-ND5 | P03915 | O75489 |
| NDUFA12 | Q9UI09 | O75380, O75489 | NDUFA13 | Q9P0J0 | Q9UI09, O75489, Q9P032, O43181 |
| NDUFA2 | O43678 | O75380, Q9UI09, O75489, P49821, O43920, O43181 | NDUFA3 | O95167 | O75489 |
| NDUFA4 | O00483 | Q9UI09 | NDUFA5 | Q16718 | O75489 |
| NDUFA6 | P56556 | O75380, Q9UI09, O75489, O43920 | NDUFA9 | Q16795 | P19404, O75380, O75489 |
| NDUFAF1 | Q9Y375 | O75489, Q9NPL8, O43920 | NDUFAF4 | Q9P032 | O75489, Q9NPL8, O43920, Q9BU61 |
| NDUFB10 | O96000 | O75489 | NDUFB11 | Q9NX14 | O75489, Q9NPL8 |
| NDUFB4 | O95168 | O75489 | NDUFB6 | O95139 | O75489 |
| NDUFB9 | Q9Y6M9 | O75489 | NDUFS1 | P28331 | O75380, O75489 |
| NDUFS2 | O75306 | P19404, O75380, O75489, Q9BU61 | NDUFS3 | O75489 | P03886, O95139, Q9UI09, P03923, Q9P032, P03915, O43920, Q9BU61, P19404, O75380, P49821, P56181, O43181 |
| NDUFS4 | O43181 | O75489 | NDUFS5 | O43920 | O75380, O75489, Q9P032 |
| NDUFS7 | O75251 | Q9UI09, O75489, Q9P032, Q5TEU4 | NDUFS8 | O00217 | O75380, O75489 |
| NDUFV1 | P49821 | O75489, P56181 | NDUFV2 | P19404 | O75380, O75489 |
| PDZK1 | Q5T2W1 | P19404 | | | |

#### 15. Gap-filling DNA repair synthesis and ligation in GG-NER (R-HSA-5696397)

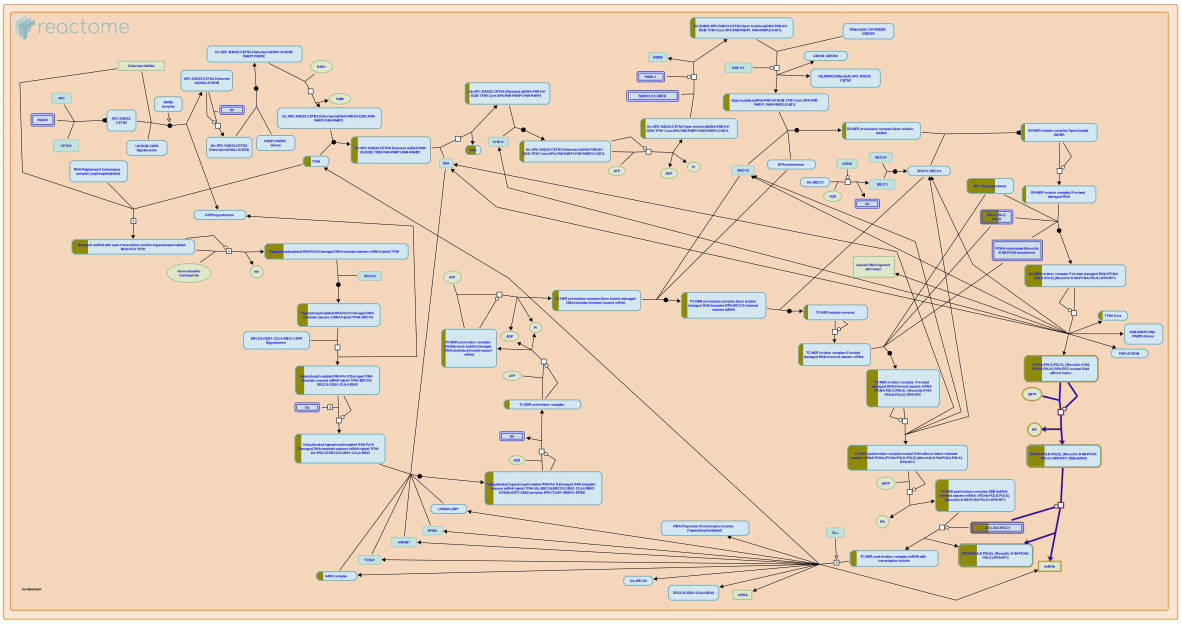

**Cellular compartments:** nucleoplasm.

Global genome nucleotide excision repair (GG-NER) is completed by DNA repair synthesis that fills the single stranded gap created after dual incision of the damaged DNA strand and excision of the ~27-30 bases long oligonucleotide that contains the lesion. DNA synthesis is performed by DNA polymerases epsilon or delta, or the Y family DNA polymerase kappa (POLK), which are loaded to the repair site after 5' incision (Staresincic et al. 2009, Ogi et al. 2010). DNA ligases LIG1 or LIG3 (as part of the LIG3:XRCC1 complex) ligate the newly synthesized stretch of oligonucleotides to the incised DNA strand (Moser et al. 2007).


##### Edit history

| Date | Action | Author |
|---|---|---|
| 2004-01-29 | Authored | Hoeijmakers JH |
| 2004-02-02 | Authored | Gopinathrao G |
| 2015-05-28 | Created | Orlic-Milacic M |
| 2015-06-16 | Revised | Orlic-Milacic M |
| 2015-06-16 | Edited | Orlic-Milacic M |

| Date | Action | Author |
|---|---|---|
| 2015-06-16 | Authored | Orlic-Milacic M |
| 2015-08-03 | Reviewed | Fousteri M |
| 2023-08-31 | Modified | Wright A |

#### 8 submitted entities found in this pathway, mapping to 9 Reactome entities

| Input | UniProt Id | Input | UniProt Id | Input | UniProt Id |
|---|---|---|---|---|---|
| LIG1 | P18858 | POLD1 | P28340 | POLD2 | P49005 |
| POLE | Q07864 | POLE3 | Q9NRF9 | POLE4 | Q9NR33 |
| RFC1 | P35251 | RFC5 | P40937, P40938 | | |

#### 16. Synthesis of DNA (R-HSA-69239)

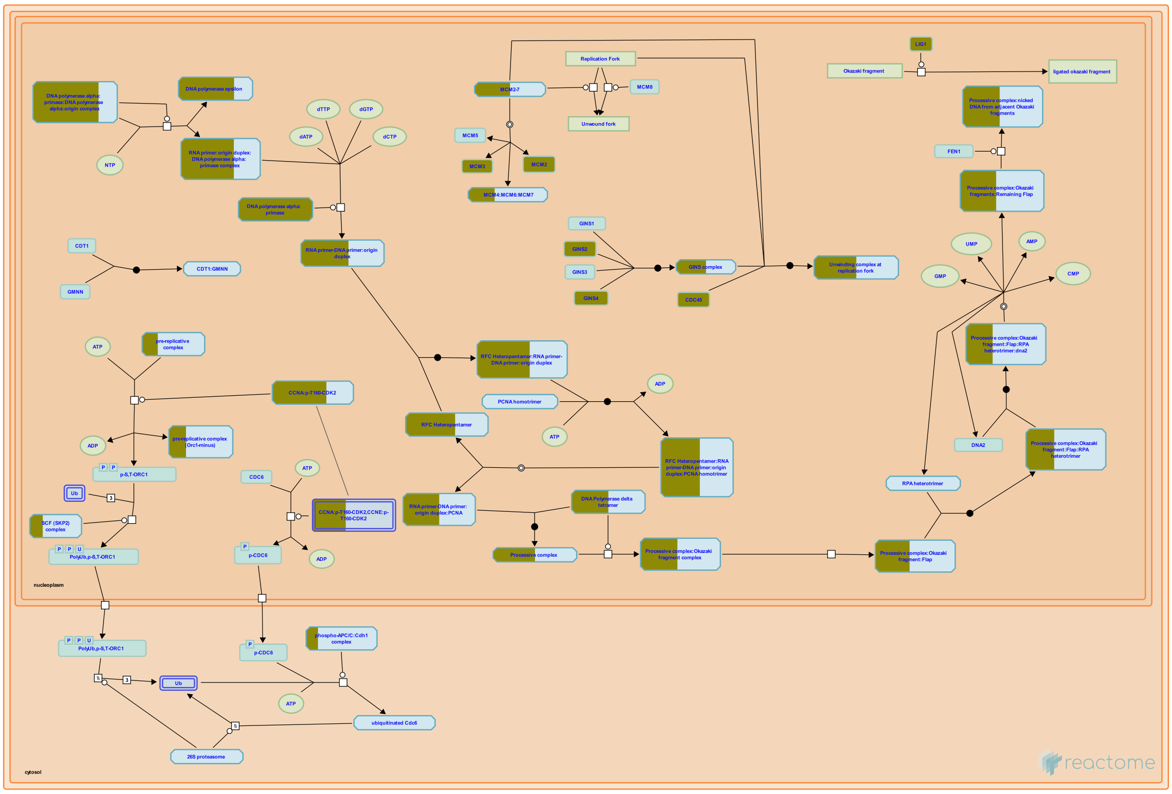

**Cellular compartments:** nucleoplasm, cytosol.

The actual synthesis of DNA occurs in the S phase of the cell cycle. This includes the initiation of DNA replication, when the first nucleotide of the new strand is laid down during the synthesis of the primer. The DNA replication preinitiation events begin in late M or early G1 phase.

##### References

##### Edit history

| Date | Action | Author |
| --- | --- | --- |
| 2023-08-26 | Modified | Wright A |

##### 24 submitted entities found in this pathway, mapping to 26 Reactome entities

| Input | UniProt Id | Input | UniProt Id | Input | UniProt Id |
| --- | --- | --- | --- | --- | --- |
| CCNA2 | P20248 | CDC23 | Q9UJX2 | CDC45 | O75419 |
| CDK2 | P24941 | GINS2 | Q9Y248 | GINS4 | Q9BRT9 |
| LIG1 | P18858 | MCM2 | P33993, P49736 | MCM3 | P25205 |
| ORC3 | Q9UBD5 | POLA1 | P09884 | POLA2 | Q14181 |
| POLD1 | P28340 | POLD2 | P49005 | POLE | Q07864 |
| POLE3 | Q9NRF9 | POLE4 | Q9NR33 | PRIM1 | P49642 |
| PRIM2 | P49643 | RFC1 | P35251 | RFC5 | P40937, P40938 |
| SKP1 | P63208 | UBE2C | O00762 | UBE2S | Q16763 |

##### Interactors found in this pathway (17)

| Input | UniProt Id | Interacts with |
|---|---|---|
| ASF1B | Q9NVP2 | P49736 |
| CDC45 | O75419 | Q14691, P49736 |
| CDK2 | P24941 | O75496, Q9H211 |
| GINS2 | Q9Y248 | Q9BRT9, Q14691 |
| KANK2 | Q63ZY3 | O75496 |
| MCM3 | P25205 | P33992, P49736 |
| PLK1 | P53350 | Q99741, P49736 |
| SAPCD2 | Q86UD0 | O75496 |
| TOPBP1 | Q92547 | O75419 |

| Input | UniProt Id | Interacts with |
|---|---|---|
| CCNA2 | P20248 | Q13415, O75496, Q9H211 |
| CDK1 | P06493 | Q99741, Q9H211 |
| CEP55 | Q53EZ4 | O75419 |
| GINS4 | Q9BRT9 | Q9Y248, Q14691 |
| MCM2 | P33993, P49736 | O75419, P33992, P25205, P49736 |
| ORC3 | Q9UBD5 | Q13415 |
| RAB4B | P61018 | O75419 |
| TFDP1 | Q14186 | O75496 |

#### 17. Gap-filling DNA repair synthesis and ligation in TC-NER (R-HSA-6782210)

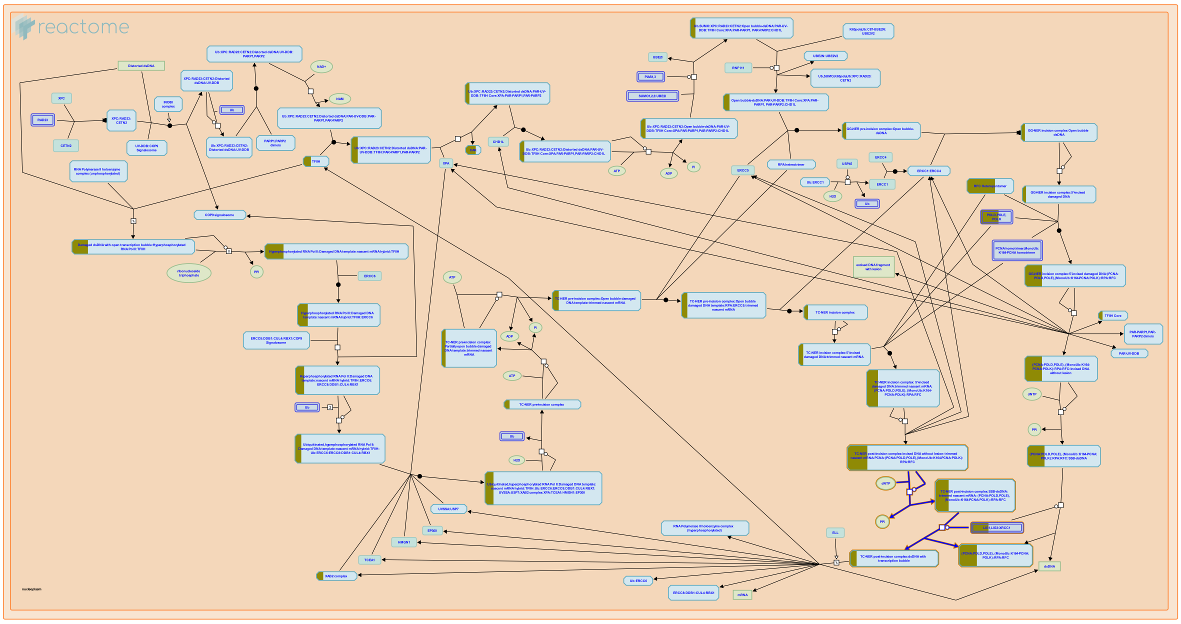

**Cellular compartments:** nucleoplasm.

In transcription-coupled nucleotide excision repair (TC-NER), similar to global genome nucleotide excision repair (GG-NER), DNA polymerases delta or epsilon, or the Y family DNA polymerase kappa, fill in the single stranded gap that remains after dual incision. DNA ligases LIG1 or LIG3, the latter in complex with XRCC1, subsequently seal the single stranded nick by ligating the 3' end of the newly synthesized patch with the 5' end of incised DNA (Moser et al. 2007, Staresincic et al. 2009, Ogi et al. 2010).


##### Edit history

| Date | Action | Author |
|---|---|---|
| 2004-01-29 | Authored | Hoeijmakers JH |
| 2015-06-05 | Created | Orlic-Milacic M |
| 2015-06-16 | Revised | Orlic-Milacic M |
| 2015-06-16 | Edited | Orlic-Milacic M |
| 2015-06-16 | Authored | Orlic-Milacic M |
| 2015-08-03 | Reviewed | Fousteri M |

| Date | Action | Author |
| --- | --- | --- |
| 2023-08-31 | Modified | Wright A |

#### 12 submitted entities found in this pathway, mapping to 13 Reactome entities

| Input | UniProt Id | Input | UniProt Id | Input | UniProt Id |
| --- | --- | --- | --- | --- | --- |
| AQR | O60306 | CCNH | P51946 | CDK7 | P50613 |
| GTF2H3 | Q13889 | LIG1 | P18858 | POLD1 | P28340 |
| POLD2 | P49005 | POLE | Q07864 | POLE3 | Q9NRF9 |
| POLE4 | Q9NR33 | RFC1 | P35251 | RFC5 | P40937, P40938 |

#### 18. Activation of the pre-replicative complex (R-HSA-68962)

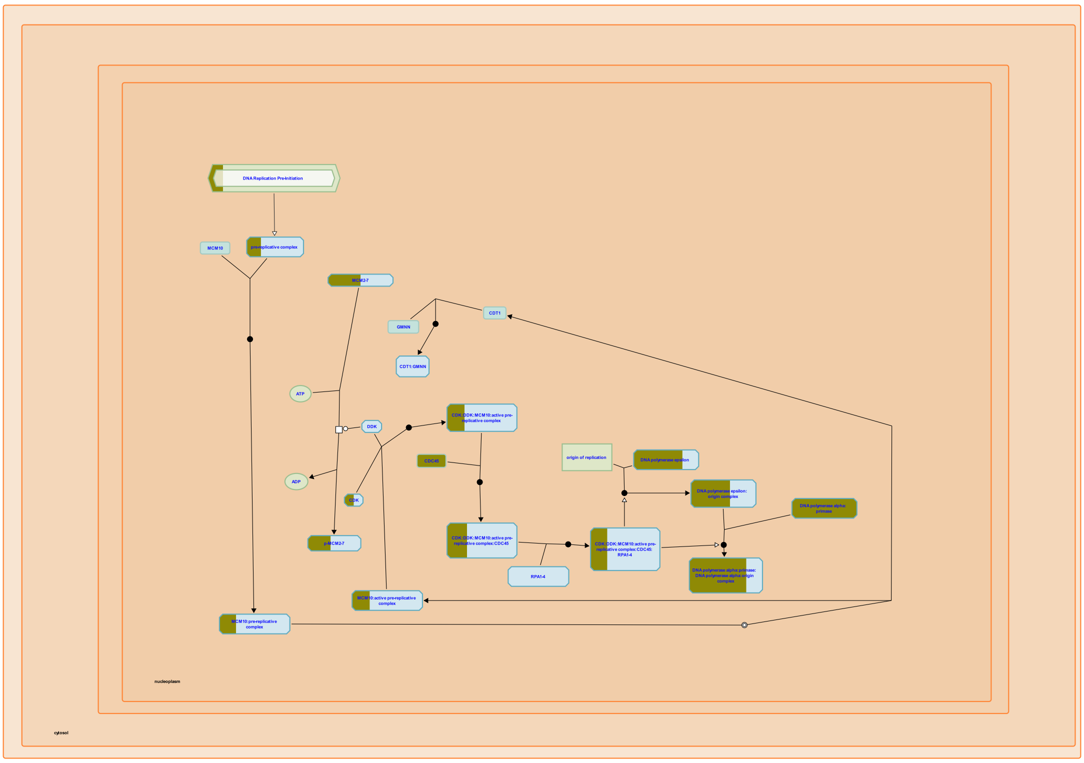


reactome

**Cellular compartments:** nucleoplasm.

In S. cerevisiae, two ORC subunits, Orc1 and Orc5, both bind ATP, and Orc1 in addition has ATPase activity. Both ATP binding and ATP hydrolysis appear to be essential functions in vivo. ATP binding by Orc1 is unaffected by the association of ORC with origin DNA (ARS) sequences, but ATP hydrolysis is ARS-dependent, being suppressed by associated double-stranded DNA and stimulated by associated single-stranded DNA. These data are consistent with the hypothesis that ORC functions as an ATPase switch, hydrolyzing bound ATP and changing state as DNA unwinds at the origin immediately before replication. It is attractive to speculate that ORC likewise functions as a switch as human pre-replicative complexes are activated, but human Orc proteins are not well enough characterized to allow the model to be critically tested. mRNAs encoding human orthologs of all six Orc proteins have been cloned, and ATP-binding amino acid sequence motifs have been identified in Orc1, Orc4, and Orc5. Interactions among proteins expressed from the cloned genes have been characterized, but the ATP-binding and hydrolyzing properties of these proteins and complexes of them have not been determined.

#### Edit history

| Date | Action | Author |
|---|---|---|
| 2003-06-05 | Created | Davey MJ, O'Donnell M |
| 2023-08-26 | Modified | Wright A |

#### 12 submitted entities found in this pathway, mapping to 13 Reactome entities

| Input | UniProt Id | Input | UniProt Id | Input | UniProt Id |
|---|---|---|---|---|---|
| CDC45 | O75419 | CDK2 | P24941 | MCM2 | P33993, P49736 |
| MCM3 | P25205 | ORC3 | Q9UBD5 | POLA1 | P09884 |
| POLA2 | Q14181 | POLE | Q07864 | POLE3 | Q9NRF9 |
| POLE4 | Q9NR33 | PRIM1 | P49642 | PRIM2 | P49643 |

#### Interactors found in this pathway (10)

| Input | UniProt Id | Interacts with | Input | UniProt Id | Interacts with |
|---|---|---|---|---|---|
| CCNA2 | P20248 | O75496, Q9H211 | CDK1 | P06493 | Q9H211 |
| CDK2 | P24941 | O75496, Q9H211 | CEP55 | Q53EZ4 | O75419 |
| KANK2 | Q63ZY3 | O75496 | MCM2 | P49736 | O75419 |
| RAB4B | P61018 | O75419 | SAPCD2 | Q86UD0 | O75496 |
| TFDP1 | Q14186 | O75496 | TOPBP1 | Q92547 | O75419 |

#### 19. Resolution of AP sites via the multiple-nucleotide patch replacement pathway (R-HSA-110373)

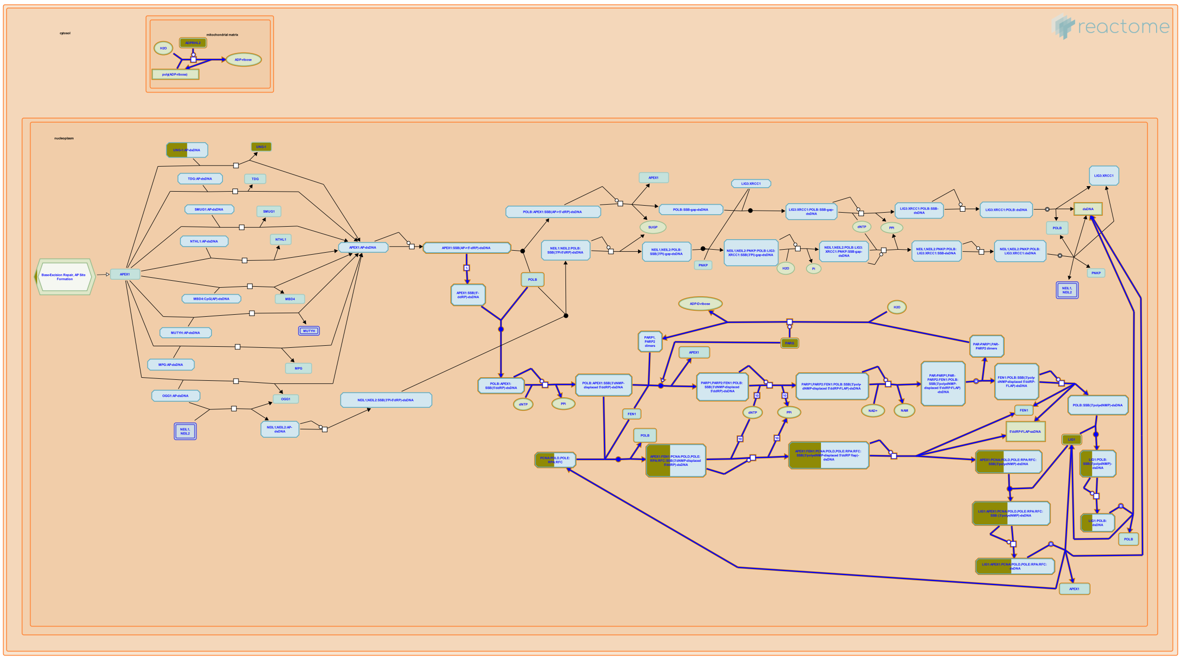

**Cellular compartments:** nucleoplasm.

While the single nucleotide replacement pathway appears to facilitate the repair of most damaged bases, an alternative BER pathway is evoked when the structure of the 5'-terminal sugar phosphate is such that it cannot be cleaved through the AP lyase activity of DNA polymerase beta (POLB). Under these circumstances, a short stretch of residues containing the abasic site is excised and replaced (Dianov et al., 1999). Following DNA glycosylase-mediated cleavage of the damaged base, the endonuclease APEX1 is recruited to the site of damage where it cleaves the 5' side of the abasic deoxyribose residue, as in the single nucleotide replacement pathway. However, POLB then synthesizes the first replacement residue without prior cleavage of the 5'-terminal sugar phosphate, hence displacing this entity. Long-patch BER can be completed by continued POLB-mediated DNA strand displacement synthesis in the presence of PARP1 or PARP2, FEN1 and DNA ligase I (LIG1) (Prasad et al. 2001). When the PCNA-containing replication complex is available, as is the case with cells in the S-phase of the cell cycle, DNA strand displacement synthesis is catalyzed by DNA polymerase delta (POLD) or DNA polymerase epsilon (POLE) complexes, in the presence of PCNA, RPA, RFC, APEX1, FEN1 and LIG1 (Klungland and Lindahl 1997, Dianova et al. 2001). In both POLB-dependent and PCNA-dependent DNA displacement synthesis, the displaced DNA strand containing the abasic sugar phosphate creates a flap structure that is recognized and cleaved by the flap endonuclease FEN1. The replacement residues added by POLB or POLD/POLE are then ligated by the DNA ligase I (LIG1) (Klungland and Lindahl, 1997; Matsumoto et al., 1999).

#### Edit history

| Date | Action | Author |
|---|---|---|
| 2004-01-29 | Created | Matthews L |
| 2004-02-03 | Edited | Matthews L |
| 2004-02-09 | Authored | Matthews L |
| 2014-12-04 | Revised | Orlic-Milacic M |
| 2014-12-04 | Edited | Orlic-Milacic M |
| 2014-12-22 | Reviewed | Borowiec JA |
| 2023-08-26 | Modified | Wright A |

#### 10 submitted entities found in this pathway, mapping to 11 Reactome entities

| Input | UniProt Id | Input | UniProt Id | Input | UniProt Id |
|---|---|---|---|---|---|
| ADPRHL2 | Q9NX46 | LIG1 | P18858 | PARG | Q86W56 |
| POLD1 | P28340 | POLD2 | P49005 | POLE | Q07864 |
| POLE3 | Q9NRF9 | POLE4 | Q9NR33 | RFC1 | P35251 |
| RFC5 | P40937, P40938 | | | | |

#### 20. PCNA-Dependent Long Patch Base Excision Repair (R-HSA-5651801)

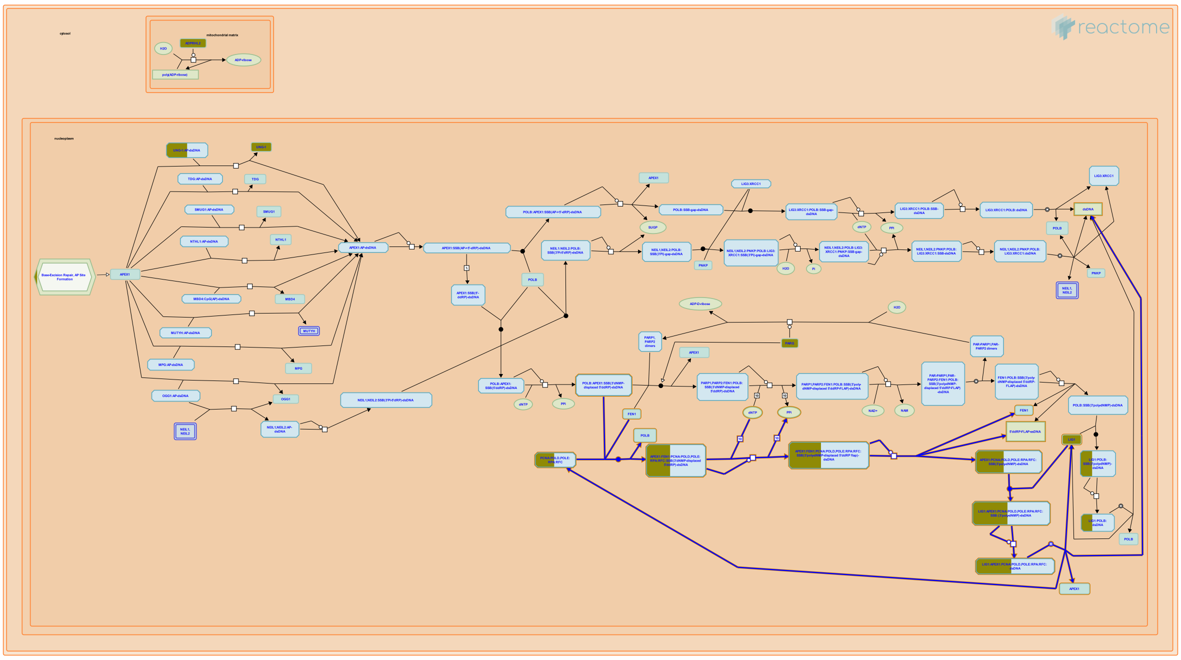

**Cellular compartments:** nucleoplasm.

Long-patch base excision repair (BER) can proceed through PCNA-dependent DNA strand displacement synthesis by replicative DNA polymerases - DNA polymerase delta complex (POLD) or DNA polymerase epsilon (POLE) complex. The PCNA-dependent branch of long-patch BER may occur in cells in the S phase of the cell cycle, when the replication complexes that contain PCNA, POLD or POLE, RPA and RFC are available. POLB incorporates the first nucleotide at the 3'-end of APEX1-generated single strand break (SSB), thus displacing the damaged AP (abasic) dideoxyribose phosphate residue at the 5'-end of SSB (5'ddRP). PCNA is recruited to BER sites by APEX1 and flap endonuclease FEN1, and loaded onto damaged DNA by RFC. POLD and POLE in complex with PCNA continue the displacement DNA strand synthesis. FEN1 cleaves the displaced DNA strand with the AP residue (5'ddRP), and DNA ligase I (LIG1) ligates the multiple nucleotide patch at the 3' end of the SSB with the FEN1-processed 5'-end of the SSB (Klungland and Lindahl 1997, Stucki et al. 1998, Dianov et al. 1999, Matsumoto et al. 1999, Podlutsky et al. 2001, Dianova et al. 2001, Ranalli et al. 2002).

#### Edit history

| Date | Action | Author |
|---|---|---|
| 2014-11-25 | Created | Orlic-Milacic M |
| 2014-12-04 | Edited | Orlic-Milacic M |
| 2014-12-04 | Authored | Orlic-Milacic M |
| 2014-12-22 | Reviewed | Borowiec JA |
| 2023-08-26 | Modified | Wright A |

#### 8 submitted entities found in this pathway, mapping to 9 Reactome entities

| Input | UniProt Id | Input | UniProt Id | Input | UniProt Id |
|---|---|---|---|---|---|
| LIG1 | P18858 | POLD1 | P28340 | POLD2 | P49005 |
| POLE | Q07864 | POLE3 | Q9NRF9 | POLE4 | Q9NR33 |
| RFC1 | P35251 | RFC5 | P40937, P40938 | | |

#### 21. Processive synthesis on the lagging strand (R-HSA-69183)

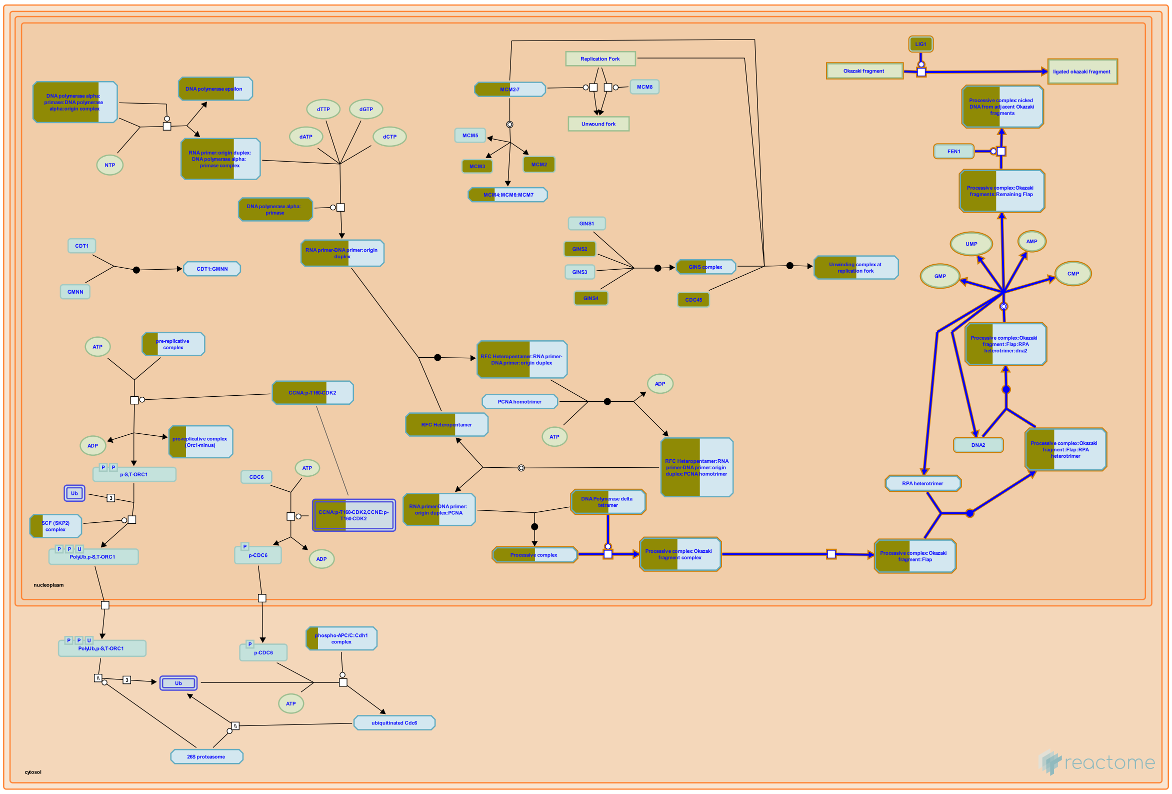


**Cellular compartments:** nucleoplasm.

The key event that allows the processive synthesis on the lagging strand, is polymerase switching from pol alpha to pol delta, as on the leading strand. However, the processive synthesis on the lagging strand proceeds very differently. DNA synthesis is discontinuous, and involves the formation of short fragments called the Okazaki fragments. During the synthesis of Okazaki fragments, the RNA primer is folded into a single-stranded flap, which is removed by endonucleases. This is followed by the ligation of adjacent Okazaki fragments.


##### Edit history

| Date | Action | Author |
|---|---|---|
| 2023-08-26 | Modified | Wright A |

##### 7 submitted entities found in this pathway, mapping to 7 Reactome entities

| Input | UniProt Id | Input | UniProt Id | Input | UniProt Id |
|---|---|---|---|---|---|
| LIG1 | P18858 | POLA1 | P09884 | POLA2 | Q14181 |
| POLD1 | P28340 | POLD2 | P49005 | PRIM1 | P49642 |
| PRIM2 | P49643 | | | | |

#### 22. tRNA processing in the mitochondrion (R-HSA-6785470)

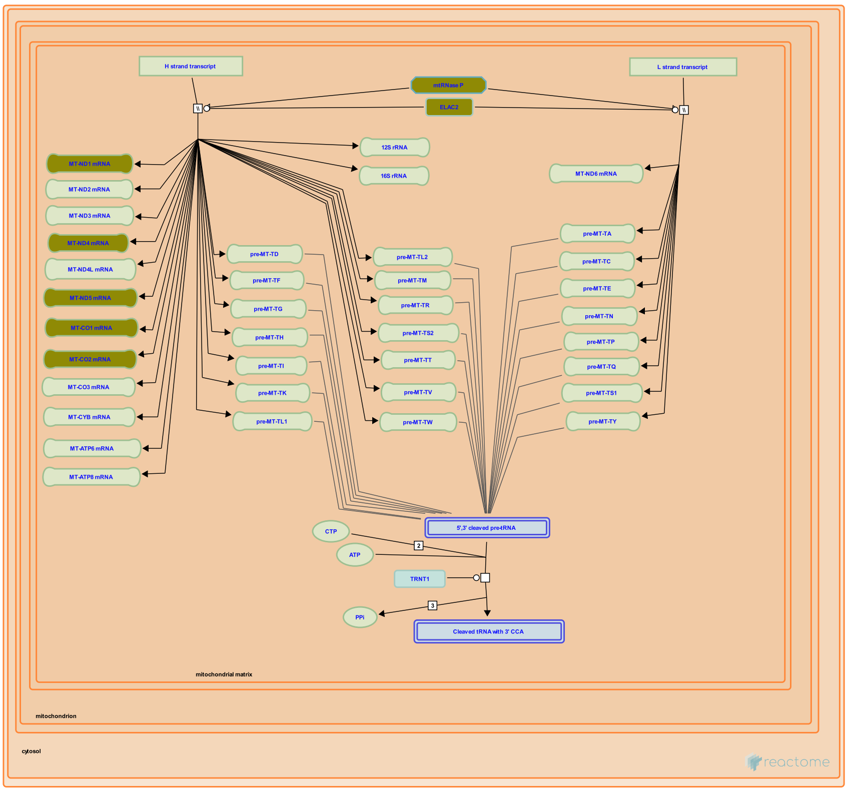


Each strand of the circular mitochondrial genome is transcribed to yield long polycistronic transcripts, the heavy strand transcript and the light strand transcript, which are then cleaved to yield tRNAs, rRNAs, and mRNAs (Mercer et al. 2011, reviewed in Suzuki et al. 2011, Rossmanith 2012, Powell et al. 2015). Mitochondrial RNase P, which is completely distinct from nuclear RNase P in having different protein subunits and no RNA component, cleaves at the 5' ends of tRNAs. RNase Z, an isoform of ELAC2 in mitochondria, cleaves at the 3' ends of tRNAs. (A different isoform of ELAC2 serves as RNase Z in the nucleus.) Unknown nucleases make additional cleavages near the 5' end of MT-CO3, the 5' end of CO1, the 5' end of CYB, and the 3' end of ND6. TRNT1 (CCA-adding enzyme) then post-transcriptionally polymerizes the universal acceptor sequence CCA onto the 3' ends of the cleaved tRNAs. In yeast, plants, and protozoa additional tRNAs encoded in the nucleus are imported into mitochondria from the cytosol (reviewed in Schneider 2011), however human mitochondria encode a complete complement of 22 tRNAs required for translation and tRNA import has not been observed in mammals. Mutations that affect mitochondrial tRNA processing cause human diseases that are generally characterized by abnormalities in energy-requiring tissues such as brain and muscle (reviewed in Suzuki et al. 2011, Sarin and Leidel 2014).

#### Edit history

| Date | Action | Author |
|---|---|---|
| 2015-06-28 | Edited | May B |
| 2015-06-28 | Authored | May B |
| 2015-06-28 | Created | May B |
| 2015-08-11 | Reviewed | Levinger L |
| 2015-08-25 | Reviewed | Motorin Y |
| 2015-10-24 | Reviewed | Jarrous N |
| 2015-11-07 | Reviewed | Powell CA |
| 2015-11-18 | Reviewed | Suzuki T |
| 2023-08-31 | Modified | Wright A |

#### 9 submitted entities found in this pathway, mapping to 9 Reactome entities

| Input | UniProt Id |
|---|---|
| ELAC2 | Q9BQ52 |
| KIAA0391 | O15091 |

| Input | UniProt Id |
|---|---|
| HSD17B10 | Q99714 |
| TRMT10C | Q7L0Y3 |

| Input | Ensembl Id |
|---|---|
| MT-CO1 | ENST00000361624 |
| MT-ND4 | ENST00000361381 |

| Input | Ensembl Id |
|---|---|
| MT-CO2 | ENST00000361739 |
| MT-ND5 | ENST00000361567 |

| Input | Ensembl Id |
|---|---|
| MT-ND1 | ENST00000361390 |

#### 23. Polymerase switching on the C-strand of the telomere (R-HSA-174411)

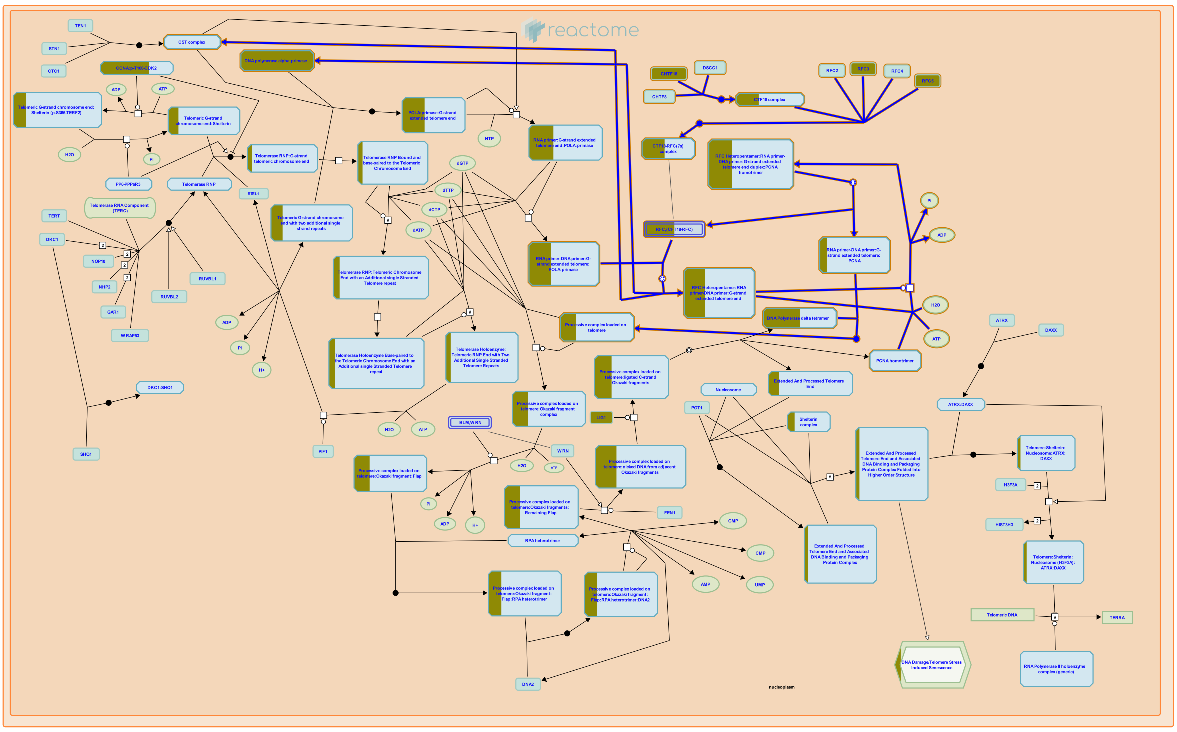

**Cellular compartments:** nucleoplasm.

After the primers are synthesized on the G-Rich strand, Replication Factor C binds to the 3'-end of the initiator DNA to trigger polymerase switching. The non-processive nature of pol alpha catalytic activity and the tight binding of Replication Factor C to the primer-template junction presumably lead to the turnover of the pol alpha:primase complex. After the Pol alpha-primase primase complex is displaced from the primer, the proliferating cell nuclear antigen (PCNA) binds to form a "sliding clamp" structure. Replication Factor C then dissociates, and DNA polymerase delta binds and catalyzes the processive synthesis of DNA.


##### Edit history

| Date | Action | Author |
|---|---|---|
| 2006-02-17 | Created | Gillespie ME |
| 2006-03-10 | Authored | Seidel J, Blackburn EH |
| 2006-07-13 | Reviewed | Price C |

| Date | Action | Author |
|---|---|---|
| 2009-06-03 | Revised | D'Eustachio P |
| 2019-12-02 | Revised | Orlic-Milacic M |
| 2020-04-29 | Reviewed | Hayashi MT |
| 2020-05-04 | Edited | Orlic-Milacic M |
| 2023-08-26 | Modified | Wright A |

#### 10 submitted entities found in this pathway, mapping to 11 Reactome entities

| Input | UniProt Id | Input | UniProt Id | Input | UniProt Id |
|---|---|---|---|---|---|
| CHTF18 | Q8WVB6 | POLA1 | P09884 | POLA2 | Q14181 |
| POLD1 | P28340 | POLD2 | P49005 | PRIM1 | P49642 |
| PRIM2 | P49643 | RFC1 | P35251 | RFC5 | P40937, P40938 |
| TPP1 | Q96AP0 | | | | |

#### Interactors found in this pathway (2)

| Input | UniProt Id | Interacts with | Input | UniProt Id | Interacts with |
|---|---|---|---|---|---|
| RFC1 | P35251 | P35250, P40938, P40937, P35249 | RFC5 | P40937 | P35249 |

#### 24. Recognition of DNA damage by PCNA-containing replication complex (R-HSA-110314)

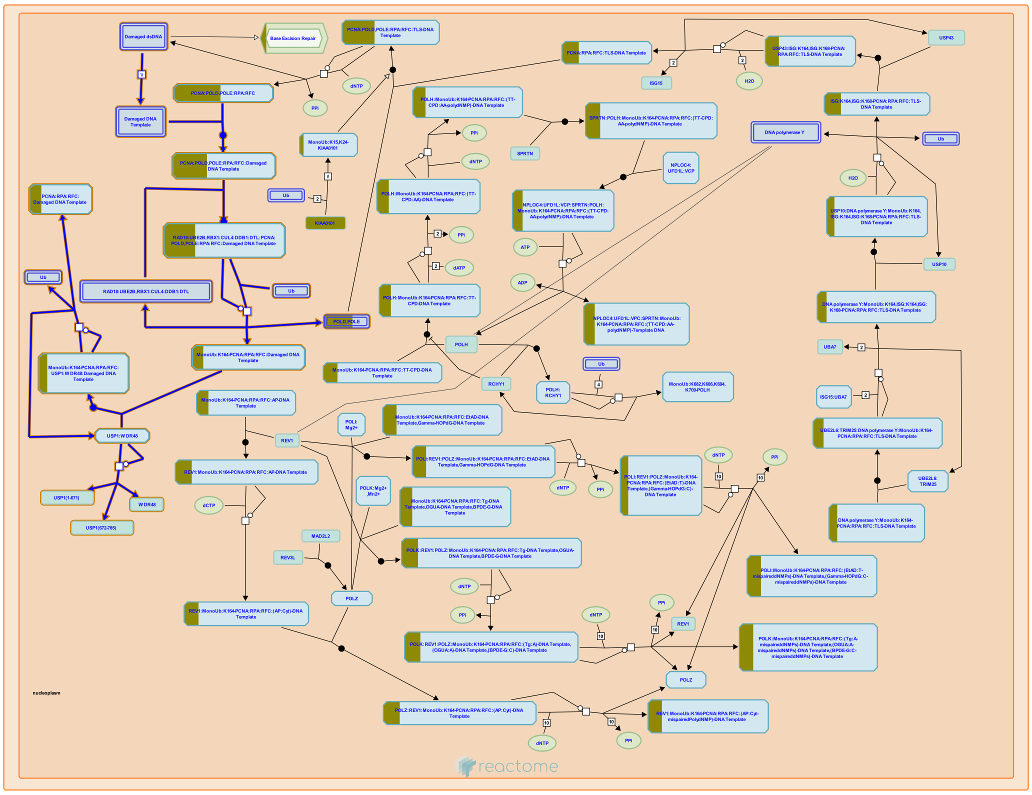

**Cellular compartments:** nucleoplasm.

Damaged double strand DNA (dsDNA) cannot be successfully used as a template by replicative DNA polymerase delta (POLD) and epsilon (POLE) complexes (Hoege et al. 2002). When the replication complex composed of PCNA, RPA, RFC and POLD or POLE stalls at a DNA damage site, PCNA becomes monoubiquitinated by RAD18 bound to UBE2B (RAD6). POLD or POLE dissociate from monoubiquitinated PCNA, while Y family DNA polymerases - REV1, POLH (DNA polymerase eta), POLK (DNA polymerase kappa) and POLI (DNA polymerase iota) - bind monoubiquitinated PCNA through their ubiquitin binding and PCNA binding motifs, resulting in a polymerase switch and initiation of translesion synthesis (TLS) (Hoege et al. 2002, Friedberg et al. 2005).


##### Edit history

| Date | Action | Author |
|---|---|---|
| 2004-01-29 | Created | Gopinathrao G |
| 2014-12-11 | Edited | Orlic-Milacic M |
| 2014-12-11 | Authored | Orlic-Milacic M |

| Date | Action | Author |
|---|---|---|
| 2015-01-07 | Reviewed | Borowiec JA |
| 2023-08-26 | Modified | Wright A |

#### 7 submitted entities found in this pathway, mapping to 8 Reactome entities

| Input | UniProt Id | Input | UniProt Id | Input | UniProt Id |
|---|---|---|---|---|---|
| POLD1 | P28340 | POLD2 | P49005 | POLE | Q07864 |
| POLE3 | Q9NRF9 | POLE4 | Q9NR33 | RFC1 | P35251 |
| RFC5 | P40937, P40938 | | | | |

#### 25. Dual incision in TC-NER (R-HSA-6782135)

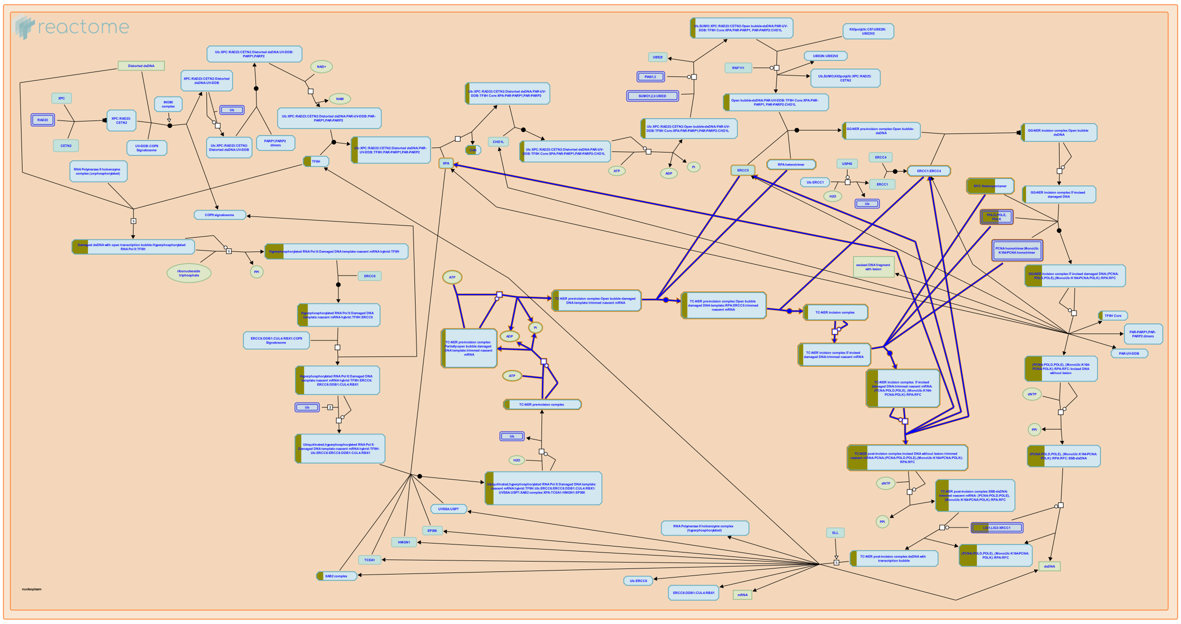

**Cellular compartments:** nucleoplasm.

In transcription-coupled nucleotide excision repair (TC-NER), similar to global genome nucleotide excision repair (GG-NER), the oligonucleotide that contains the lesion is excised from the open bubble structure via dual incision of the affected DNA strand. 5' incision by the ERCC1:ERCC4 (ERCC1:XPF) endonuclease precedes 3' incision by ERCC5 (XPG) endonuclease. In order for the TC-NER pre-incision complex to assemble and the endonucleases to incise the damaged DNA strand, the RNA polymerase II (RNA Pol II) complex has to backtrack - reverse translocate from the damage site. Although the mechanistic details of this process are largely unknown in mammals, it may involve ERCC6/ERCC8-mediated chromatin remodelling/ubiquitination events, the DNA helicase activity of the TFIIH complex and TCEA1 (TFIIS)-stimulated cleavage of the 3' protruding end of nascent mRNA by RNA Pol II (Donahue et al. 1994, Lee et al. 2002, Sarker et al. 2005, Vermeulen and Fousteri 2013, Hanawalt and Spivak 2008, Staresincic et al. 2009, Epshtein et al. 2014).


##### Edit history

| Date | Action | Author |
|---|---|---|
| 2004-01-29 | Authored | Hoeijmakers JH |
| 2004-02-11 | Authored | Gopinathrao G |
| 2015-05-28 | Edited | Orlic-Milacic M |
| 2015-06-04 | Created | Orlic-Milacic M |
| 2015-06-16 | Revised | Orlic-Milacic M |
| 2015-06-16 | Authored | Orlic-Milacic M |
| 2015-08-03 | Reviewed | Fousteri M |
| 2023-08-31 | Modified | Wright A |

#### 11 submitted entities found in this pathway, mapping to 12 Reactome entities

| Input | UniProt Id | Input | UniProt Id | Input | UniProt Id |
|---|---|---|---|---|---|
| AQR | O60306 | CCNH | P51946 | CDK7 | P50613 |
| GTF2H3 | Q13889 | POLD1 | P28340 | POLD2 | P49005 |
| POLE | Q07864 | POLE3 | Q9NRF9 | POLE4 | Q9NR33 |
| RFC1 | P35251 | RFC5 | P40937, P40938 | | |

#### Interactors found in this pathway (3)

| Input | UniProt Id | Interacts with | Input | UniProt Id | Interacts with |
|---|---|---|---|---|---|
| CDK7 | P50613 | P28715 | HERC2 | O95714 | P23025 |
| TRIM27 | P14373 | P23025 | | | |

### 6. Identifiers found

Below is a list of the input identifiers that have been found or mapped to an equivalent element in Reactome, classified by resource.

##### 533 of the submitted entities were found, mapping to 662 Reactome entities

| Input | UniProt Id | Input | UniProt Id | Input | UniProt Id |
|---|---|---|---|---|---|
| AAMP | Q13685 | AARS2 | Q5JTZ9 | ABAT | P80404 |
| ABCB7 | Q9NWR8 | ABL2 | P42684 | ACAD9 | Q9H845 |
| ACADM | P11310 | ACADVL | P49748 | ACO2 | Q99798 |
| ACOT1 | Q86TX2 | ACOT9 | Q9Y305 | ACP2 | P15309 |
| ACSF3 | Q4G176 | ADD1 | P35611 | ADPRHL2 | Q9NX46 |
| ADRBK1 | P25098 | AGA | P20933 | AGMAT | Q9BSE5 |
| AGO1 | Q9H9G7, Q9HCK5, Q9UL18 | AIFM2 | Q9BRQ8 | AK6 | Q9Y3D8 |
| AKR1C1 | Q04828 | AKR1C2 | P52895 | AKR1C3 | P42330 |
| ALCAM | Q13740 | ALDH1B1 | P30837 | ALDH6A1 | Q02252 |
| ALG9 | Q9H6U8 | ALKBH5 | Q6P6C2 | ANLN | Q9NQW6 |
| ANXA1 | P04083 | APOOL | Q6UXV4 | AQR | O60306 |
| ARAP1 | Q96P48 | ARHGEF11 | O15085 | ARHGEF2 | Q92974 |
| ASNS | P08243 | ASS1 | P00966 | ATAD2 | Q6PL18 |
| ATP5H | O75947 | ATP5J2 | P56134 | ATP5L | O75964 |
| ATP7A | Q04656 | AURKA | O14965 | AURKB | Q96GD4 |
| AZGP1 | P25311 | BCAS2 | O75934 | BCKDHB | P21953 |
| BCKDK | O14874 | BCS1L | Q9Y276 | BDH1 | Q02338 |
| BMP7 | P18075 | BRIP1 | Q9P0J6 | C1QBP | Q07021 |
| C2orf47 | Q8WWC4 | C2orf49 | Q9BVC5 | CA5B | P35218, Q9Y2D0 |
| CALML5 | Q9NZT1 | CAPN2 | P17655 | CARKD | Q8IW45 |
| CARS | P49589 | CAT | P04040 | CBS | P35520 |
| CCNA2 | P20248 | CCNB1 | P14635 | CCNB2 | O95067 |
| CCND1 | P24385 | CCNH | P51946 | CD47 | Q08722 |
| CD55 | P08174 | CD59 | P13987 | CDC20 | Q12834 |
| CDC23 | Q9UJX2 | CDC45 | O75419 | CDCA5 | Q96FF9 |
| CDCA8 | Q53HL2 | CDK1 | P06493 | CDK2 | P24941 |
| CDK7 | P50613 | CEBPB | P17676 | CEP55 | Q92753 |
| CERS1 | P27544 | CHCHD2 | Q9Y6H1 | CHDH | Q8NE62 |
| CHTF18 | Q8WVB6 | CHTOP | Q9Y3Y2 | CIT | O14578, O14578-3 |
| CKB | P12277 | CLCN7 | P51798 | COL18A1 | P39060 |
| COMMD8 | Q9NX08 | COQ5 | Q5HYK3 | COQ6 | Q9Y2Z9 |
| CORO1A | P31146 | COX11 | Q9Y6N1 | COX15 | Q7KZN9 |
| COX16 | Q9P0S2 | COX4I1 | P13073 | COX7A2L | O14548 |
| CPOX | P36551 | CRABP1 | P29762 | CST3 | P01034 |
| CYBA | P13498 | D2HGDH | Q8N465 | DAP3 | P51398 |
| DARS2 | Q6PI48 | DBT | P11182 | DEGS1 | O15121 |
| DHFR | P00374 | DHTKD1 | Q96HY7 | DLAT | P10515 |
| DLGAP5 | Q15398 | DNMT1 | P26358 | DNPH1 | O43598 |
| DPH1 | Q9BZG8 | DPH2 | Q9BQC3 | DPH7 | Q9BTV6 |
| DPP7 | Q9UHL4 | DSC1 | Q08554 | DYNLL1 | P63167 |
| EARS2 | Q5JPH6 | ECH1 | Q13011 | ECHS1 | P30084 |

| Input | UniProt Id | Input | UniProt Id | Input | UniProt Id |
|---|---|---|---|---|---|
| ECSIT | Q9BQ95 | ELAC2 | Q9BQ52 | ELP3 | Q9H9T3 |
| EPS15 | P42566 | ERAL1 | O75616 | ERCC6L | Q2NKX8 |
| ETNK1 | Q9HBU6 | EZH2 | Q15910 | FANCD2 | Q9BXW9 |
| FANCI | Q9NVI1 | FASTKD2 | Q9NYY8 | FBXL15 | Q9H469 |
| FBXO21 | O94952 | FCF1 | Q9Y324 | FDFT1 | P37268 |
| FDX1 | P10109 | FECH | P22830 | FHL2 | Q14192 |
| FN1 | P02751 | FNBP1L | Q5T0N5 | FPGS | Q05932-1, Q05932-2 |
| FTH1 | P02794 | FUCA1 | P04066 | FUK | Q8N0W3 |
| FUNDC1 | Q8IVP5 | FXYD3 | Q14802 | GABARAPL2 | P60520 |
| GADD45GIP1 | Q8TAE8 | GARS | P41250 | GCDH | Q92947 |
| GFM1 | Q96RP9 | GGCX | P38435 | GINS2 | Q9Y248 |
| GINS4 | Q9BRT9 | GLS | O94925 | GOLGA1 | Q92805 |
| GPT2 | P24298, Q8TD30-1 | GPX4 | P36969 | GSN | P06396 |
| GSTK1 | Q9Y2Q3 | GTF2H3 | Q13889 | HADH | P40939, Q16836 |
| HADHA | P40939 | HADHB | P55084 | HDHD1 | Q08623 |
| HERC2 | O95714 | HINT2 | Q9BX68 | HMGCR | P04035-1 |
| HMOX1 | P09601 | HSD17B10 | Q99714 | HSPA5 | P11021 |
| ICMT | O60725 | ICT1 | Q14197 | IDH1 | P51553 |
| IDH3G | P51553 | IFI30 | P13284 | IGFBP2 | P18065 |
| IQGAP3 | Q86VI3 | ISCA1 | Q9BUE6 | ITGA5 | P08648 |
| JAK1 | P23458 | JMJD6 | Q6NYC1 | JMY | Q8N9B5 |
| JUNB | P17275 | KAT8 | Q9H7Z6 | KDM3A | Q9Y4C1 |
| KDM5C | P41229, Q9BY66 | KIAA0101 | Q15004 | KIAA0391 | O15091 |
| KIF11 | P52732 | KIF15 | Q9NS87 | KIF1A | Q12756 |
| KIF20A | O95235 | KIF22 | Q14807 | KIF23 | Q02241 |
| KIF2C | Q99661 | KIF4A | O95239 | KIFAP3 | Q92845 |
| KIFC1 | Q9BW19 | KLC4 | Q9NSK0 | KNTC1 | P50748 |
| KPNA2 | P52292 | L2HGDH | Q9H9P8 | LARS2 | Q15031 |
| LDLR | P98164 | LETM1 | O95202 | LGALS3 | P17931 |
| LGALS3BP | Q08380 | LHPP | Q9H008 | LIG1 | P18858 |
| LOXL2 | Q9Y4K0 | LRP10 | Q7Z4F1 | LSM1 | O15116 |
| MAD2L1 | Q13257 | MADD | Q8WXG6-3 | MAN1A1 | P33908 |
| MAN2C1 | Q9NTJ4 | MANBA | O00462 | MARCKS | P29966 |
| MB21D1 | Q8N884 | MCCC2 | Q9HCC0 | MCM2 | P33993, P49736 |
| MCM3 | P25205 | MDC1 | Q14676 | ME2 | P23368 |
| MECP2 | P51608-1, P51608-2 | MGA | O43451 | MIA3 | Q5JRA6 |
| MICU1 | Q9BPX6 | MICU2 | Q8IYU8 | MIOS | Q9NXC5 |
| MIS18A | Q9NYP9 | MLH1 | P40692 | MMADHC | Q9H3L0 |
| MORF4L1 | Q9UBU8 | MRPL1 | Q9BYD6 | MRPL10 | Q7Z7H8 |
| MRPL11 | Q9Y3B7 | MRPL12 | P52815 | MRPL13 | Q9BYD1 |
| MRPL14 | Q6P1L8 | MRPL15 | P49406, Q9P015 | MRPL16 | Q9NX20 |
| MRPL17 | Q9NRX2 | MRPL18 | Q9H0U6 | MRPL19 | P49406 |
| MRPL2 | Q5T653, Q9BZE1 | MRPL20 | Q9BYC9 | MRPL21 | Q7Z2W9 |
| MRPL22 | Q9NWU5 | MRPL23 | Q16540 | MRPL24 | Q96A35 |
| MRPL27 | Q8IXM3, Q9P0M9 | MRPL28 | Q13084, Q8TCC3 | MRPL3 | P09001 |
| MRPL32 | Q6P1L8, Q9BYC8 | MRPL34 | Q9BQ48 | MRPL37 | Q9BZE1 |
| MRPL38 | Q96DV4 | MRPL39 | Q9NYK5 | MRPL4 | Q9BYD3 |
| MRPL40 | Q9NQ50 | MRPL41 | Q8IXM3 | MRPL42 | Q9Y6G3 |
| MRPL44 | Q9H9J2 | MRPL45 | Q9BRJ2 | MRPL46 | Q9H2W6 |

| Input | UniProt Id | Input | UniProt Id | Input | UniProt Id |
|---|---|---|---|---|---|
| MRPL47 | Q9HD33 | MRPL48 | Q96GC5 | MRPL49 | Q13405 |
| MRPL51 | Q4U2R6 | MRPL52 | Q86TS9 | MRPL53 | Q96EL3 |
| MRPL54 | Q6P161 | MRPL55 | Q7Z7F7 | MRPL9 | Q9BYD2 |
| MRPS10 | P82664 | MRPS11 | P82912 | MRPS12 | O15235 |
| MRPS14 | O60783 | MRPS15 | P82914 | MRPS16 | Q9Y3D3 |
| MRPS17 | Q9Y2R5 | MRPS18A | Q9NVS2 | MRPS18B | Q9Y676 |
| MRPS18C | Q9Y3D5 | MRPS2 | Q9Y399 | MRPS21 | P82921 |
| MRPS22 | P82650 | MRPS23 | Q9Y3D9 | MRPS24 | Q96EL2 |
| MRPS25 | P82663 | MRPS26 | Q9BYN8 | MRPS27 | Q92552 |
| MRPS28 | P82673, Q9Y2Q9 | MRPS30 | Q9NP92 | MRPS31 | Q92665 |
| MRPS33 | Q9Y291 | MRPS34 | P82930 | MRPS35 | P82673, Q9Y2Q9 |
| MRPS5 | P82675 | MRPS6 | P82932 | MRPS7 | Q9Y2R9 |
| MRPS9 | P82933 | MRRF | Q96E11 | MSMO1 | Q15800 |
| MT-CO1 | P00395 | MT-CO2 | P00403 | MT-ND1 | P03886 |
| MT-ND4 | P03905 | MT-ND5 | P03915 | MTERF3 | Q96E29 |
| MTERF4 | Q7Z6M4 | MTHFD2 | P13995 | MTIF2 | P46199 |
| MTIF3 | Q9H2K0 | MTMR1 | Q13613 | MUC1 | P15941 |
| MUT | P22033 | NARS2 | Q96I59 | NBR1 | Q14596 |
| NCAPD2 | Q15021 | NCAPD3 | P42695 | NCAPG2 | Q86XI2 |
| NDOR1 | Q9UHB4 | NDRG1 | Q92597 | NDUFA10 | O95299 |
| NDUFA12 | Q9UI09 | NDUFA13 | Q9P0J0 | NDUFA2 | O43678 |
| NDUFA3 | O95167 | NDUFA4 | O00483 | NDUFA5 | Q16718 |
| NDUFA6 | P56556 | NDUFA9 | Q16795 | NDUFAB1 | O14561 |
| NDUFAF1 | Q9Y375 | NDUFAF2 | Q8N183 | NDUFAF4 | Q9P032 |
| NDUFB10 | O96000 | NDUFB11 | Q9NX14 | NDUFB4 | O95168 |
| NDUFB6 | O95139 | NDUFB9 | Q9Y6M9 | NDUFS1 | P28331 |
| NDUFS2 | O75306 | NDUFS3 | O75489 | NDUFS4 | O43181 |
| NDUFS5 | O43920 | NDUFS7 | O75251 | NDUFS8 | O00217 |
| NDUFV1 | P49821 | NDUFV2 | P19404 | NFS1 | Q9Y697-1 |
| NHP2L1 | P55769 | NQO1 | P15559 | NR2F6 | P10588 |
| NSMCE4A | Q8N140, Q9NXX6 | OAT | P04181 | OGDH | Q02218 |
| OGFOD1 | Q8N543 | ORC3 | Q9UBD5 | OXA1L | Q15070 |
| P4HTM | Q9NNW5 | PAM16 | Q9Y3D7 | PARG | Q86W56 |
| PC | P11498 | PDCD4 | P15848 | PDHA1 | P08559, P29803 |
| PDHB | P11177 | PDK3 | Q15120 | PDP1 | Q9P0J1 |
| PDPR | Q8NCN5 | PDZD11 | Q5EBL8 | PEX3 | P56589 |
| PFDN5 | Q99471 | PHGDH | O43175 | PKMYT1 | Q99640 |
| PKP1 | Q13835 | PKP4 | Q99569 | PLIN2 | Q99541 |
| PLK1 | P53350 | PLOD1 | Q02809 | PNPLA2 | Q96AD5 |
| PNPT1 | Q8TCS8 | POFUT2 | Q9Y2G5 | POLA1 | P09884 |
| POLA2 | Q14181 | POLD1 | P28340 | POLD2 | P49005 |
| POLE | Q07864 | POLE3 | Q9NRF9 | POLE4 | Q9NR33 |
| POLG | P27958 | POLR3A | O14802 | POLRMT | O00411 |
| PPAP2C | O43688 | PPAT | Q06203 | PPP2R1B | P30154 |
| PPP2R5C | Q13362 | PRC1 | O43663 | PRIM1 | P49642 |
| PRIM2 | P49643 | PRKACG | P22612 | PRMT7 | Q9NVM4 |
| PRPS1 | P60891 | PRR11 | Q86W47 | PSAP | Q8IWL1 |
| PSPH | P78330 | PTCD3 | Q96EY7 | PTPMT1 | Q8WUK0 |
| PTPN9 | P43378 | PXDN | Q92626 | PXK | Q8NC69 |

| Input | UniProt Id | Input | UniProt Id | Input | UniProt Id |
|---|---|---|---|---|---|
| PYCR1 | P32322 | PYCR2 | Q53H96, Q96C36 | QIL1 | Q5XKP0 |
| RAB25 | P57735 | RAB2B | Q8WUD1 | RAB4B | P61018 |
| RACGAP1 | Q9H0H5 | RAD51C | O43502 | RAF1 | P04049 |
| RARS2 | Q5T160 | RB1 | P06400 | RBP1 | O95153 |
| RCN1 | Q15293 | RFC1 | P35251 | RFC5 | P40937, P40938 |
| RIF1 | Q5UIP0 | RIOK2 | Q9BVS4 | RMI2 | Q96E14 |
| RNASEL | Q05823 | RNF14 | Q9UBS8 | RNMTL1 | Q9HC36 |
| RPL26L1 | Q9UNX3 | RPLP1 | P05386 | RPS27L | Q71UM5 |
| RRM1 | P23921 | RRM2 | P31350 | RRN3 | Q9NYV6 |
| S100P | P25815, P80511 | SCCPDH | Q8NBX0 | SDHA | P31040 |
| SDHB | P21912 | SDHC | Q99643 | SEMG1 | P04279 |
| SEPHS2 | Q99611 | SESN2 | P58004 | SF3B5 | Q9BWJ5 |
| SGSH | P51688 | SH3KBP1 | Q96B97 | SKP1 | P63208 |
| SLC11A2 | P49281 | SLC16A1 | P53985 | SLC25A10 | Q9UBX3 |
| SLC29A1 | Q99808 | SLC30A5 | Q8TAD4 | SLC31A1 | O15431 |
| SLC33A1 | O00400 | SLC3A2 | P08195 | SLC7A2 | P52569-1, P52569-2 |
| SLC7A5 | Q01650 | SMYD2 | P42785 | SMYD3 | Q9H7B4, Q9NRG4 |
| SNAP23 | O00161 | SNX18 | Q96RF0 | SPAST | Q9UBP0 |
| SPC24 | Q8NBT2 | SPC25 | Q15005 | SRC | P12931-1 |
| STC2 | O76061 | STEAP3 | Q658P3 | SUPV3L1 | Q8IYB8 |
| SURF1 | Q15526 | SWAP70 | Q9UH65 | SYTL1 | Q8IYJ3 |
| TACC3 | Q9Y6A5 | TACO1 | Q9BSH4 | TAF7 | Q15545, Q5H9L4 |
| TARBP2 | Q15633 | TARS2 | Q9BW92 | TBC1D30 | P48145 |
| TBP | P20226 | TBRG4 | Q969Z0 | TES | Q9NZU5 |
| TFB2M | Q9H5Q4 | TFDP1 | Q14186 | TGM1 | P22735 |
| TIMELESS | Q9UNS1 | TIMM23 | O14925 | TIMM50 | Q3ZCQ8 |
| TK1 | P04183 | TMEM126A | Q8IUX1 | TOP2A | P11388 |
| TOP3A | Q13472 | TOPBP1 | Q92547 | TPP1 | Q96AP0 |
| TPX2 | Q9ULW0 | TRAM1 | Q15629 | TRIM27 | P14373 |
| TRIM3 | O75382 | TRMT10C | Q7L0Y3 | TST | Q16762 |
| TUBGCP4 | Q9UGJ1 | TYMS | P04818 | UBE2C | O00762 |
| UBE2G1 | P62253 | UBE2S | Q16763 | UBE2T | Q9NPD8 |
| UCKL1 | Q9NWZ5 | UHRF1 | Q96T88 | UNG | P13051-1 |
| UQCRC1 | P31930 | UQCRC2 | P22695 | UQCRFS1 | P47985 |
| UQCRH | P07919 | USF2 | Q15853 | USP13 | Q92995 |
| VIMP | Q9BQE4 | WARS | P23381 | XPO6 | Q9NY97 |
| YARS | P54577 | YARS2 | Q9Y2Z4 | YME1L1 | Q96TA2 |
| ZFP91 | Q6PHW0 | ZFYVE16 | Q7Z3T8 | ZKSCAN1 | P17029 |
| ZNF24 | Q96GE5 | ZWILCH | Q9H900 | | |

| Input | Ensembl Id | Input | Ensembl Id | Input | Ensembl Id |
|---|---|---|---|---|---|
| AAMP | ENSG00000127837 | ACADM | ENSG00000117054 | ACADVL | ENSG00000072778 |
| ADD1 | ENSG00000087274 | AIFM2 | ENSG00000042286 | ANXA1 | ENSG00000135046 |
| ASNS | ENSG00000070669 | CAT | ENSG00000121691 | CCNA2 | ENSG00000145386 |
| CCNB1 | ENSG00000134057 | CCNB2 | ENSG00000157456 | CCND1 | ENSG00000110092 |
| CDC45 | ENSG00000093009 | CDK1 | ENSG00000170312 | CEBPB | ENSG00000172216 |
| DHFR | ENSG00000228716 | DLGAP5 | ENSG00000126787 | EZH2 | ENSG00000106462 |
| FANCD2 | ENSG00000144554, ENST00000419585 | FANCI | ENSG00000140525, ENST00000310775 | FDFT1 | ENSG00000079459 |

| Input | Ensembl Id | Input | Ensembl Id | Input | Ensembl Id |
|---|---|---|---|---|---|
| FHL2 | ENSG00000115641 | FN1 | ENSG00000115414 | FTH1 | ENST00000273550 |
| HMGCR | ENSG00000113161 | HMOX1 | ENSG00000100292 | HSPA5 | ENSG00000044574 |
| IDH1 | ENSG00000138413 | IFI30 | ENSG00000216490 | ITGA5 | ENSG00000161638 |
| JUNB | ENSG00000171223, ENST00000302754 | KPNA2 | ENST00000330459.7 | LGALS3 | ENSG00000131981 |
| MDC1 | ENSG00000137337, ENST00000376406 | MECP2 | ENST00000303391, ENST00000453960 | MLH1 | ENSG00000076242 |
| MRPL18 | ENSG00000112110 | MT-CO1 | ENST00000361624 | MT-CO2 | ENST00000361739 |
| MT-ND1 | ENST00000361390 | MT-ND4 | ENST00000361381 | MT-ND5 | ENST00000361567 |
| MUC1 | ENSG00000185499 | NDRG1 | ENSG00000104419 | NQO1 | ENSG00000181019 |
| PDCD4 | ENSG00000150593 | PLIN2 | ENSG00000147872 | PLK1 | ENSG00000166851 |
| POLA1 | ENSG00000101868 | POLRMT | ENSG00000099821 | PSAP | ENSG00000122852, ENSG00000185303 |
| PTPN9 | ENSG00000169410 | RNASEL | ENSG00000135828 | RRM2 | ENSG00000171848 |
| SESN2 | ENSG00000130766 | STEAP3 | ENSG00000115107 | TFB2M | ENSG00000162851 |
| TK1 | ENSG00000167900 | TOP2A | ENSG00000131747 | TPP1 | ENSG00000166340 |
| TRIM3 | ENSG00000110171 | TYMS | ENSG00000176890 | | |

#### Interactors (462)

| Input | UniProt Id | Interacts with | Input | UniProt Id | Interacts with |
|---|---|---|---|---|---|
| AAMDC | Q9H7C9 | P54252 | AAMP | C9JG97 | O14503 |
| AARS2 | Q5JTZ9 | Q8IWL3 | ABL2 | P42684 | P19174 |
| ACAD9 | Q9H845 | O75489, Q9NPL8 | ACADM | P11310 | P0DTC5 |
| ACADVL | P49748 | Q53T94 | ACO2 | Q99798 | Q8IWL3 |
| ACOT9 | Q9Y305 | P13569 | ACP2 | P11117 | Q13153 |
| ACSF3 | Q4G176 | P10809 | ACTBL2 | Q562R1 | Q8IW35 |
| ADD1 | P35611 | Q13153 | ADRBK1 | P25098 | Q13635 |
| AGA | P20933 | P07196 | AGO1 | Q9UL18 | P11940 |
| AHNAK | Q09666-2 | Q5SY16 | AK6 | Q9UIJ7 | P02649 |
| AKR1C1 | Q04828 | P26045 | ALCAM | Q13740 | P17931 |
| ANKRD52 | Q8NB46 | P53350 | ANKS1A | Q49AR9 | Q9NX63 |
| ANXA1 | P04083 | Q13546 | AQR | O60306 | P62306, P62304 |
| ARAP1 | Q96P48 | P62993 | ARHGEF11 | O15085 | P61586 |
| ARHGEF2 | Q92974 | O43524 | ARL6IP6 | Q8N6S5 | Q9NPL8 |
| ASF1B | Q9NVP2 | P49736 | ATAD2 | Q6PL18 | Q9Y6Q9 |
| ATG2A | Q2TAZ0 | Q9GZQ8 | ATP5J2 | P56134 | P01106 |
| ATPAF1 | Q5TC12 | P06576 | ATPAF2 | Q8N5M1 | P49247 |
| AURKA | O14965 | P53350 | AURKB | Q96GD4 | P06493 |
| AZGP1 | P25311 | P04156 | BCAS2 | O75934 | Q99757 |
| BCKDK | O14874 | P41208 | BMP7 | P18075 | Q5VWX1 |
| BOLA3 | Q53S33 | Q86SX6 | BRAT1 | Q6PJG6 | Q13315 |
| BRIP1 | Q9BX63 | P54278, P40692 | C11orf68 | Q9H3H3 | Q15056 |
| C14orf1 | Q9UKR5 | Q6IN84 | C1QBP | O35658 | Q00059 |
| C2orf47 | Q8WWC4 | P55056, P02654 | C2orf49 | Q9BVC5 | O76024 |
| C5orf22 | Q49AR2 | Q9Y2W2 | CAPN2 | P17655 | P07237 |
| CAT | P04040 | Q2T9J0 | CBS | P35520 | P06241 |
| CCNA2 | P20248 | Q13415, O75496, Q9H211 | CCNB1 | P14635 | Q99640 |
| CCNB2 | O95067 | Q99640 | CCND1 | P24385 | P06400 |
| CCNH | P51946 | P24941 | CD47 | Q08722-3 | Q9H2K0 |
| CD59 | P13987 | Q15363 | CDC20 | Q12834 | P24941 |

| Input | UniProt Id | Interacts with | Input | UniProt Id | Interacts with |
|---|---|---|---|---|---|
| CDC23 | Q8BGZ4 | Q9UKT4 | CDC45 | O75419 | Q14691 |
| CDCA5 | Q9CPY3, Q96FF9 | O60216 | CDCA8 | Q53HL2 | Q96GD4 |
| CDK1 | P06493 | P21796 | CDK2 | P24941 | O75496, Q9H211 |
| CDK7 | P50613 | P28715 | CEBPB | P17676 | Q13485 |
| CEP55 | Q53EZ4 | O75419 | CHAF1B | Q13112 | P62805 |
| CHCHD2 | Q9Y6H1 | P07237 | CHTOP | Q9Y3Y2-3, Q9Y3Y2 | Q5VWX1 |
| CLCN7 | P51798 | Q53GS7 | CLGN | O14967 | P04578 |
| CLIC3 | O95833 | Q9UN37 | CLK3 | P49761 | Q93008 |
| CLPP | Q16740 | P54252 | COL18A1 | P39060-PRO_0000005794 | P19338 |
| COMMD8 | Q9NX08 | Q14186, O75461 | COQ5 | Q5HYK3 | Q9NZJ6 |
| COQ6 | Q9Y2Z9 | Q9NZJ6, Q5HYK3 | CORO1A | P31146 | P40763 |
| COX15 | Q7KZN9 | P27824 | COX4I1 | P13073 | O75489 |
| CRABP1 | P29762 | P13569 | CST3 | P01034 | P05067 |
| DAP3 | P51398 | P48431 | DARS2 | Q6PI48 | Q8NFJ9 |
| DDX55 | Q8NHQ9 | Q9NRD5 | DEPTOR | Q8TB45 | Q13077 |
| DLGAP5 | Q15398 | Q14974 | DNMT1 | P26358 | Q13547 |
| DNPH1 | O43598 | O43598 | DPH2 | Q9BQC3 | O95363 |
| DSC1 | Q08554 | P13569 | DYNLL1 | P63167 | Q13627 |
| ECD | O95905 | Q9Y265 | ECH1 | Q13011 | P26641 |
| ECHS1 | P30084 | P40763 | ECSIT | Q9BQ95 | O75489, Q9NPL8, P03905 |
| EDRF1 | Q3B7T1 | O14832 | EPS15 | P42566 | P52594 |
| ERCC6L | Q2NKX8 | P53350 | EZH2 | Q15910 | P46100 |
| FAIM | Q9NVQ4 | O43597 | FAM114A1 | Q8IWE2 | Q9UBN6 |
| FAM129A | Q9BZQ8 | P31749 | FANCD2 | Q9BXW9 | P38398 |
| FANCI | Q9NVI1 | Q9Y5B0 | FASTKD2 | Q9NYY8 | P13569 |
| FBXL15 | Q9H469 | P63208, Q13616 | FBXO21 | O94952 | Q13616 |
| FDFT1 | P37268 | Q9H1C4 | FHL2 | Q14192 | P05556 |
| FN1 | P02751 | P05556 | FNBP1L | Q5T0N5 | Q9Y4D1 |
| FNDC3B | Q53EP0-3 | O14561 | FOXP4 | Q8IVH2 | Q9BZS1 |
| FRG1 | Q14331 | Q9HCG8 | FTH1 | P02794 | Q92574 |
| FUCA1 | P04066 | Q9BTT4 | FUNDC1 | Q8IVP5 | Q9GZQ8 |
| FXYD3 | Q14802-3 | Q9H2K0 | GABARAPL2 | P60520 | Q96A56 |
| GADD45GIP1 | Q8TAE8 | P40763 | GARS | P41250 | Q8WXH5 |
| GCDH | Q92947 | P29474 | GFM1 | Q96RP9 | P03508 |
| GGCX | P38435 | P0DTC5 | GINS2 | Q9Y248 | Q9BRT9, Q14691 |
| GINS4 | Q9BRT9 | Q9Y248, Q14691 | GNL2 | Q13823 | P0DTC9 |
| GOLGA1 | Q92805 | P18848 | GSN | P06396 | P24941, P38936 |
| GTF2H2C | Q6P1K8 | P50613 | GTF2H3 | Q13889 | P50613 |
| GTPBP10 | A4D1E9 | Q08379 | HADH | P40939 | P04233 |
| HADHA | P40939 | P04233 | HADHB | P55084 | O15530 |
| HDDC2 | Q7Z4H3 | P01189 | HDHD1 | Q08623 | Q8IVS8 |
| HEATR3 | Q7Z4Q2 | P0DTC7 | HELLS | Q9NRZ9 | P01106 |
| HERC2 | O95714 | P23025 | HMGCR | P04035 | Q9Y5Z9 |
| HMGXB4 | Q9UGU5 | O75928-2 | HMOX1 | P09601 | Q9NUX5 |
| HSD17B10 | Q99714 | P13639 | HSP90AB2P | Q58FF8 | P13569 |
| HSPA5 | P11021 | P14625 | ICMT | O60725 | Q3SXY8 |
| ICT1 | Q14197 | Q00059 | IDH1 | O75874 | P0DP23 |
| IFI30 | P13284 | P22735 | IQGAP3 | Q86VI3 | Q92845 |
| ITGA5 | P08648 | P05556 | JAK1 | P23458 | P40763 |

| Input | UniProt Id | Interacts with | Input | UniProt Id | Interacts with |
|---|---|---|---|---|---|
| JMJD6 | Q6NYC1 | P50750 | JMY | Q9QXM1 | Q09472 |
| JUNB | P17275 | P18848 | KANK2 | Q63ZY3 | P06730 |
| KAT8 | Q9H7Z6 | P04637 | KCTD5 | Q9NXV2 | Q9UBU9 |
| KDM3A | Q9Y4C1 | O75582 | KDM5C | P41229 | Q13127 |
| KIAA0101 | Q15004 | P12004 | KIAA1324 | Q6UXG2-3 | P07196 |
| KIAA1524 | Q8TCG1 | P01106 | KIF1A | Q12756 | O76024 |
| KIF23 | Q02241-2, Q02241 | Q9H0H5 | KIFAP3 | Q92845 | Q9Y496 |
| KLC4 | Q9NSK0 | P31946 | KPNA2 | P52292 | O96017 |
| LDLR | P01130 | Q9H2K0 | LGALS3 | P17931 | P17931 |
| LGALS3BP | Q08380 | Q9BZR8 | LIG1 | Q96JA1 | P22681 |
| LIMA1 | Q9UHB6 | P60953 | LOXL2 | Q9Y4K0 | P02751 |
| LRP10 | O75096 | Q9BQB4 | LXN | Q9BS40 | Q9UQK1 |
| LYRM7 | Q5U5X0 | O14561 | MAD2L1 | Q13257, Q13257-1, Q9Z1B5 | O60566, Q13257, Q12834, Q9Y6D9 |
| MADD | Q8WXG6 | Q9NUX5 | MAK16 | Q9BXY0 | Q9HAP6 |
| MAP1B | P46821 | P10636-8 | MAP7D1 | Q3KQU3 | P01106 |
| MARCKS | P29966 | P13569 | MCCC2 | Q9HCC0 | P13569 |
| MCM2 | P49736 | O75419 | MCM3 | P25205 | P33992, P49736 |
| MDC1 | Q14676 | Q13315 | MECP2 | P51608 | P10599 |
| MGA | Q8IWI9 | Q13547 | MGME1 | Q9BQP7 | Q9BQ95 |
| MIA3 | Q5JRA6 | Q15436 | MICU1 | Q9BPX6 | Q9H4I9, Q9BPX6, Q8NE86 |
| MICU2 | Q8IYU8 | Q9H4I9, Q9BPX6, Q8NE86 | MIOS | Q9NXC5 | P55735 |
| MIS18A | Q9NYP9 | P53350 | MKI67 | P46013 | Q9NS87 |
| MLH1 | P40692 | P54278 | MMADHC | Q9H3L0 | Q9P0N9 |
| MORF4L1 | Q9UBU8-2, Q9UBU8 | Q86YC2 | MRFAP1 | Q9Y605 | Q9H2K0 |
| MRPL11 | Q9Y3B7 | Q13371 | MRPL12 | P52815 | P21673 |
| MRPL13 | Q9BYD1 | P42858 | MRPL15 | P49406 | P67809 |
| MRPL17 | Q9NRX2 | P52597 | MRPL19 | P49406 | P67809 |
| MRPL2 | Q9BZE1 | P10809 | MRPL23 | Q16540 | O14561 |
| MRPL27 | Q8IXM3 | Q969Q1 | MRPL28 | Q13084 | O60437 |
| MRPL32 | Q9BYC8 | Q9GZV8 | MRPL37 | Q9BZE1 | P10809 |
| MRPL38 | Q96DV4 | Q96LA8 | MRPL40 | Q9NQ50 | P14373 |
| MRPL41 | Q8IXM3 | Q969Q1 | MRPL42 | Q9Y6G3 | P04618 |
| MRPL44 | Q9H9J2 | Q16891 | MRPL45 | Q9BRJ2 | Q6NZI2 |
| MRPL46 | Q9H2W6 | O14561 | MRPL47 | Q9HD33 | P10809 |
| MRPL49 | Q13405 | P16284 | MRPL53 | Q96EL3 | Q15370 |
| MRPL9 | Q9BYD2 | P21673 | MRPS14 | O60783 | P06396 |
| MRPS17 | Q9Y2R5 | Q92845 | MRPS18B | Q9Y676 | P06400 |
| MRPS2 | Q9Y399 | P03372 | MRPS22 | P82650 | P48431 |
| MRPS23 | Q9Y3D9 | O15160 | MRPS25 | P82663 | Q92845 |
| MRPS26 | Q9BYN8 | P48431 | MRPS27 | Q92552 | P03372 |
| MRPS28 | P82673 | P03372 | MRPS30 | Q9NP92 | P10809 |
| MRPS31 | Q92665 | P40763 | MRPS35 | P82673 | P03372 |
| MRPS9 | P82933 | P48431 | MRRF | Q96E11 | P14373 |
| MSMO1 | Q15800 | P09601 | MT-ND1 | P03886 | O75489 |
| MT-ND4 | P03905 | O75380 | MT-ND5 | P03915 | O75489 |
| MTERF3 | Q96E29 | P22732 | MTERF4 | Q7Z6M4 | Q6ZPD8 |
| MTG1 | Q9BT17 | Q9NRD5 | MTHFD2 | P13995 | Q5S007 |
| MTIF3 | Q9H2K0 | P01130 | MTPAP | Q9NVV4 | Q9UHD2 |

| Input | UniProt Id | Interacts with | Input | UniProt Id | Interacts with |
|---|---|---|---|---|---|
| MUC1 | P15941-PRO_0000317447 | P17676 | MUT | P22033 | P42858 |
| NARS2 | Q96I59 | P10809 | NBR1 | Q14596 | P49841 |
| NCAPD3 | P42695 | P05919 | NCOA4 | Q13772 | O95714 |
| NDOR1 | Q9UHB4 | Q9UBE8 | NDRG1 | Q92597 | Q9BYZ2 |
| NDUFA12 | Q9UI09 | O75380, O75489 | NDUFA13 | Q9P0J0 | Q9UI09, O75489, Q9P032, O43181 |
| NDUFA2 | O43678 | O75380, Q9UI09, O75489, P49821, O43920, O43181 | NDUFA3 | O95167 | O75489 |
| NDUFA4 | O00483 | Q9UI09 | NDUFA5 | Q16718 | O75489 |
| NDUFA6 | P56556 | O75380, Q9UI09, O75489, O43920 | NDUFA9 | Q16795 | P19404, O75380, O75489 |
| NDUFAB1 | O14561 | P26441 | NDUFAF1 | Q9Y375 | O75489, Q9NPL8, O43920 |
| NDUFAF2 | Q8N183 | O75396 | NDUFAF4 | Q9P032 | O75489, Q9NPL8, O43920, Q9BU61 |
| NDUFB10 | O96000 | O75489 | NDUFB11 | Q9NX14 | O75489, Q9NPL8 |
| NDUFB4 | O95168 | O75489 | NDUFB6 | O95139 | O75489 |
| NDUFB9 | Q9Y6M9 | O75489 | NDUFS1 | P28331 | O75380, O75489 |
| NDUFS2 | O75306 | P19404, O75380, O75489, Q9BU61 | NDUFS3 | O75489 | P03886, O95139, Q9UI09, P03923, Q9P032, P03915, O43920, Q9BU61, P19404, O75380, P49821, P56181, O43181 |
| NDUFS4 | O43181 | O75489 | NDUFS5 | O43920 | O75380, O75489, Q9P032 |
| NDUFS7 | O75251 | Q9UI09, O75489, Q9P032, Q5TEU4 | NDUFS8 | O00217 | O75380, O75489 |
| NDUFV1 | P49821 | O75489, P56181 | NDUFV2 | P19404 | O75380, O75489 |
| NFATC2IP | Q8NCF5-2 | O96015 | NFS1 | Q9Y697 | Q9NZJ6 |
| NHP2L1 | P55769 | P19320 | NQO1 | P15559 | P15559 |
| NR2F6 | P10588 | P83916 | OAT | P04181 | P05067 |
| OGDH | Q02218 | P42858 | OGFOD1 | Q8N543 | P62266 |
| OGFOD2 | Q6N063-2 | P55212 | ORC3 | Q9UBD5 | Q13415 |
| OXA1L | Q2M1J6 | Q92993 | PBK | Q96KB5 | P04637 |
| PBXIP1 | Q96AQ6 | Q9UER7 | PC | P11498 | P48163 |
| PDCD2 | Q16342 | Q92993 | PDCD4 | Q53EL6 | P02649 |
| PDF | Q99988 | Q9Y6A5 | PDHA1 | P08559 | P42229 |
| PDHB | P75391 | P00747 | PDK3 | Q15120 | Q15119 |
| PDP1 | Q8IY26 | Q14627 | PDZD11 | Q5EBL8 | Q04656 |
| PDZK1 | Q5T2W1 | P19404 | PEG10 | Q86TG7-2 | Q9NQT4 |
| PEX3 | P56589 | Q9Y5Y5, P40855 | PFDN5 | Q99471 | Q13547 |
| PHGDH | O43175 | P03372 | PIH1D1 | Q9NWS0 | Q9Y265, Q9Y230 |
| PKMYT1 | Q99640 | P61024, P06493 | PKP1 | Q13835-2 | P08670 |
| PKP4 | Q99569 | Q96RT1 | PLIN2 | Q99541 | Q8WTS1 |
| PLK1 | P53350 | P23588 | PLOD1 | Q02809 | P55317 |
| PLSCR1 | O15162 | Q15077 | PNPLA2 | Q96AD5 | P00533 |
| PNPT1 | Q8TCS8 | Q8IWL3 | POLA2 | Q14181 | P31948 |
| POLD1 | P28340 | P12004 | POLD2 | P49005 | P12004 |
| POLDIP2 | Q9Y2S7 | P13196 | POLE | Q07864 | P01106 |
| POLE3 | Q9NRF9 | Q01658 | POLG | Q9WMX2 | O14980 |
| POLR3A | O14802 | P19388 | POLRMT | O00411 | Q00059 |
| PPP1R9B | Q96SB3 | P38398 | PPP2R1B | P30154 | Q9BRV8 |
| PPP2R5C | Q13362-3, Q13362-2, Q13362-1 | O96017 | PRIM1 | P49642 | P09884 |

| Input | UniProt Id | Interacts with | Input | UniProt Id | Interacts with |
|---|---|---|---|---|---|
| PRKACG | P22612 | P05067 | PRMT7 | Q582G4 | P62805 |
| PRPS1 | P60891 | Q86WV5 | PSAP | Q8IWL2 | Q9UGM3 |
| PTCD3 | Q96EY7 | P67809 | PTPMT1 | Q8WUK0 | P14373 |
| PTPN9 | P43378 | Q9NZJ6 | PTPRE | P23469 | P62993 |
| PYCR1 | P32322 | Q99497 | QIL1 | Q5XKP0 | Q96CM8 |
| QRICH1 | Q2TAL8 | Q14938 | RAB2B | Q8WUD1 | Q96FJ0 |
| RAB4B | P61018 | O75419 | RACGAP1 | Q9H0H5 | P53350 |
| RAD51C | O43502 | Q06609 | RAF1 | P04049 | P43246 |
| RB1 | P06400 | P06400 | RBM45 | Q8IUH3 | Q14974 |
| RBP1 | P09455 | Q9UBN6 | RCN1 | Q15293 | Q969Y2 |
| REEP6 | Q96HR9-2, Q96HR9 | Q99757 | RFC1 | P35251 | P35250, P40938, P40937, P35249 |
| RFC5 | P40937 | P35249 | RFX5 | P48382 | Q92769 |
| RIF1 | Q5T3J3 | P45973 | RMI2 | Q96E14 | Q14159 |
| RNASEL | Q05823 | P46940 | RNF14 | Q9UBS8 | P20823 |
| RNMTL1 | Q9HC36 | Q9HC36 | RPL26L1 | Q9UNX3 | P0DTC9 |
| RPLP1 | P05386 | Q9GZQ8 | RPS27L | Q71UM5 | Q00987 |
| RRM1 | P23921 | P31350 | RRM2 | P31350 | Q9UM11 |
| RTN4IP1 | Q8WWV3 | Q9HAU4 | S100P | P25815 | P35579 |
| SAPCD2 | Q86UD0 | O75496 | SDHA | P31040 | P26045 |
| SDHB | P21912 | Q8IWL3 | SDHC | Q99643 | Q92993 |
| SEMA4C | Q9C0C4 | Q96PM5 | SEMG1 | P04279-2 | Q96JC9 |
| SESN2 | P58004 | P62877 | SF3B5 | Q9BWJ5 | Q96FJ0 |
| SFXN2 | Q96NB2 | Q7Z5P4 | SGSH | P51688 | Q8NBK3 |
| SH3KBP1 | Q96B97 | Q99962 | SHCBP1 | Q8NEM2, Q9Z179 | Q9H0H5 |
| SKP1 | P63208 | P46527, P61024, P24941, P38936, Q9UKT4 | SLAIN2 | Q9P270 | Q9Y6A5 |
| SLC25A10 | Q9UBX3 | P13569 | SLC31A1 | O15431 | O15431 |
| SLC33A1 | O00400 | P13569 | SLC3A2 | P08195-4 | Q9NX09 |
| SLMAP | Q14BN4 | Q96CV9 | SLMO2 | Q9Y3B1 | O43715 |
| SMYD2 | Q9NRG4 | Q15185 | SMYD3 | Q9H7B4 | Q16512 |
| SNAP23 | O00161 | Q15836 | SNX18 | Q96RF0 | O14672 |
| SPC25 | Q15005 | P0DTC3 | SRC | P00523 | Q9Y4D1 |
| STC2 | O76061 | P0DTC8 | SWAP70 | Q9UH65 | Q8WZA2 |
| SYPL1 | Q16563 | Q9BW92 | TACC1 | O75410 | Q96GD4 |
| TACC2 | B2RWP4 | P22735 | TACC3 | Q9Y6A5 | Q00994 |
| TACO1 | Q9BSH4 | P21673 | TAF7 | Q15545 | P20226 |
| TARBP2 | Q15633 | Q96T60 | TARS2 | Q9BW92 | O60664 |
| TBC1D30 | Q9Y2I9-2 | Q9UGI0 | TBP | P20226 | Q00403 |
| TBRG4 | Q969Z0 | Q9UBN6 | TCAF1 | Q9Y4C2-2 | Q9Y6D9 |
| TELO2 | Q9Y4R8 | Q13315 | TES | Q9UGI8 | P13569 |
| TFB2M | Q9H5Q4 | O00411 | TFDP1 | Q14186 | O75496 |
| TGM1 | P22735 | Q9Y5Y2 | TGM2 | P21980-2 | Q5T160 |
| TIMM23 | O14925 | Q7Z5P4 | TIMM50 | Q3ZCQ8 | P04049 |
| TK1 | P04183 | P04183 | TMED1 | Q13445 | O60645, Q9NV70 |
| TMEM106B | Q9NUM4 | P42858 | TONSL | Q96HA7 | Q08945 |
| TOP2A | P11388 | P38398 | TOP3A | Q13472 | Q14159 |
| TOPBP1 | Q92547 | O75419 | TPP1 | P49638 | Q6IN84 |
| TPT1 | P13693 | P29692 | TPX2 | Q9ULW0 | O75330 |
| TRAM1 | Q9Y6Q9 | Q09472 | TRIM27 | P14373 | Q96E11 |

| Input | UniProt Id | Interacts with | Input | UniProt Id | Interacts with |
|---|---|---|---|---|---|
| TRIM3 | O75382 | P49674 | TTC19 | Q6DKK2 | P09455 |
| TUBGCP4 | Q9UGJ1 | Q8TES7 | TYMS | P04818 | P19320 |
| UBE2C | Q5TZN3, O00762 | Q13164 | UBE2G1 | P62253 | P06307 |
| UBE2S | Q16763 | P13569 | UHRF1 | Q96T88 | P26358 |
| UNG | P13051-2 | P16671 | UQCRC1 | P31930 | P26641 |
| UQCRC2 | P22695 | Q13573 | UQCRFS1 | P47985 | Q8IWL3 |
| USF2 | Q15853 | Q15562 | USP13 | Q92995 | P54252 |
| VEZF1 | Q14119 | Q96KQ7 | VIMP | Q9BQE4 | Q6Q788 |
| WBSCR16 | Q96I51 | Q92993 | WDHD1 | O75717 | Q92830 |
| XPNPEP3 | Q9NQH7 | P08670 | YARS | P54577 | O75925 |
| YARS2 | Q9Y2Z4 | Q12834 | ZFP91 | Q96JP5 | P04608 |
| ZFPL1 | O95159 | Q9HC36 | ZFYVE16 | Q7Z3T8 | P62136 |
| ZKSCAN1 | P17029 | Q96CV9 | ZNF24 | P17028 | P02649 |
| ZNF330 | Q9Y3S2 | Q9UMX1 | ZNF579 | Q8NAF0 | P04608 |

| Input | ChEBI Id | Interacts with | Input | ChEBI Id | Interacts with |
|---|---|---|---|---|---|
| HADH | P40939 | 28494 | HADHA | P40939 | 28494 |
| IDH1 | O75874 | 16810 | MMADHC | Q9H3L0, Q99LS1 | 16856 |
| MUT | P22033 | 18408 | SRC | P12931 | 15532 |

### 7. Identifiers not found

These 59 identifiers were not found neither mapped to any entity in Reactome.

| | | | | | | | |
|---|---|---|---|---|---|---|---|
| ARHGAP36 | BCAP29 | BCL7B | C16orf13 | C8orf82 | CECR5 | CKAP2 | CLIC4 |
| CLPX | DHX40 | DNLZ | DTD2 | EIF1B | ENDOG | ENOSF1 | FAM160B2 |
| FAM174B | FLYWCH2 | FOXRED1 | GPR107 | HIGD2A | HMCES | HMGN4 | HSDL2 |
| ICA1 | IFRD1 | ISOC2 | KIAA0513 | KTI12 | LACTB | LRRC58 | METTL9 |
| MIPEP | MYCBP | NCAM2 | NCBP2-AS2 | NOL8 | NT5DC2 | NT5DC3 | NUSAP1 |
| PHLDA3 | PREPL | PTER | R3HDM4 | RPUSD2 | RSRC1 | SBNO1 | SBSN |
| SDHAF2 | SELH | TAMM41 | THYN1 | TMEM64 | TMEM70 | TTC39B | UBL3 |
| UQCC1 | VPS13C | VWA8 | | | | | |
